# Bacterial DNA invasion triggers transposable element proliferation and genome expansion

**DOI:** 10.64898/2026.07.14.738529

**Authors:** Zachary P. Cohen, Lindsey Perkin, Paul B. Frandsen, Michael DeGiorgio, Raquel Assis

## Abstract

Genome size variation in eukaryotes is driven largely by transposable elements (TEs), yet the biological mechanisms that initiate their proliferation remain understudied. Here, we identify a recurrent association between bacterial horizontal gene transfer (HGT) and bursts of TE activity that contribute to genome expansion. By leveraging comparative genomics and genus-level pangenome analyses across three species of the nut weevil, *Curculio*, we detect extensive bacterially derived DNA sequences embedded within structurally dynamic genomic regions. These HGT-associated regions are dominated by a small number of young, proliferating TE families, particularly DNA type II Mavericks, which encapsulate transferred bacterial sequences and comprise a substantial fraction of recent genomic DNA in derived lineages. Analyses of codon usage bias, intron length, and functional enrichment suggest that most transferred genes undergo progressive pseudogenization over evolutionary time, whereas a subset of selectively advantageous HGTs persist. Together, our findings support a model linking foreign DNA invasion with TE proliferation, genome size variation, and molecular innovation.

## Introduction

Eukaryotic genomes vary substantially in size and organization, even within individuals of the same species^1–4^. Despite this variation, most eukaryotic genomes encode broadly similar numbers of genes, typically on the order of tens of thousands^5^. This disconnect between gene number and genome size, referred to as the C-value paradox, has motivated decades of research into the mechanisms shaping genome size, structure, and function^2,6–10^. While aneuploidy and polyploidy account for some of the observed variation in genome size and gene content, and occasionally lead to speciation^11–13^, TE proliferation remains the predominant explanation for genome expansion across plants and animals^14–20^. Although abiotic stress, selection, and demography have been implicated in TE activity, the biological mechanisms triggering TE proliferation remain unclear^13,21–28^.

One potential source of TE proliferation and genome size variability is horizontal gene transfer (HGT), through which genetic material is acquired from distantly related organisms. Beyond the ancient endosymbiotic events that gave rise to mitochondria and plastids^29,30^, HGT from symbiotic microbes continues to introduce foreign DNA into host eukaryotic genomes^31^. HGT is a well-documented evolutionary force in prokaryotes^32^, yet its influence on eukaryotic genome evolution remains less clear. While HGT can have profound phenotypic^33^ and fitness effects^34^ at the individual level, evolutionary persistence requires integration into the germline and subsequent transmission to offspring, a process thought to be rare in multicellular eukaryotes^35^. Yet, several classes of TEs are implicated in the movement of DNA across evolutionarily distinct lineages^36^. Notably, the DNA Type II Mavericks, large (∼15-20 kb) virus-like DNA transposons^37^, have been observed to capture and mobilize foreign DNA between distantly related species^38^. Together, these observations raise the possibility that interactions between foreign DNA integration and TEs could contribute to genome restructuring, though the evolutionary significance of these processes remains unknown.

Insects provide a particularly informative system for investigating the evolutionary consequences of HGTs because transfers from endosymbiotic bacteria are ubiquitous across this cosmopolitan clade^39,40^ and contribute to adaptation in many lineages^41–47^. The nut weevil, genus *Curculio*, in which endosymbiotic bacteria have been documented^48^, offers a unique opportunity to examine this phenomenon and its genetic consequences in an evolutionary framework. Despite sharing similar ecological niches, life histories, and karyotypes (2n = 24A + Xy*_p_*), *Curculio* species exhibit striking differences in genome size and HGT burden. These characteristics provide a natural comparative system for investigating how bacterial DNA integration shaped genome restructuring and TE proliferation throughout evolutionary history.

Here, we study the relationship among HGT, TE dynamics, and genome evolution across three *Curculio* species: the pecan weevil (*C. caryae*; 2.2 Gb) and the western acorn weevil (*C. nanulus*; 1.5 Gb), which are both endemic to North America, as well as the more distantly related European acorn weevil (*C. glandium*; 1.1 Gb). By integrating comparative pangenomic, molecular, and phylogenomic analyses, we uncover a relationship among foreign DNA integration, TE proliferation, and genome size variation. Furthermore, we find specific TE families linked to the amplification and domestication of bacterially transferred genes. These findings are consistent with prior observations that extensive TE proliferation can be tolerated over evolutionary timescales^49^ and reveal previously underappreciated links among symbiosis, genome architecture, and evolution.

## Results

### Comparative genomics reveals extensive HGT in *Curculio*

To characterize the evolutionary history of HGTs across the three *Curculio* species, we reconstructed a time-calibrated phylogeny using 2,542 orthologous genes (see *Methods*). We rooted this tree using the distantly related Colorado potato beetle (*Chrysomelidae Leptinotarsa decemlineata*) and estimated the divergence of *C. caryae* and *C. nanulus* from *C. glandium* at approximately 28 million years ago (MYA), and from each other at approximately seven MYA (Fig. 1). The topology and timing of these events are consistent with prior phylogenetic reconstructions of this group^50,51^.

**Figure 1.**
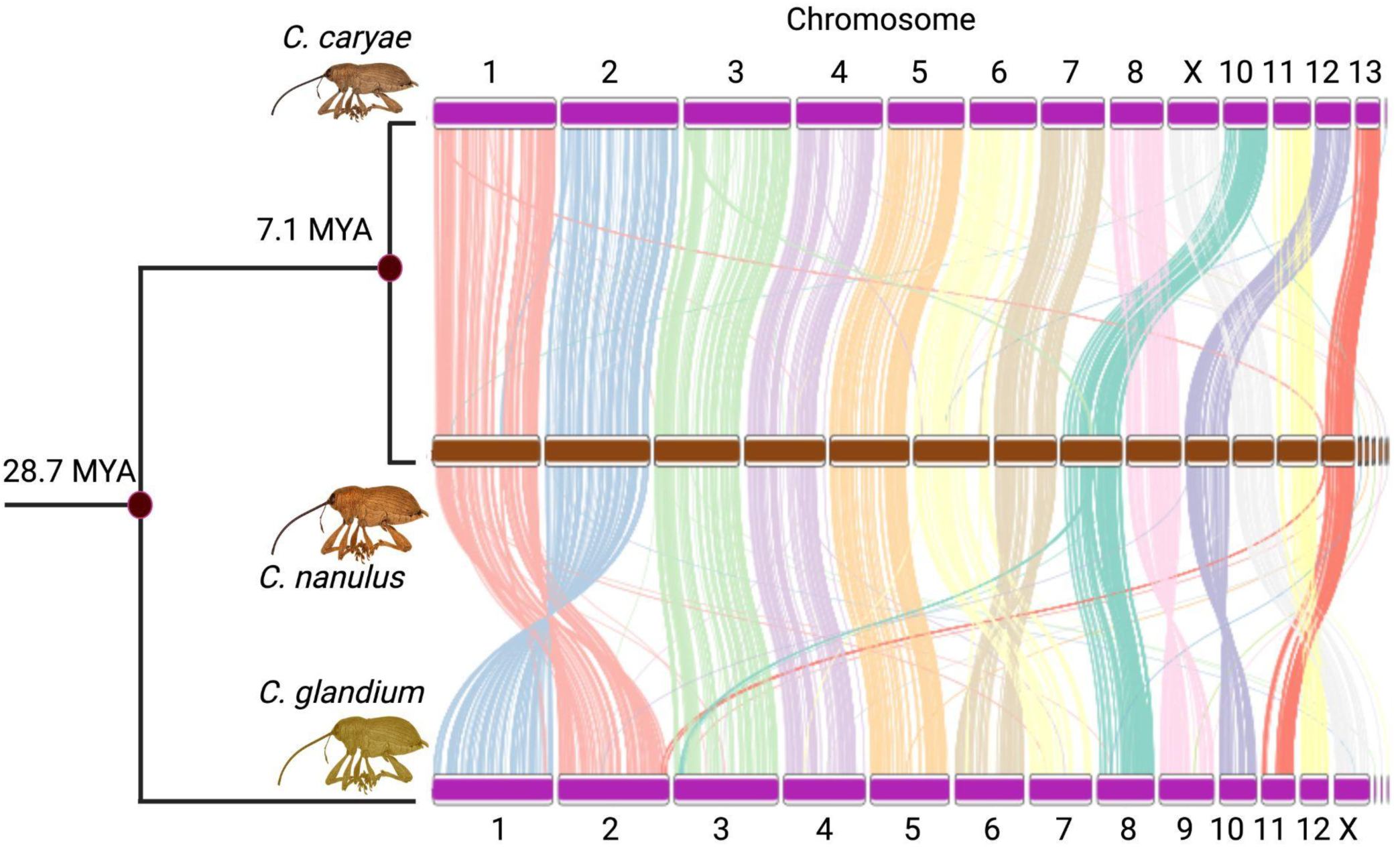
Time-calibrated phylogeny of sampled *Curculio* species with adjacent chromosomal ideographs illustrating 2,542 shared orthologous genes.

BLAST searches against non-repeat prokaryotic databases revealed hundreds of bacterially derived matches in each *Curculio* species (Supplementary Table 1). In *C. caryae*, we identified 132 species-specific chromosomal HGT matches to the known *Curculio* symbiont *Candidatus Curculioniphilus buchneri* (*Ccb*)^48^, a γ-proteobacterium originally isolated from *C. glandium*, and 68 matches to an extra chromosomal contig. Conversely, *C. glandium* and *C. nanulus* show no evidence of HGT from this symbiont.

The *Ccb*-derived regions in *C. caryae* encompass one contiguous tract on Chromosome 1 (∼500 kb), two tracts on Chromosome 3 separated by approximately 300 kb (∼335 kb and ∼220 kb), and an extrachromosomal contig (∼500kb; Fig. 2). Although the upstream and downstream Chromosome 3 regions do not share any genes, the *Ccb*-derived genes in each region map to overlapping sections of the extant *Ccb* reference genome, suggesting they may represent independent insertion events with complementary gene retention. Based on contiguity and gene content, the Chromosome 1 region contains the largest block of transferred genes (83 genes), followed by the extrachromosomal contig (61 genes), the upstream Chromosome 3 region (47 genes), and the downstream Chromosome 3 region (23 genes). No genes are shared among all four *Ccb*-derived regions. However, the Chromosome 1 region and extrachromosomal contig share the greatest number of genes (39 genes), whereas overlaps between the Chromosome 1 region and the two Chromosome 3 regions are more modest (Supplementary Table 2).

**Figure 2.**
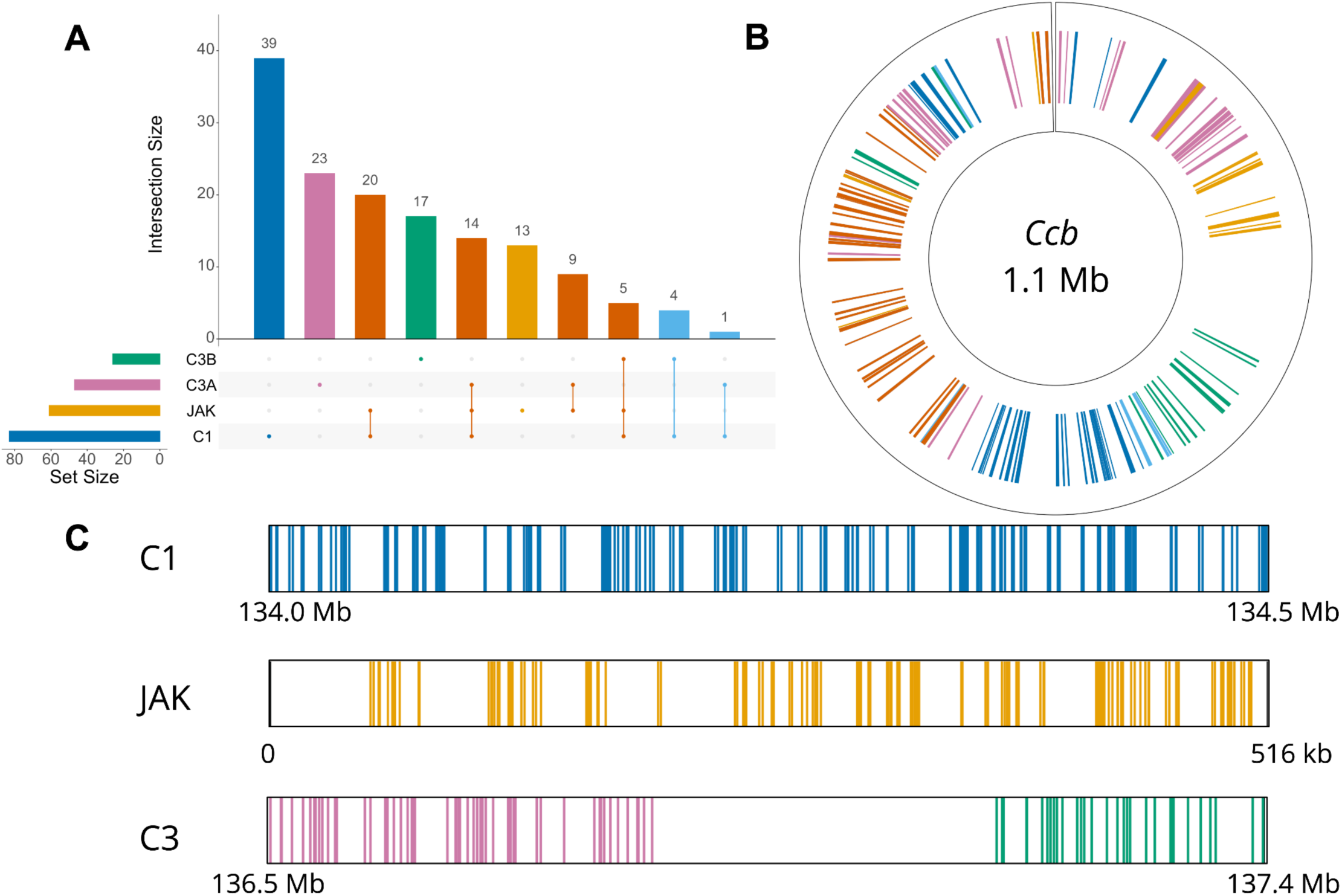
Distribution of *Ccb*-derived genes among the four HGT regions in *C. caryae:* the Chromosome 1 region (C1), the extrachromosomal contig JAKZMK010000236 (JAK), the upstream Chromosome 3 region (C3A), and the downstream Chromosome 3 region (C3B). (A) UpSet plot showing the number of shared and unique genes among the four regions. (B) Genomic coordinates of *Ccb*-derived genes from the four *C. caryae* regions mapped to the *Ccb* reference genome. (C) Genomic coordinates of *Ccb*-derived genes mapped to the four HGT regions in *C. caryae*. Colors indicate the corresponding region combinations: dark blue, C1; pink, C3A; green, C3B; yellow, JAK; light blue, combinations involving C1 with either C3A or C3B; and orange, combinations involving C1, C3A, C3B, and JAK.

Pairwise genetic distances across shared *Ccb* regions revealed that the extrachromosomal contig is least divergent from the Chromosome 1 region (0.16%), followed by the upstream (1.4%) and downstream (2.5%) Chromosome 3 regions (Supplementary Table 3). Among the chromosomal regions, the upstream and downstream Chromosome 3 regions differ from the Chromosome 1 region by 2.8% and 5.6%, respectively, and from each other by 11.0%. In contrast, the extrachromosomal contig and all three chromosomal regions are similarly divergent from the *Ccb* reference genome (21.4-22.9%).

Beyond the *Ccb*-derived insertions, we identified an additional 529 bacterially derived HGT annotations in *C. caryae*, 500 of which are attributed to the *Rickettsia* symbiont originally described from the cabbage seedpod weevil (*Ceutorhynchus assimilis*). In the two acorn-feeding species (*C. nanulus* and *C. glandium*), we detected 283 and 137 HGTs, respectively, with the majority also attributed to the same *Rickettsia* species (275 in *C. nanulus* and 135 in *C. glandium*). Thus, while *Ccb*-derived transfers appear restricted to *C. caryae*, *Rickettsia*-derived HGTs are shared across all three species and constitute the majority of detected HGTs.

### HGTs localize to TE-rich regions of significant structural variation

To investigate the relationship between HGTs and chromosomal architecture at the genus level, we generated pangenome graphs for the 13 homologous chromosomes based on the synteny of shared orthologous genes (Fig. 1; see *Methods*). At this scale, the distribution of structural variation is highly similar between *C. caryae* and *C. nanulus* (Spearman’s ρ = 0.97; *p* = 2.2 × 10^−60^), but differs markedly in *C. glandium* relative to both *C. caryae* (ρ = 0.45; *p* = 2.06 × 10^−53^) and *C. nanulus* (ρ = 0.46, *p* = 3.16 × 10^−56^, see *Methods*).

Approximately 20% of HGT regions overlap with TE-rich regions above the 90th percentile of graph-defined structural density (Fig. 3). Additionally, nearly all HGTs, regardless of node density, coincide with annotated TEs (Supplementary Figs. 1-39). These observations indicate that HGTs are nonrandomly associated with TE-rich genomic regions. Kimura divergence^52^ estimates (*D*) further revealed that structurally dense regions are dominated by young (*D* < 0.05) TEs in both *C. caryae* (85%) and *C. nanulus* (75%; Supplementary Fig. 40; Supplementary Tables 4 and 5; see *Methods*), linking regions of elevated structural variation with recent TE activity.

**Figure 3.**
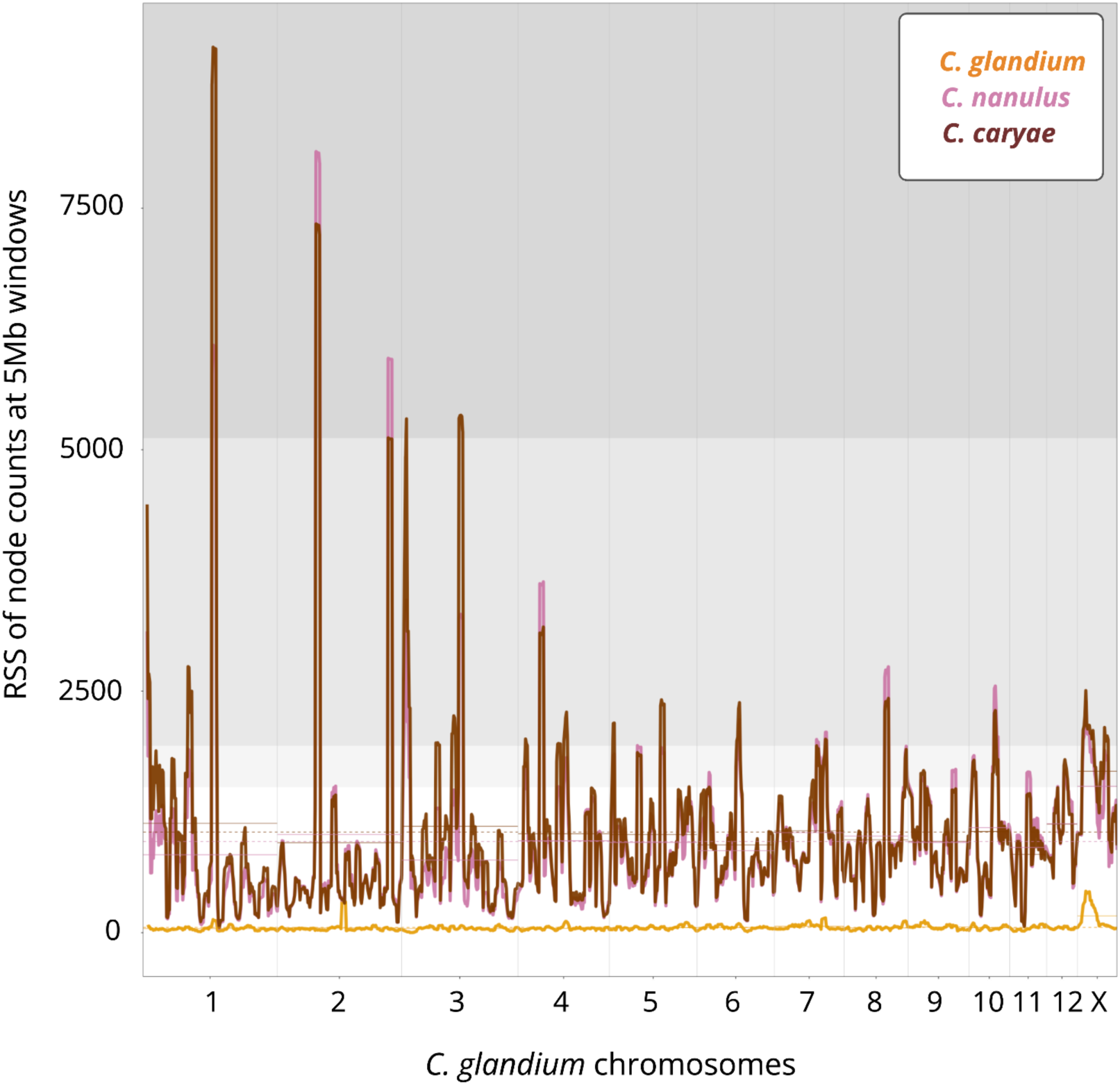
Root sum of squares (RSS) rendering of nodes isolated from the pangenome graph for each *Curculio* species, relative to *C. glandium* genomic coordinates. Shaded gray background bands represent the 90th, 95th, and 99th percentiles of node density (light to dark). Elevated peaks of species-specific structural variants occur at evolutionarily homologous TE “proliferation springs.” Horizontal dotted lines indicate the chromosomal mean RSS for each species.

### Mavericks dominate HGT regions and contribute to genome expansion

To test whether TE accumulation around HGTs reflects nonrandom genomic association, we first performed region-specific permutation tests comparing TE density within HGT cores and flanking regions to size-matched genomic backgrounds (see *Methods)*. We then extended this analysis using a multinomial regression framework to identify the genomic predictors that distinguish TE-associated HGTs, TE-independent HGTs, and flanking regions from chromosome-matched background regions (see *Methods*).

The large *Ccb* insertions on Chromosomes 1 and 3 of *C. caryae* generally occur in TE-depauperate cores (<2.5th percentile) flanked by TE-rich regions (>97.5th percentile). While this pattern is pronounced on both flanks of the Chromosome 1 HGT region (Supplementary Fig. 1), only the upstream flank of the Chromosome 3 insertion is TE-rich (Supplementary Fig. 3). Permutation tests showed that the observed flank-minus-core TE density contrasts for both HGT regions are significantly greater than expected under random placement (*p* < 10^−4^, see *Methods*). Additionally, the three contiguous *Ccb* insertions in *C. caryae* overlap significantly fewer TEs than expected relative to a size-controlled permuted genomic background (*p* = 9.9 × 10^−4^, see *Methods*). In contrast, *Rickettsia*-derived HGT windows span a broader range of TE densities across all three species.

Multinomial regression consistently identified coding sequence density and TE density as the strongest predictors of HGT (Supplementary Figs. 41-44). Relative to background genomic regions, TE-independent HGT windows exhibit significantly elevated coding sequence density (*p* = 3.76 × 10^−5^), whereas TE-associated HGT windows (*p =* 1.27 × 10^−4^) and flanking regions (*p* = 7.68 × 10^−12^) show significantly reduced coding density (Supplementary Figs. 41 and 42; see *Methods*). Specifically, a one standard deviation increase in coding sequence density raises the odds of TE-independent HGT windows by approximately 55%, while reducing the odds of TE-associated HGT windows by approximately 16% and flanking regions by approximately 23% (Supplementary Fig. 42; see *Methods*).

Across both young and old TE-associated HGT classes, increasing TE density is linked to greater odds of HGT occurrence relative to the genomic background. This corresponded to odds increases of approximately 220% (*p* = 3.28 × 10^−7^) for old TEs and 893% increase for young TEs (*p* = 4.18 × 10^−7^, see *Methods*; Supplementary Figs. 43 and 44). In contrast, GC content and GC skew exhibit comparatively weak and inconsistent effects. Together, these results indicate that HGT regions occupy genomic environments distinct from the broader genome and identify TE density as the strongest predictor of HGT occurrence.

The age of TEs within HGT regions is also highly correlated. The Chromosome 1 *Ccb* insertion spans a region containing six TEs, two of which exhibit high Kimura divergence (*D* > 0.50) and therefore cannot be reliably classified. The youngest identifiable element, a Maverick (*D* = 0.20 × 10^−3^; ∼0.33 MYA), establishes a lower bound for the insertion age well after *Curculio* speciation, while the oldest element, a LINE/Penelope transposon (*D* = 0.23; ∼39 MYA), provides an upper bound predating speciation. The two *Ccb* insertions on Chromosome 3 are associated with four and one annotated TE, respectively, no Mavericks, and a large repeat-rich block with a mean divergence of *D* = 0.11 across unclassified repeat elements.

This TE-sparse architecture contrasts sharply with the *Rickettsia*-derived HGTs, which are overwhelmingly embedded within Mavericks across all three species: 97% (484/500) in *C. caryae*, 93% (268/289) in *C. nanulus*, and 94% (132/141) in *C. glandium*. Mavericks within and flanking these HGT regions are significantly younger than those in the broader flank-minus-core background (*p* = 3.7 × 10^−48^, see *Methods*; Supplementary Fig. 45). In *C. caryae*, 90 paralogous sequences (representing 82 unique genes; Table 1) reside within these Mavericks (Supplementary Fig. 46), consistent with copy-number amplification of a subset of protein-coding genes following HGT.

**Table 1.** Enriched gene ontology (GO) terms for HGTs within Mavericks.

| GO term | Annotation | Enrichment | Adjusted *p*-value |
| --- | --- | --- | --- |
| Molybdopterin cofactor biosynthetic process<br>GO:0032324 | The chemical reactions and pathways resulting in the formation of the molybdopterin cofactor (Moco). | 251.4 | 0.04623 |
| radial spoke<br>GO:0001534 | Protein complex that links the outer microtubule doublet of a 9+2 type ciliary or flagellar axoneme with the sheath that surrounds the central pair of microtubules. | 125.7 | 0.04623 |
| THO complex part of transcription export complex: GO:0000445 | The THO complex when it is part of the TREX (TRanscription EXport) complex that is involved in coupling transcription to export of mRNAs to the cytoplasm. | 125.7 | 0.04623 |
| isomerase activity<br>GO:0016853 | Catalysis of the geometric or structural changes within one molecule. | 251.4 | 0.04623 |
| SMAD binding<br>GO:0046332 | Binding to a SMAD signaling protein. | 125.7 | 0.04623 |
| acyltransferase activity, acyl groups converted into alkyl on transfer<br>GO:0046912 | Catalysis of the transfer of an acyl group from one compound (donor) to another (acceptor), with the acyl group being converted into alkyl on transfer. | 83.80 | 0.04623 |
| ribonuclease III activity<br>GO:0004525 | Catalysis of the endonucleolytic cleavage of RNA with 5'-phosphomonoesters and 3'-OH termini. | 83.80 | 0.04623 |
| GTPase activator activity<br>GO:0005096 | Binds to and increases the activity of a GTPase. | 18.85 | 0.01846 |
| ligand-gated monoatomic ion channel activity<br>GO:0015276 | Enables the transmembrane transfer of an ion by a channel that opens when a specific ligand has been bound by the channel complex. | 12.89 | 0.04623 |

While *C. caryae* and *C. nanulus* have experienced recent TE proliferation (Supplementary Fig. 45; Table 1), total TE content is 1.7-fold greater in *C. caryae* (762.6 Mb) than in *C. nanulus* (449.6 Mb). Over 60% of this size difference is attributable to the expansion of a handful of TE families: Mavericks, LTR/Gypsy, and DNA/TcMar-Tc1 (Supplementary Fig. 45; Supplementary Tables 4 and 5). Whereas LTR/Gypsy and DNA/TcMar-Tc1 elements are also enriched for young TEs and HGTs in *C. glandium*, Mavericks are not (Supplementary Table 6). Consistent with these lineage-specific differences, the derived species *C. caryae* and *C. nanulus* exhibit both a greater HGT burden and more extensive recent proliferation of type II DNA elements, while *C. glandium* retains fewer HGTs, a smaller genome, and a distinct TE landscape.

### Domestication, pseudogenization, and positive selection shape HGT evolution

To investigate the biological role and evolutionary fate of transferred genes, we examined codon usage bias, intron length, and gene ontology (GO)^53^ enrichment in *C. caryae*, the only species with inferred recent (*Ccb*) and ancient (*Rickettsia*) HGTs. Collectively, *Ccb*-derived genes differ significantly from endogenous weevil genes in codon usage bias and intron length (Fig. 4).

**Figure 4.**
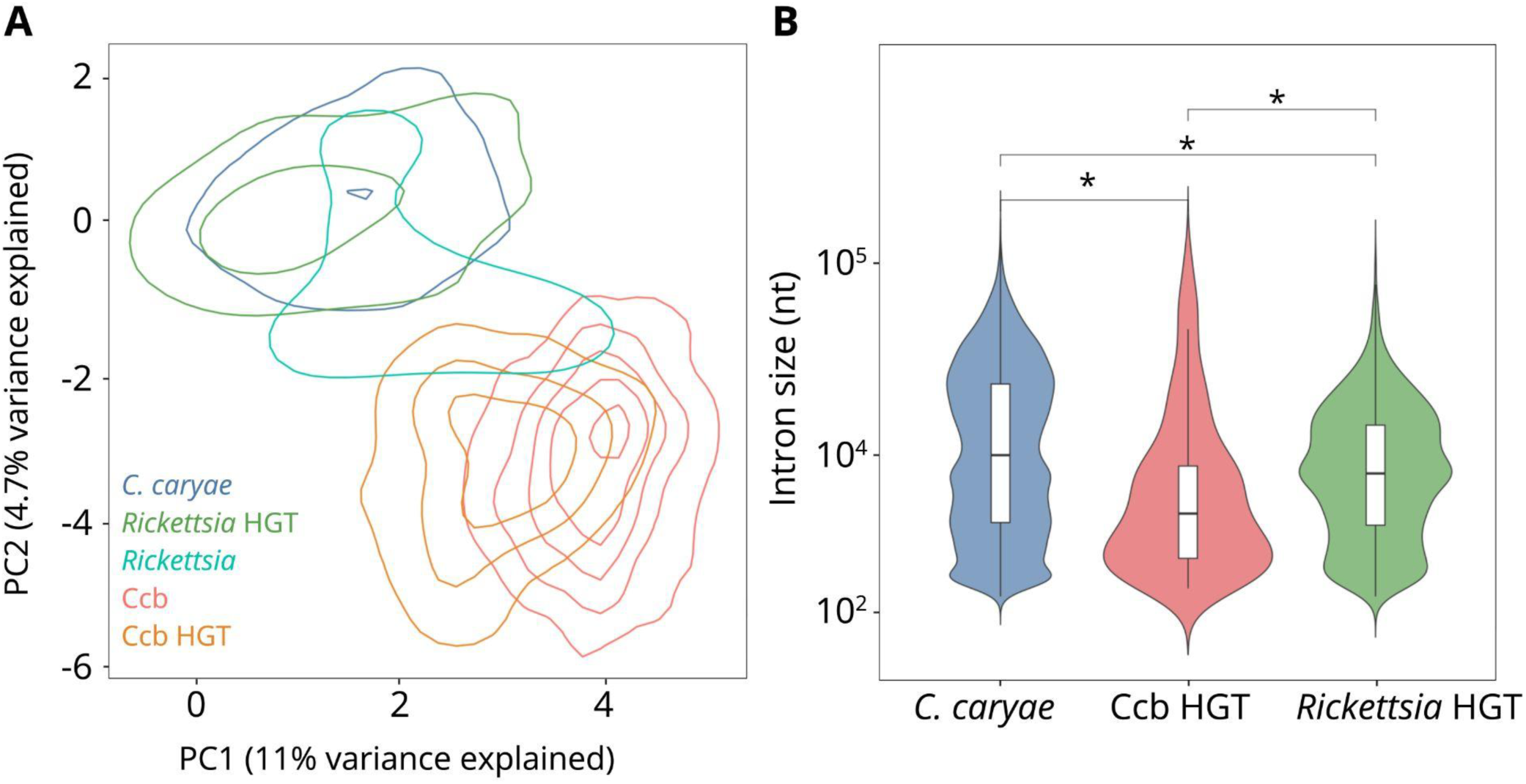
Gene feature differences among host, symbiont, and horizontally transferred coding sequences in *C. caryae*. (A) Two-dimensional kernel density plot of principal component analysis (PCA) based on codon usage frequencies for *C. caryae* genes, Ccb, *Rickettsia*, and HGTs derived from Ccb and *Rickettsia*. (B) Violin plots with embedded box plots summarizing the distribution of intron length in nucleotides (nt) for native *C. caryae* genes, Ccb-derived HGTs, and *Rickettsia*-derived HGTs. * *p* ≤ 10^−4^ (see *Methods*).

We tested for codon-level positive selection using branch-site analyses of 17 genes shared among the Chromosome 1 and 3 HGT insertions, the *Ccb* reference genome, and the distant outgroup *Haemophilus influenzae* (see *Methods*). Eight of these genes show strong evidence of positive selection in the inserted Chromosome 1 and 3 HGT insertions relative to the bacterial outgroups (bolded terms in Supplementary Table 2). The remaining *Ccb*-derived genes encode proteins involved in canonical bacterial metabolic processes, including respiration, stress response, and components of the isoprenoid methylerythritol phosphate pathway, which has not previously been documented in animals (Supplementary Table 2). These genes show little evidence of branch-specific positive selection.

In contrast, older *Rickettsia*-derived HGTs are less distinct from endogenous genes in codon usage bias and intron length, consistent with progressive host domestication and degradation over evolutionary time. Though many of these HGTs are pseudogenized, the retained genes embedded within Mavericks are enriched for functions related to metabolism, transcriptional regulation, and signal transduction (Table 1; Supplementary Fig. 47). Notably, these functional genes reside within TE deserts flanked by young (*D* = 0.035; ∼6.1 MYA), TE-rich regions.

## Discussion

Here, we demonstrate a recurrent association between bacterial HGT and lineage-specific TE proliferation across three *Curculio* species. By integrating comparative pangenomics, structural variation analyses, TE age estimation, and molecular evolutionary inference, we uncover a pattern linking bacterial DNA acquisition with TE proliferation, genome expansion, and molecular innovation. Furthermore, we identify specific genomic features associated with HGT regions and implicate Mavericks in the amplification and persistence of transferred bacterial genes.

Genome size variation remains a longstanding enigma in evolutionary biology. While demographic history, speciation, and environmental stress have been proposed as contributors to genome expansion, these factors are often difficult to distinguish and rarely provide a direct biological source of DNA accumulation. Our analyses identify HGT followed by TE proliferation as a previously underappreciated mechanism that directly adds genomic material and promotes long-term genome evolution. Across *Curculio*, larger genomes are associated with greater HGT burden and more extensive TE proliferation, with *C. caryae* exhibiting the largest genome, the greatest number of HGTs, and the clearest signal of recent TE activity. Differences in HGT burden could reflect partial or complete symbiont loss^54–56^, as *C. glandium* is the only lineage examined here confirmed to retain the bacterial *Ccb* endosymbiont. If so, retention of *Ccb* genes through HGT may compensate for the loss or reduction of the endosymbiont by maintaining advantageous symbiotic functions^55^ in *C. caryae* and *C. nanulus*. Consistent with this hypothesis, genes within transferred regions are indeed enriched for specialized metabolic functions.

Alternatively, differences in HGT abundance may reflect lineage-specific variation in TE activity^57,58^, population size^59^, chromatin structure^60,61^, and recombination rate^62^. Under this model, HGT burden would be shaped primarily by genomic properties that influence the persistence of transferred sequences rather than by differences in symbiont dependence or HGT fitness effects. These explanations are not mutually exclusive and may jointly contribute to the lineage-specific accumulation of HGTs observed in *Curculio*. Future comparative analyses of symbiont status, TE activity, and chromosomal architecture will be necessary to disentangle the drivers of HGT retention.

In addition to the more recent *Ccb*-derived integrations, we identify extensive *Rickettsia*-derived HGTs across all three species. The shared distribution of these genes is consistent with an ancestral transfer event predating the *Curculio* radiation, followed by lineage-specific amplification, retention, and decay. The persistence of ancient *Rickettsia*-derived HGTs alongside younger *Ccb*-derived integrations in *C. caryae* suggests that bacterial DNA acquisition is not restricted to a single historical event, but rather represents a recurrent source of genomic novelty. Notably, both *Rickettsia*-derived HGTs and *Ccb*-derived HGTs occur in structural variation hotspots characterized by young, abundant, and heterogeneous TEs. The repeated co-occurrence of HGTs with these TE-rich structural hotspots suggests that certain genomic environments may function as persistent regions of both TE activity and HGT integration (Fig. 3). We refer to these regions as TE proliferation springs because they appear to facilitate TE proliferation and the genome-wide dissemination of HGT-derived genes.

Although multiple DNA II transposable element families contribute to recent genome expansion in *C. caryae* and *C. nanulus,* Mavericks are disproportionately associated with HGT regions across all three species, as they have been observed in eukaryotic nematodes^38^. Their enrichment within HGT cores and flanking regions suggests they play a central role in the capture, amplification, and long-term persistence of *Rickettsia-*derived HGTs. Furthermore, their discrete age structure is consistent with episodic proliferation across time, paralleling the expansion of HGTs following integration. More broadly, our genome-wide analyses identify TE density as the strongest predictor of HGT. TE-independent HGTs were enriched in gene-dense regions, whereas TE-associated HGTs occurred predominantly within repeat-rich regions.

Central to our evolutionary model, HGTs linked to the youngest TEs exhibit the strongest association with TE density, with approximately a ten-fold increase in odds relative to background windows, compared with a three-fold increase for HGTs associated with old TEs. In contrast, HGTs associated with intermediate-age TEs show no significant enrichment. This temporal pattern is consistent with acute, episodic bursts of TE proliferation surrounding a subset of HGTs, followed by progressive decay through turnover and sequence divergence. The lack of TE-density enrichment among HGTs associated with intermediate-age TEs implies that HGT-associated TE proliferation is not characterized by continuous decay but instead occurs in punctuated episodes. HGTs associated with young TEs likely reflect recent proliferation, whereas those associated with old TEs may represent remnants of earlier proliferation events. HGTs associated with intermediate-age TEs may comprise heterogeneously aged or degraded TEs, resulting in genomic regions that more closely resemble background windows. Repeated cycles of HGT acquisition, TE proliferation, and subsequent decay may therefore contribute to genome expansion through the accumulation of both TE-derived and HGT-derived DNA.

Our analyses further suggest that the evolutionary fate of transferred genes is shaped by amplification and decay following integration into the host genome. The ancestral, shared *Rickettsia*-derived HGTs exhibit copy number variation and are distributed throughout the genome, embedded within Mavericks on every chromosome (Supplementary Fig. 48). Domestication of *Rickettsia*-derived HGTs within these elements is inferred from less pronounced, but still significant, codon usage bias and intron structure relative to endogenous *Curculio* genes (Fig. 4). Functional annotation of these retained genes reveals enrichment for transcriptional regulation, metabolism, and signal transduction pathways (Table 1).

The *C. caryae*-specific *Ccb*-derived transfers are localized to four discrete, gene-rich regions: one on Chromosome 1, two on Chromosome 3, and one on an extrachromosomal contig. Pairwise sequence comparisons show that the extrachromosomal contig is nearly identical to the Chromosome 1 region and substantially more similar to both Chromosome 3 regions than to the *Ccb* reference genome. These relationships indicate that the four *Ccb*-derived regions have undergone distinct evolutionary histories following integration and that the Chromosome 1 region and extrachromosomal contig share a more recent common ancestor than either Chromosome 3 region. In contrast, all four regions are similarly diverged from the extant *C. glandium* endosymbiont *Ccb* genome, suggesting that the *C. caryae* donor lineage was already differentiated from the contemporary endosymbiont or may represent an unsampled *Curculioniphilus*-like lineage. Additional evidence will be required to test this hypothesis. *Ccb*-derived genes also exhibit distinct codon usage bias and shorter introns, indicating comparatively limited genetic domestication by the *C. caryae* host genome. Nevertheless, several transferred genes show signatures of positive selection relative to their corresponding *Ccb* genes, suggesting that a subset of transferred genes may be undergoing rapid adaptive evolution following integration.

Collectively, these findings support an evolutionary model in which foreign DNA invasion, TE-mediated amplification, selective retention, and progressive decay interact to shape the long-term evolutionary trajectory of genomes (Fig. 5). Under this framework, HGTs contribute molecular novelty to host genomes while interacting with TE-rich genomic environments that facilitate their persistence and amplification. Our results suggest that recurrent cycles of foreign DNA acquisition may contribute to genome size variation across evolutionary timescales and provide a framework for evaluating whether similar processes operate in other eukaryotic lineages.

**Figure 5.**
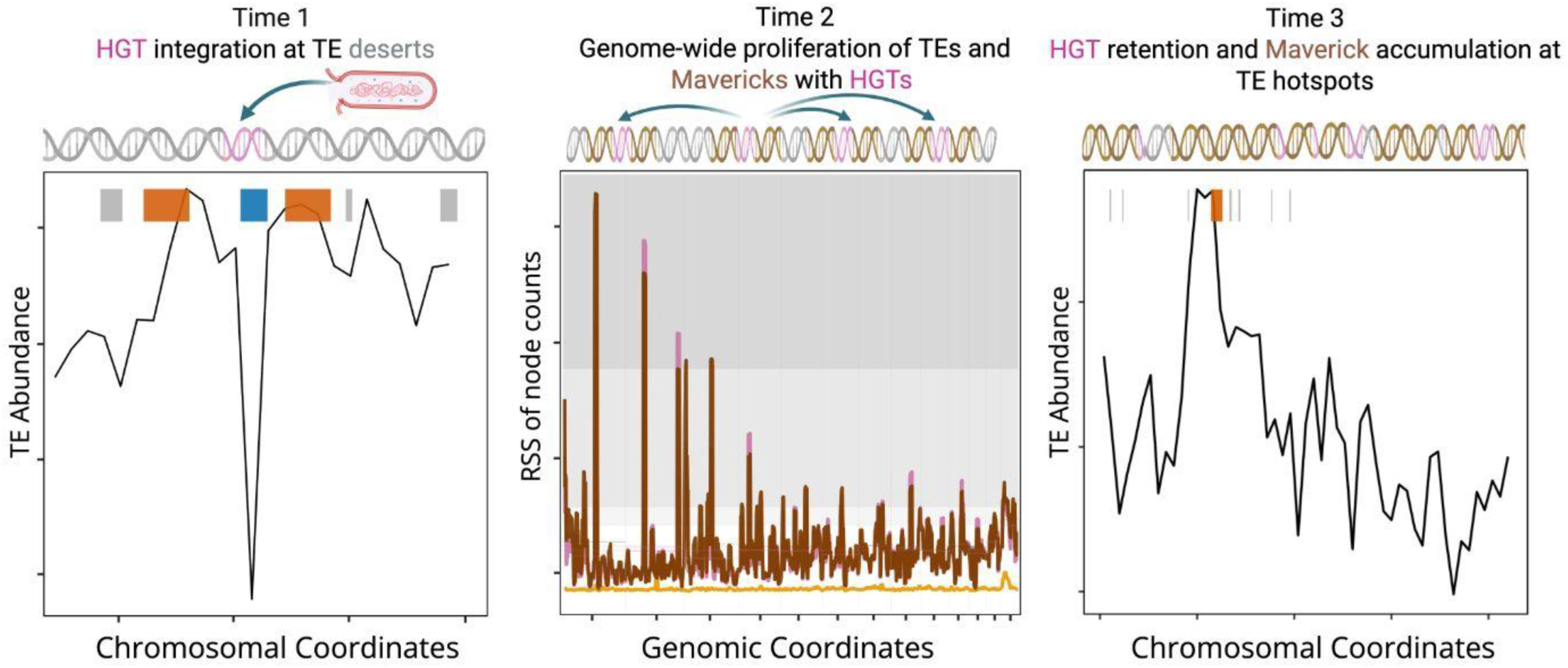
Evolutionary model for genome size variation. Each panel depicts chromosomal organization (top helix) and an associated genomic landscape profile (bottom plot) through evolutionary time. (Time 1) Initial horizontal transfer of bacterial DNA (pink) occurs within a TE-depauperate region (gray). The TE abundance profile below the schematic illustrates local TE density across chromosomal coordinates, with a TE-poor insertion site (blue), flanking TE-rich regions (orange), and moderately TE-abundant regions extending outward (gray). (Time 2) Genome-wide proliferation of TEs, including Mavericks (light brown), restructures the genomic landscape and facilitates redistribution of HGT-derived sequences across the genome (see also Fig. 3). The lower plot shows RSS of node counts across several species determined from the pangenome graph, reflecting increasing structural complexity during TE expansion. During this phase, genome size increases through TE accumulation surrounding the HGT region. (Time 3) Over evolutionary time, HGT-derived sequences undergo diversification, fragmentation, and pseudogenization within TE-rich regions. The TE abundance profile indicates persistence of TE-dense regions coinciding with HGT-derived regions. Retention of Maverick-HGT complexes on smaller chromosomes suggests long-term turnover of TE/HGT-rich regions rather than continuous genome expansion.

## Methods

### Ortholog identification and phylogenetic reconstruction

Genome assemblies for *L. decemlineata* (GCA_024712935.1), *C. glandium* (icCurGlan1.1), and *C. caryae* (icCurCar2.1)^63^ were retrieved from NCBI, and the *C. nanulus* assembly was accessed at https://doi.org/10.6084/m9.figshare.30546164. Super-scaffolding of *C. nanulus* was performed using RagTag v2.1.0^64^ with *C. caryae* as the reference. To account for potential biases introduced by reference-guided scaffolding of *C. nanulus* and manual super-scaffolding of *C. caryae* and *C. glandium*, we identified scaffold breakpoints from stretches of 100 Ns (ambiguous nucleotides) and restricted node-centric analyses to regions located at least five kb from these regions^65^.

Single-copy orthologs across the three *Curculio* species and *L. decemlineata* were identified using BUSCO v6^66^ with the Insecta odb12 reference. These genes were aligned using MAFFT v7.526^67^ and trimmed for shared missing data using ClipKIT v2.6.1^68^. Gene trees and best-fitting substitution models were inferred for each alignment using IQ-TREE v2.4.0^69^. A time-calibrated ultrametric tree was estimated using MCMCtree in PAML^70,71^, with two independent runs to assess convergence. Analyses employed an independent rates model, an approximate likelihood method^70^, and a prior on the origin of Phytophaga in the early Triassic (245-255 MYA)^65,72^. Orthologs were used to infer chromosomal homology based on species-specific genomic coordinates. Putative HGT regions were validated by remapping long-read sequences to the source assemblies and confirming that these regions do not coincide with read termini or contig breakpoints.

*Ccb-*derived genomic regions were identified in the *C. caryae* reference genome by nucleotide similarity searches against the *Ccb* reference genome using the blastn program of BLAST v2.15.0+^73^. These regions were subsequently isolated from the genome using the samtools^74^ v1.22.1 faidx command based on the reciprocal blastn results. Length-weighted nucleotide divergence was estimated by pooling mismatches across all ungapped aligned sites within each region and applying the Jukes-Cantor (JC69) correction^75^ using a custom Python script^65^. *Ccb-*derived genes within these regions were also identified by blastn searches against the *Ccb* reference gene set and visualized in R v4.3.2^76^ using the UpSetR v1.4.0^77^ package.

### Pangenome construction

A pangenome graph for the three *Curculio* species was constructed using PGGB v0.7.4^78^ for each homologous chromosome, based on synteny inferred from shared single-copy ortholog alignments derived from BUSCO results. Node size and genomic position relative to the *C. glandium* reference were quantified using ODGI v0.9.2-0-gbe6a0202^79^ with the –view and –position arguments, respectively^65^.

To identify node-dense regions representing local enrichments of structural variation, we quantified lineage-specific node counts in consecutive one Mb windows across each chromosome. These values were summarized using a rolling five Mb root sum of squares (RSS) statistic, providing a measure of local graph complexity, with higher values indicating greater accumulation of structural variation. Specifically, for each lineage and chromosome, node counts were calculated per one Mb window, and the magnitude of local structural variation was estimated as

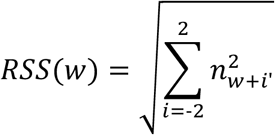

where *n_w_* represents the node count in the focal window *w*. Because this approach summarizes lineage-specific node density rather than pairwise divergence, *C. glandium* was used as the coordinate reference for graph position^65^.

We focused primarily on the 90th percentile of node density to retain a broad set of structurally dynamic windows and maximize sensitivity to lineage-specific variation. To determine physical associations with HGT regions and TEs, node positions were mapped back to source genome coordinates using the – position argument in ODGI^79^.

### TE identification and age estimation

EarlGrey v2.0.7^80^ was used with default settings^65^ to identify, annotate, and estimate the age of TEs in all genome assemblies, with time calibration based on the *L. decemlineata* mutation rate^81^. TE age and genomic coordinates for all *Curculio* species were compared with node and HGT coordinates^65^. By integrating TE age, location, and type with structural variant and HGT coordinates, we estimated the approximate timing of these events relative to *Curculio* speciation.

### HGT identification and evolutionary characterization

Endosymbiont gene discovery was conducted in *C. glandium*^82^, *C. nanulus*^83^, and *C. caryae*. Searches were performed using BLASTN v2.15.0+^73^ against the NCBI nucleotide database, restricted to bacterial sequences with more than 10 million accessions (accessed March 2025). Putative hits were conservatively retained if they were at least one kb in length, had an e-value ≤10^−10^, and a bitscore ≥70. Additional screening for redundancy among bacterial taxa blast hits was performed by retaining the top accession within overlapping genomic ranges based on e-value and bitscore^65^. The curated BLAST tables were then analyzed for each species^65^.

*Ccb*-derived HGTs identified at Chromosomes 1 and 3 were aligned to homologous gene sequences from ancestral *Ccb* and *H. influenzae* bacteria using MACSE v2.07^84^. Alignments were tested for positive selection by iteratively alternating the foreground branches (Chromosome 1, both regions on Chromosome 3, and the branch leading to both Chromosomes 1 and 3) using the codeml program in PAML v4.10.9^70^. For each gene and focal branch, likelihood ratio tests compared alternative and null models for significance based on one degree of freedom. Likelihood ratio tests were evaluated against the 50:50 mixture null, appropriate for the branch-site test, with log-likelihood ratio test statistics >5.41 considered significant at α = 0.01. Codons from the favored branch-site models were identified using Bayes empirical Bayes posterior probabilities, and sites with posterior probability >0.95 were retained for interpretation^65^. *Rickettsia*-derived HGTs identified across the *C. caryae* genome were tested for GO enrichment with *rrvgo* v1.14.2^85^ in R v4.3.2^76^ and confirmed with a Fisher’s exact test with false discovery rate correction implemented in the R package *stats* v4.3.2^76^.

Codon usage bias and gene structure were measured in putative *Rickettsia*– and *Ccb*-derived HTGs from the *C. caryae* genome assembly relative to host and symbiont reference genes. Codon usage was measured across coding sequences (cDNA) for 1) *Rickettsia*-derived HGTs, 2) *Ccb*-derived HGTs, 3) *C.caryae* ribosomal-associated genes, 4) endogenous *C. cary*ae genes, 4) the *Ccb* reference genome, and 5) the *Rickettsia* endosymbiont of *Ceutorhynchus assimilis* using the Measure Independent of Length and Composition (MILC) metric^86^ implemented in the R package coRdon v1.24.0^86^. Gene structure was inferred from the BRAKER v3^87^ Iso-Seq gene annotations (*braker.gtf*). Intron lengths were calculated from annotated exon coordinates by summing the lengths of intronic intervals for each predicted gene^65^. Genes overlapping HGT regions were subsequently identified using genomic coordinates and BLAST results, and total intron lengths were compared between HGT-derived and endogenous *C. caryae* genes using Wilcoxon rank sum and signed rank tests in R^76^.

### Permutation analyses for HGT-matched genomic regions

Genomic regions matching the number and size of HGT cores, along with their two kb flanking regions, were sampled uniformly at random across the pecan weevil reference genome (icCurCar2.1) 10,000 times. TE density was calculated for each sampled region, generating null distributions for core-like and flank-like regions against which the observed HGT cores, flanks, and genomic backgrounds were compared^65^.

### Multinomial regression modeling of HGT-associated regions

Genomic predictors associated with HGT regions were also evaluated using two multinomial regression models of the form

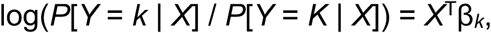

where *Y* ∈ {1, 2, …, *K*} is the response class label for *K* classes (with the final class *K* treated as the reference class), *X*^T^ = [1, *X*_1_, *X*_2_, …, *X_p_*] is a (*p*+1)-dimensional vector of *p* genomic predictors (including the intercept), and β*_k_* = [β*_k_*_,0_, β*_k_*_,1_, β*_k_*_,2_, …, β*_k_*_,*p*_] is the associated (*p*+1)-dimensional vector of regression coefficients for class *k*.

These models were fitted to non-overlapping 5 kb windows of the pecan weevil reference genome^65^. HGT-categorized windows, defined as those in which ≥20% of a 5 kb window (≥1 kb) overlapped an HGT annotation, were further distinguished by their association with TEs. Separate analyses were performed for TE-associated and TE-independent HGT compartments. All models were fitted using 500 chromosome-matched background subsampling replicates, in which HGT-core and flanking windows were retained while background windows were randomly resampled without replacement within chromosomes. All predictors were standardized (*z*-transformed) to have a mean of zero and a standard deviation of one prior to analysis. Because regression coefficients (β*_kj_*) are related to the log odds ratio, odds ratios were calculated as exp(β*_kj_*), and percent changes in odds were calculated as (exp(β*_kj_*) – 1) × 100.

The first model distinguished HGT windows based on the presence or absence of TE annotations. The four response classes (*K* = 4) were: TE-associated HGT cores (class *Y* = 1), TE-independent HGT cores (class *Y =* 2), flanking windows (class *Y* = 3), and genomic background (class *Y* = 4, reference class *K*). This model included *p* = 3 genomic predictors: coding sequence density (*X*_cds_), GC content (*X*_gc_), and GC skew (*X*_gc_skew_), yielding the input vector *X* = [1, *X*_cds_, *X*_gc_skew_, *X*_gc_]. Coding sequence density was log-transformed using ln(1+*x*).

The second model distinguished TE-associated HGT windows based on TE annotations age. These windows were stratified by TE age using the 25th and 75th percentiles for young (*D* < 0.02) and old (*D* > 0.07) TEs, respectively. The five response classes (*K* = 5) were: Young (class *Y =* 1), intermediate (class *Y* = 2), old (class *Y* = 3), flanking regions (class *Y* = 4), and genomic background (class *Y* = 5, reference class *K*). This model included *p* = 4 genomic predictors: coding sequence density (*X*_cds_), TE density (*X*_te_), GC content (*X*_gc_), and GC skew (*X*_gc_skew_), yielding input vector *X* = [1, *X*_cds_, *X*_gc_skew_, *X*_gc_, *X*te]. TE and coding sequence densities were log-transformed using ln(1+*x*).

## Data availability

Code associated with this study is archived on Zenodo (DOI:10.5281/zenodo.19598494) and publicly available on GitHub^65^ (https://github.com/zpcohen/Curculio_HTE). The datasets generated and analyzed during this study, together with all files required to reproduce the reported results, are available through Figshare at 10.6084/m9.figshare.32605731.

## Acknowledgements

This work was supported by the National Institutes of Health (2R35GM128590 to M.D.; R35GM142438 to R.A.) and the National Science Foundation (DBI-2130666 to R.A. and M.D.). Computations for this research were performed using the services provided by Research Computing at Florida Atlantic University and resources provided by the SCINet project of the USDA Agricultural Research Service (ARS project number 0201-88888-003-000D).

## References

1. Huang, W. et al. Natural variation in genome architecture among 205 *Drosophila melanogaster* Genetic Reference Panel lines. Genome Res. 24, 1193–208 (2014).

2. Gregory, T.R. Genome size evolution in animals. In The Evolution of the Genome (ed. Gregory, T.R.) 3–87 (Elsevier, San Diego, 2005).

3. Ellis, L.L. et al. Intrapopulation Genome Size Variation in *D. melanogaster* Reflects Life History Variation and Plasticity. PLoS Genet. 10, e1004522 (2014).

4. Yawako, W., et al. Extensive Copy Number Variation Explains Genome Size Variation in the Unicellular Zygnematophycean Alga, Closterium peracerosum–strigosum–littorale Complex. evad115 Genome Biol. Evol. 15, evad115 (2023).

5. Koonin, E.V. et al. A comprehensive evolutionary classification of proteins encoded in complete eukaryotic genomes. Genome Biol 5, R7 (2004).

6. Thomas, C.A. The genetic organization of chromosomes. Annu. Rev. Genet. 5, 237–256 (1971).

7. Elliott, T. A. & Gregory, T. R. What’s in a genome? The C-value enigma and the evolution of eukaryotic genome content. Philos. Trans. R. Soc. B 370, 20140331 (2015).

8. Van Straalen, N. I. & Roelofs, D. Introduction to Ecological Genetics (Oxford Univ. Press, New York, 2006).

9. Eddy, S. R. The C-value paradox, junk DNA and ENCODE. Curr. Biol. 22, R898–R899 (2012).

10. Blommaert J., Genome size evolution: towards new model systems for old questions. Proc Biol Sci 1; 287 1933 (2020).

11. Huettel, B., et al. Effects of aneuploidy on genome structure, expression, and interphase organization in *Arabidopsis thaliana*. PLoS Genet. 4, e1000226 (2008).

12. Paterson, A. H., et al. Repeated polyploidization of *Gossypium* genomes and the evolution of spinnable cotton fibres. Nature 492, 423–427 (2012).

13. Zande, P.V., Zhou, X., and Selmecki, A. The dynamic fungal genome: polyploidy, aneuploidy and copy number variation in response to stress. Annu. Rev. Microbiol. 77, 341–361 (2023).

14. SanMiguel, P., et al. The paleontology of intergene retrotransposons of maize. Nat. Genet. 20, 43–45 (1998).

15. Kidwell, M.G. Transposable elements and the evolution of genome size in eukaryotes. Genetica 115, 49–63 (2002).

16. Locke, D.P. et al. Large-scale variation among human and great ape genomes determined by array comparative genomic hybridization. Genome Res. 13, 347–357 (2003).

17. Lee, S.I., Kim, N.S. Transposable elements and genome size variations in plants. Genomics Inform. 12, 87–97 (2014).

18. Naville, M., et al. Massive changes of genome size driven by expansions of non-autonomous transposable elements. Curr. Biol. 29, 1161–1168 (2019).

19. Wang D., et al. Which factors contribute most to genome size variation within angiosperms? Ecol. Evol. 11, 2660–2668 (2021).

20. Belyayev, A. Bursts of transposable elements as an evolutionary driving force. J. Evol. Biol. 27, 2573–2584 (2014).

21. Kalendar, R., et al. Genome evolution of wild barley (Hordeum spontaneum) by BARE-1 retrotransposon dynamics in response to sharp microclimatic divergence. Proc. Natl Acad. Sci. USA 97, 6603–6607 (2000).

22. de Boer, J.G., et al. Bursts and horizontal evolution of DNA transposons in the speciation of pseudotetraploid salmonids. BMC Genomics 8, 422 (2007).

23. Belyayev, A., et al. Transposable elements in a marginal plant population: temporal fluctuations provide new insights into genome evolution of wild diploid wheat. Mob. DNA 1, 6 (2010).

24. Chénais, B. et al. The impact of transposable elements on eukaryotic genomes: From genome size increase to genetic adaptation to stressful environments. Gene 509, 7–15 (2012).

25. Charlesworth, B., Charlesworth, D. The population dynamics of transposable elements. Genet. Res. 42, 1–27 (1983).

26. Bourgeois, Y. & Boissinot, S. On the Population Dynamics of Junk: A Review on the Population Genomics of Transposable Elements. Genes 10, 419 (2019).

27. Muñoz-López, M. & García-Pérez, J.L. DNA transposons: nature and applications in genomics. Curr. Genomics 11, 115–128 (2010).

28. Hughes, J. & Vogler, A.P. The phylogeny of acorn weevils (genus Curculio) from mitochondrial and nuclear DNA sequences: the problem of incomplete data. Mol. Phylogenet. Evol. 32, 601–615 (2004).

29. Keeling, P. & Palmer, J. Horizontal gene transfer in eukaryotic evolution. Nat Rev Genet 9, 605–618 (2008).

30. Archibald, J. Endosymbiosis and Eukaryotic Cell Evolution. Current Biology, 25, R911–R921 (2015).

31. Husnik, F. & McCutcheon, J. Functional horizontal gene transfer from bacteria to eukaryotes. Nat Rev Microbiol 16, 67–79 (2018).

32. Thomas, C. & Nielsen, K. Mechanisms of, and Barriers to, Horizontal Gene Transfer between Bacteria. Nat Rev Microbiol 3, 711–721 (2005).

33. Riley, D. R., et al. Bacteria-Human Somatic Cell Lateral Gene Transfer Is Enriched in Cancer Samples. PLOS Computational Biology 9(6): e1003107 (2013).

34. R. Stouthamer, J. A. J. Breeuwer & G. D. D. Hurst. *Wolbachia Pipientis*: Microbial Manipulator of Arthropod Reproduction. Annual Review Microbiology. 53:71–102 (1999).

35. Jensen, L. et al. Assessing the effects of a sequestered germline on interdomain lateral gene transfer in Metazoa, *Evolution*, Volume 70, Issue 6, 1 1322–1333 (1999).

36. Schaack, S., Gilbert, C. & Feschotte, C. Promiscuous DNA: horizontal transfer of transposable elements and why it matters for eukaryotic evolution. Trends in Ecology & Evolution 25, 537–546 (2010).

37. Pritham, E. J. et al. Mavericks, a novel class of giant transposable elements widespread in eukaryotes and related to DNA viruses, Gene 390, 3–17 (2007).

38. Widen, S. A. et al. Virus-like transposons cross the species barrier and drive the evolution of genetic incompatibilities. Science 380, eade0705 (2023).

39. Zhan, D., et al. Horizontal gene transfer in insects: Insights into origins, detection, and functional roles. The Innovation Life 4:100223. (2026).

40. Ewart, K. M., et al. Uncovering thousands of endosymbiont DNA transfer events within single cockroach genomes, Proc. Natl. Acad. Sci. U.S.A. 123 (25) e2604240123, (2026).

41. Wybouw, N. et al. Horizontal Gene Transfer Contributes to the Evolution of Arthropod Herbivory, *Genome Biology and Evolution*, Volume 8, Issue 6, 1785–1801 (2016).

42. Rinke, J.L., et al. Comparative analysis of 163 ant genomes reveals recurrent horizontal gene transfer from bacteria to ants, GigaScience, giag043 (2026).

43. Henry, L., et al. Horizontally Transmitted Symbionts and Host Colonization of Ecological Niches. Current Biology, 23, 1713–1717 (2013).

44. Wybouw, N., et al. A gene horizontally transferred from bacteria protects arthropods from host plant cyanide poisoning eLife 3:e02365, (2014).

45. Pedezzi, R., et al. A novel β-fructofuranosidase in Coleoptera: characterization of a β-fructofuranosidase from the sugarcane weevil, *Sphenophorus levis*. Insect Biochem Mol Biol (2014).

46. Daimon, T., et al. Beta-fructofuranosidase genes of the silkworm, Bombyx mori: insights into enzymatic adaptation of B. mori to toxic alkaloids in mulberry latex J Biol Chem, 283, 15271–15279 (2008).

47. Zhao, C., Doucet, D., Mittapalli, O., Characterization of horizontally transferred beta-fructofuranosidase (ScrB) genes in *Agrilus planipennis*. Insect Mol Biol, 23, 821–832 (2014).

48. Toju, H. et al. “*Candidatus* Curculioniphilus buchneri,” a Novel Clade of Bacterial Endocellular Symbionts from Weevils of the Genus *Curculio*. Appl. Environ. Microbiol. 76, 275–282 (2010).

49. Naito, K. et al. Unexpected consequences of a sudden and massive transposon amplification on rice gene expression. Nature 461, 1130–1134 (2009).

50. Mynhardt, G., Harris, M.K., Cognato, A.I. Population Genetics of the Pecan Weevil (Coleoptera: Curculionidae) Inferred from Mitochondrial Nucleotide Data. Ann. Entomol. Soc. Am. 100, 582–590 (2007).

51. Shin, S. et al. Phylogenomic Data Yield New and Robust Insights into the Phylogeny and Evolution of Weevils. Mol. Biol. Evol. 35, 823–836 (2018).

52. Kimura, M. A simple method for estimating evolutionary rates of base substitutions through comparative studies of nucleotide sequences. J. Mol. Evol. 16, 111–120 (1980).

53. Ashburner, M., *et. al.* Gene ontology: tool for the unification of biology. Nature Genetics, 25: 25–29 (2000)

54. Hotopp, J. C. D. Horizontal gene transfer between bacteria and animals. Trends in Genetics, 27: 157–163 (2011).

55. Casiraghi, M. Mapping the presence of *Wolbachia* pipientis on the phylogeny of filarial nematodes: evidence for symbiont loss during evolution. Int. J. Parasitol. 34:191–203 (2004).

56. McNulty, S.N., et al. Endosymbiont DNA in Endobacteria-Free Filarial Nematodes Indicates Ancient Horizontal Genetic Transfer. PLOS ONE 5(6): e11029 (2010).

57. Groth, S. B., & Blumenstiel, J. P. Horizontal Transfer Can Drive a Greater Transposable Element Load in Large Populations, Journal of Heredity, 108, 1: 36–44 (2017).

58. Rezvykh, A.P. et al. Transposable elements as drivers of genome evolution in *Drosophila virilis*, Nucleic Acids Research. Nucleic Acids Res. gkag139 (2026).

59. Clark, A.G., et al. Evolution of genes and genomes on the Drosophila phylogeny. Nature. 450, 203–218 (2007).

60. Bushman, F. Targeting Survival: Integration Site Selection by Retroviruses and LTR-Retrotransposons. Cell. 115, 135–138 (2003).

61. Zhou, J. & Eickbush, T. H. The Pattern of R2 Retrotransposon Activity in Natural Populations of Drosophila simulans Reflects the Dynamic Nature of the rDNA Locus. PLOS Genetics 5(2): e1000386 (2009).

62. Nguyen, A.N.T., et al. Recombination resolves the cost of horizontal gene transfer in experimental populations of *Helicobacter pylori*, Proc. Natl. Acad. Sci. U.S.A. 119 (12) e2119010119 (2022).

63. Perkin, L.C. et al., A chromosome level reference genome for the pecan weevil, *Curculio caryae*. Sci. Data 13, 703 (2026).

64. Alonge, M. et al. Automated assembly scaffolding elevates a new tomato system for high-throughput genome editing. Genome Biol 23, 258 (2022).

65. Cohen, Z. Curculio HTE pipeline (v1.0.0). Zenodo. DOI:10.5281/zenodo.19598494 (2026).

66. Manni, M., et al. BUSCO update: novel and streamlined workflows along with broader and deeper phylogenetic coverage for scoring of eukaryotic, prokaryotic, and viral genomes. Mol. Biol. Evol. 38, 4647–4654 (2021).

67. Katoh, K. & Standley, D.M. MAFFT multiple sequence alignment software version 7: improvements in performance and usability. Mol. Biol. Evol. 30, 772–780 (2013).

68. Steenwyk, J.L. et al. (2020) ClipKIT: A multiple sequence alignment trimming software for accurate phylogenomic inference. PLoS Biol. 18, e3001007 (2020).

69. Nguyen, L.T., et al. IQ-TREE: a fast and effective stochastic algorithm for estimating maximum-likelihood phylogenies. Mol. Biol. Evol. 32, 268–274 (2015).

70. Yang, Z. PAML: a program package for phylogenetic analysis by maximum likelihood. Comput. Appl. Biosci. 13, 555–556 (1997).

71. Yang, Z. PAML 4: Phylogenetic Analysis by Maximum Likelihood. Mol. Biol. Evol. 24, 1586–1591 (2007).

72. dos Reis, M. & Yang, Z. Approximate likelihood calculation for Bayesian estimation of divergence times. Mol. Biol. Evol. 28, 2161–2172 (2011).

73. Camacho, C., et al., BLAST+: architecture and applications. BMC Bioinformatics 10, 421 (2009).

74. Li H., et al. 1000 Genome Project Data Processing Subgroup. The Sequence Alignment/Map format and SAMtools. Bioinformatics. Aug 15;25(16):2078–9 (2009).

75. Jukes, T.H., Cantor, C.R. (1969). Evolution of Protein Molecules. New York: Academic Press. pp. 21–132.

76. R Core Team. R: A language and environment for statistical computing. R Foundation for Statistical Computing, Vienna, Austria (2021).

77. Conway J. R., Lex A., & Gehlenborg N., UpSetR: an R package for the visualization of intersecting sets and their properties, Bioinformatics, 33, 8: 2938–2940 (2017).

78. Garrison E et al., Building pangenome graphs. Nat. Methods 21, 2008–2012 (2024).

79. Guarracino, A., et al. ODGI: understanding pangenome graphs. Bioinformatics 38, 3319–3326 (2022).

80. Baril, T., et al. A Fully Automated User-Friendly Transposable Element Annotation and Analysis Pipeline. Mol. Biol. Evol. 41, msae068 (2024).

81. Xu, S., et al. Trio-sequencing reveals high germline mutation rates in the Colorado potato beetle (*Leptinotarsa decemlineata*). Genome Biol. Evol. 18, evag027 (2026).

82. The Darwin Tree of Life Project Consortium, Sequence locally, think globally: The Darwin Tree of Life Project, Proc. Natl Acad. Sci. USA 119, e2115642118 (2022).

83. Davis, D.D., et al. Whole-genome assembly and annotation of the acorn weevil, Curculio nanulus (Coleoptera: Curculionidae). G3 Genes|Genomes|Genetics. 16, jkaf292 (2025)

84. Ranwez, V., et al. MACSE v2: Toolkit for the Alignment of Coding Sequences Accounting for Frameshifts and Stop Codons. Mol. Biol. Evol. 35, 2582–2584 (2018).

85. Sayols, S. rrvgo: a Bioconductor package for interpreting lists of Gene Ontology terms. microPublication Biology (2023).

86. Supek, F. & Vlahovicek, K. Comparison of codon usage measures and their applicability in prediction of microbial gene expressivity. BMC Bioinformatics. 9;6:182. Erratum in: BMC Bioinformatics. 2010;11:463 (2005).

87. Gabriel, L., et al. BRAKER3: Fully automated genome annotation using RNA-seq and protein evidence with GeneMark-ETP, AUGUSTUS, and TSEBRA. Genome Res. 34(5):769–777 (2024).

