## Supplementary for "Bacterial DNA invasion triggers transposable element proliferation and genome expansion"

**Supplementary material for *Bacterial DNA invasion triggers transposable element proliferation and genome expansion***

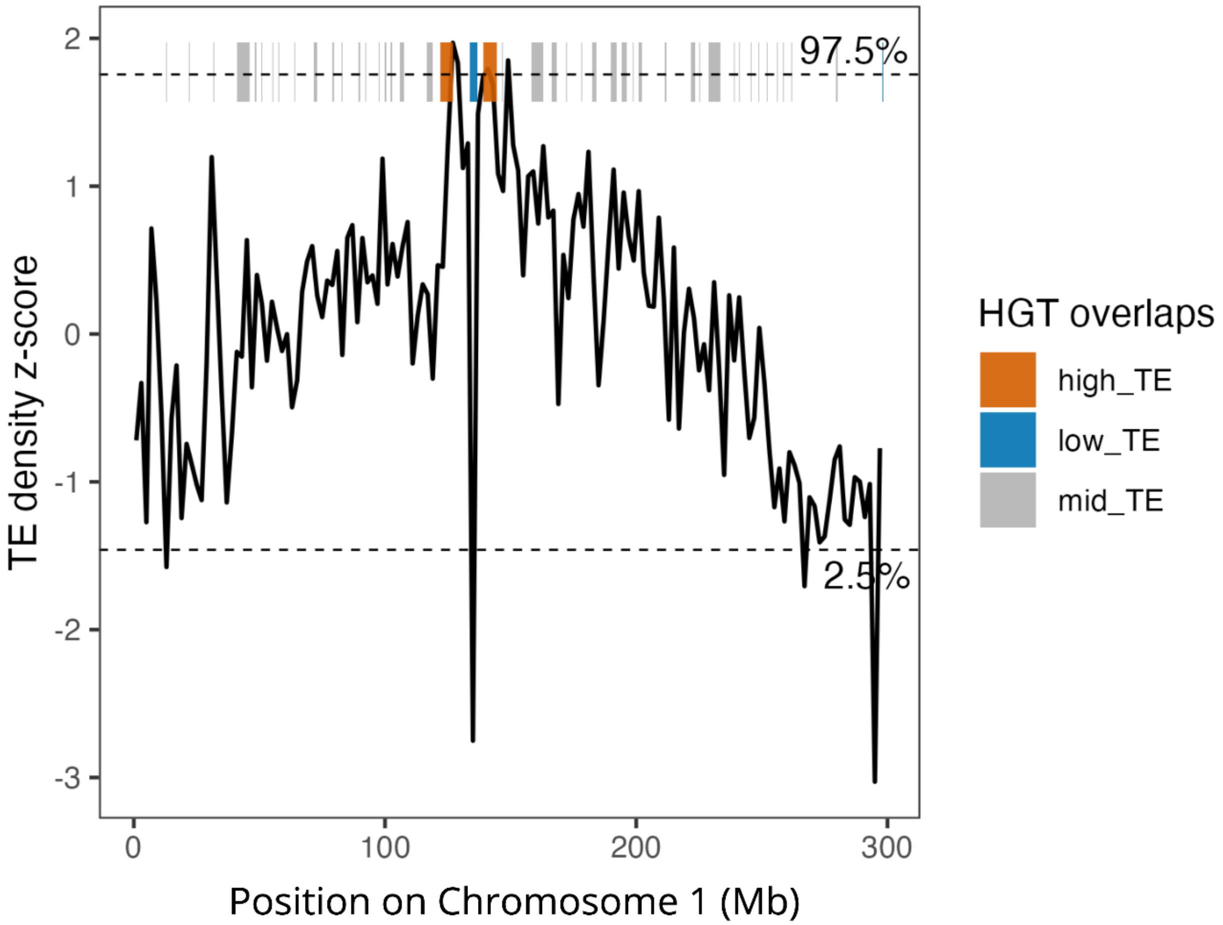


Supplementary Figure 1. Z-score transformed TE density in sliding 2 Mb windows across Chromosome 1 of *C. caryae*. HGTs located within the upper (97.5th percentile) and lower (2.5th percentile) tails of the TE density distribution are indicated by orange and blue bars, respectively. Gray bars indicate HGTs falling outside these tails.

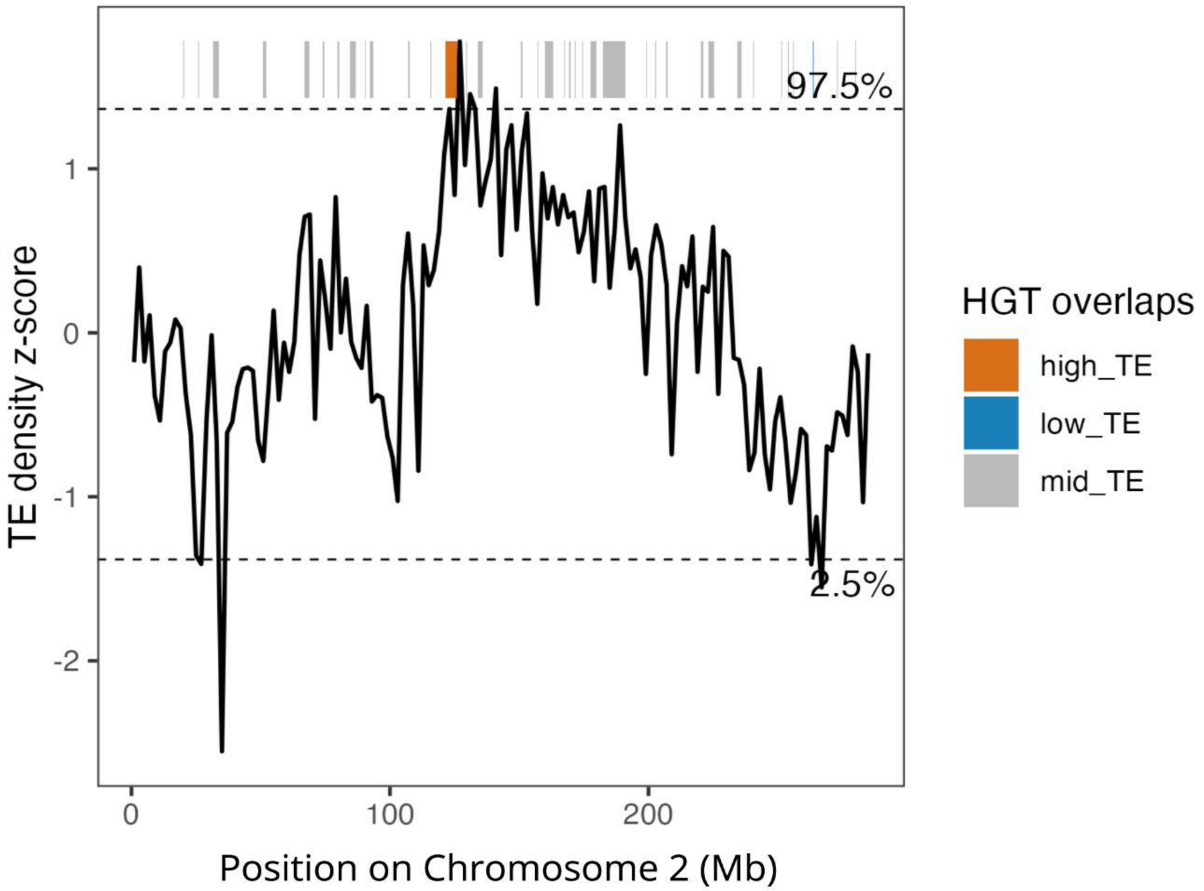


Supplementary Figure 2. Z-score transformed TE density by type in sliding 2 Mb windows across Chromosome 2 of *C. caryae*. HGTs located within the upper (97.5th percentile) and lower (2.5th percentile) tails of the TE density distribution are indicated by orange and blue bars, respectively. Gray bars indicate HGTs falling outside these tails.

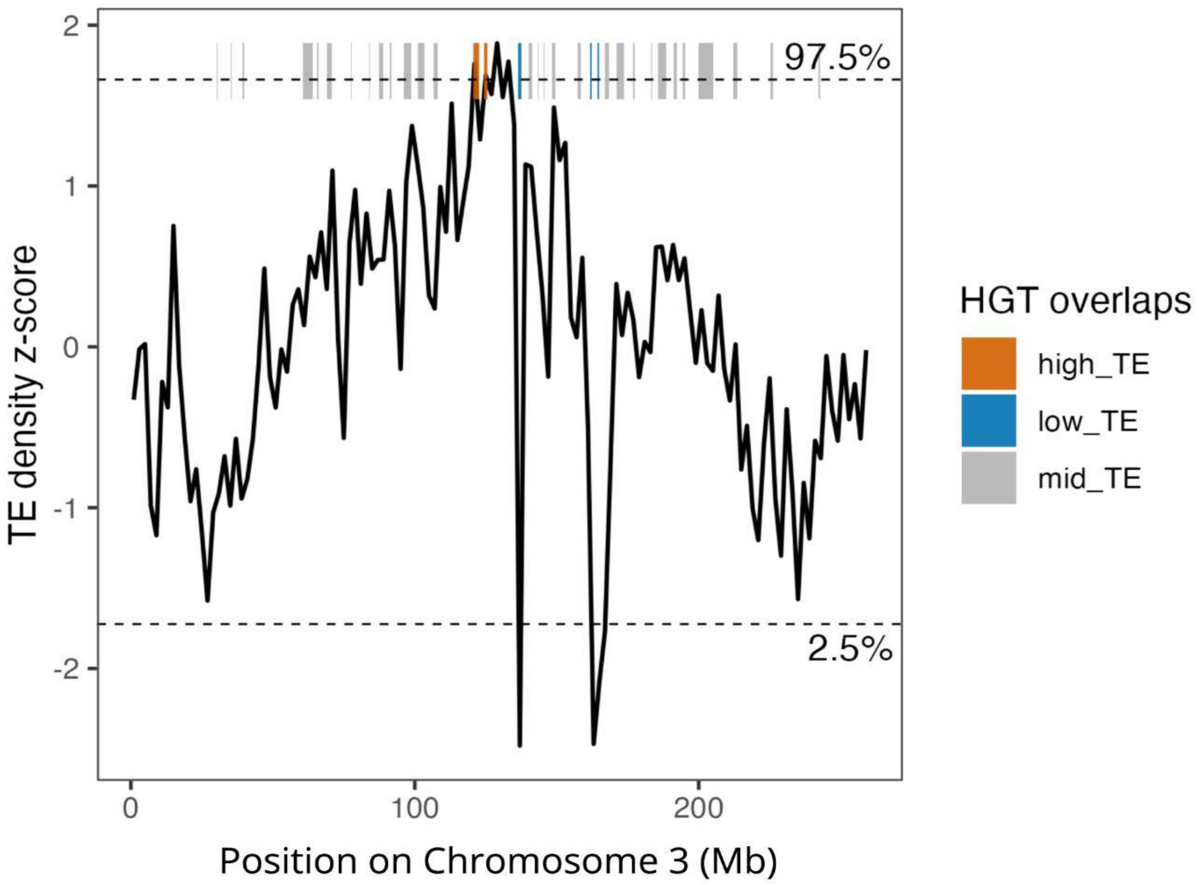


Supplementary Figure 3. Z-score transformed TE density by type in sliding 2 Mb windows across Chromosome 3 of *C. caryae*. HGTs located within the upper (97.5th percentile) and lower (2.5th percentile) tails of the TE density distribution are indicated by orange and blue bars, respectively. Gray bars indicate HGTs falling outside these tails.

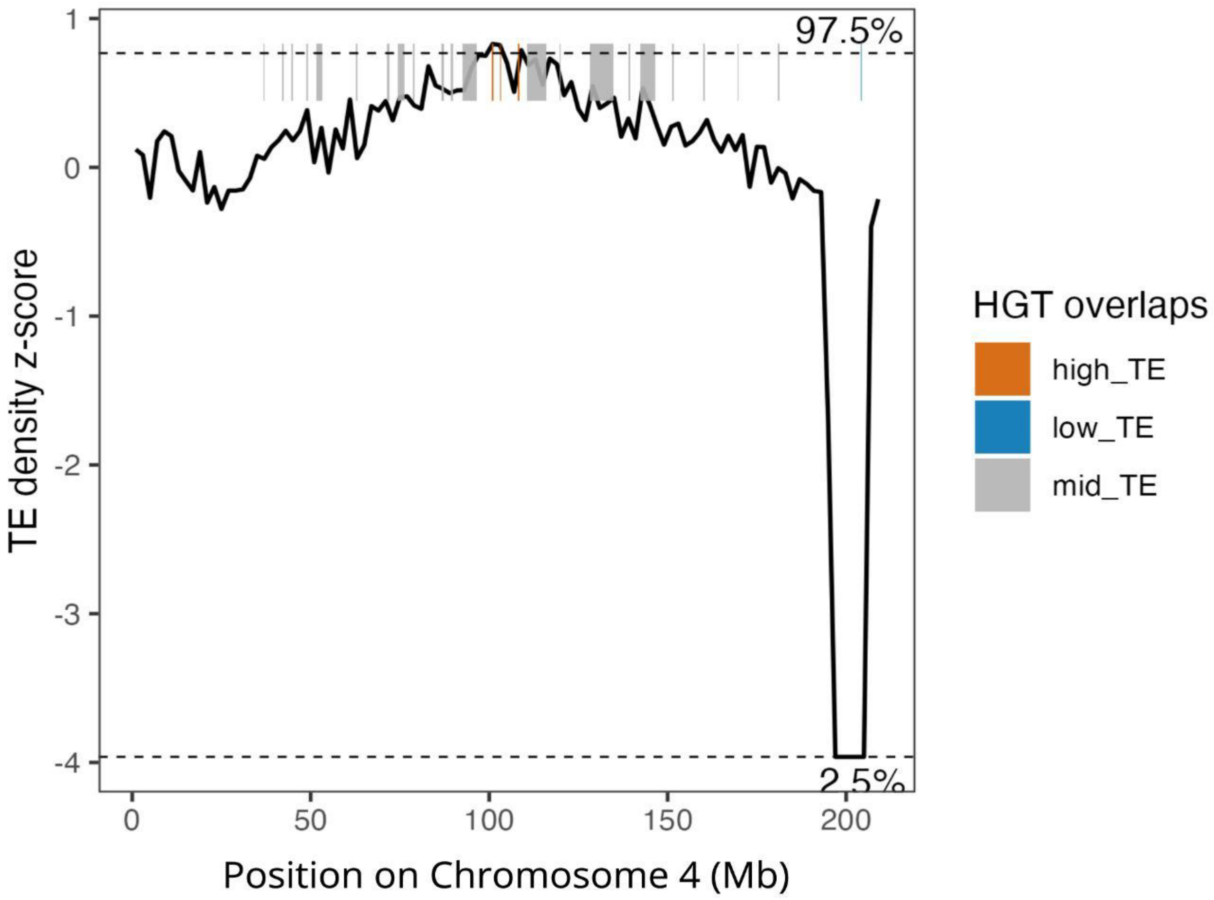


Supplementary Figure 4. Z-score transformed TE density by type in sliding 2 Mb windows across Chromosome 4 of *C. caryae*. HGTs located within the upper (97.5th percentile) and lower (2.5th percentile) tails of the TE density distribution are indicated by orange and blue bars, respectively. Gray bars indicate HGTs falling outside these tails.

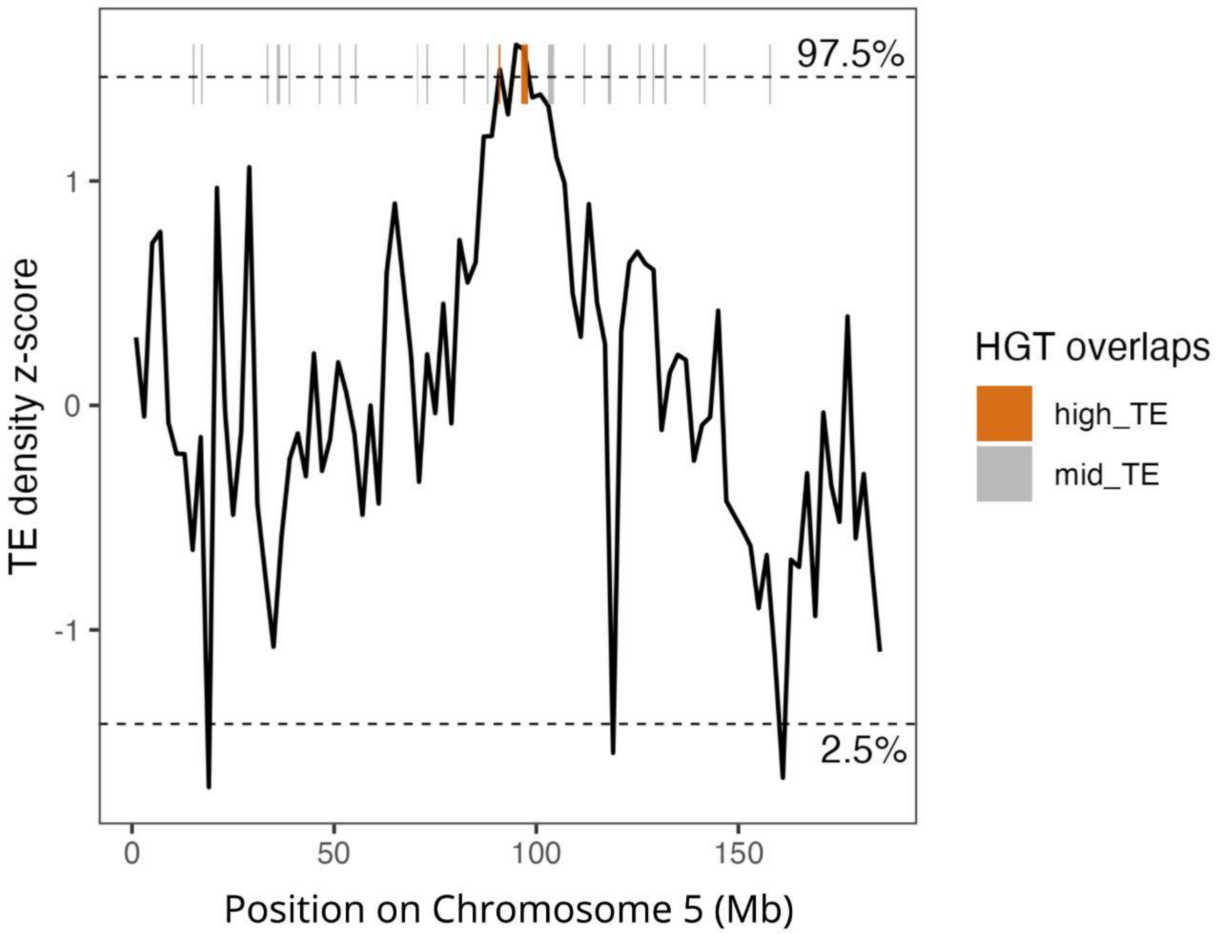


Supplementary Figure 5. Z-score transformed TE density by type in sliding 2 Mb windows across Chromosome 5 of *C. caryae*. HGTs located within the upper (97.5th percentile) tail of the TE density distribution are indicated by orange bars. Gray bars indicate HGTs falling outside this tail.

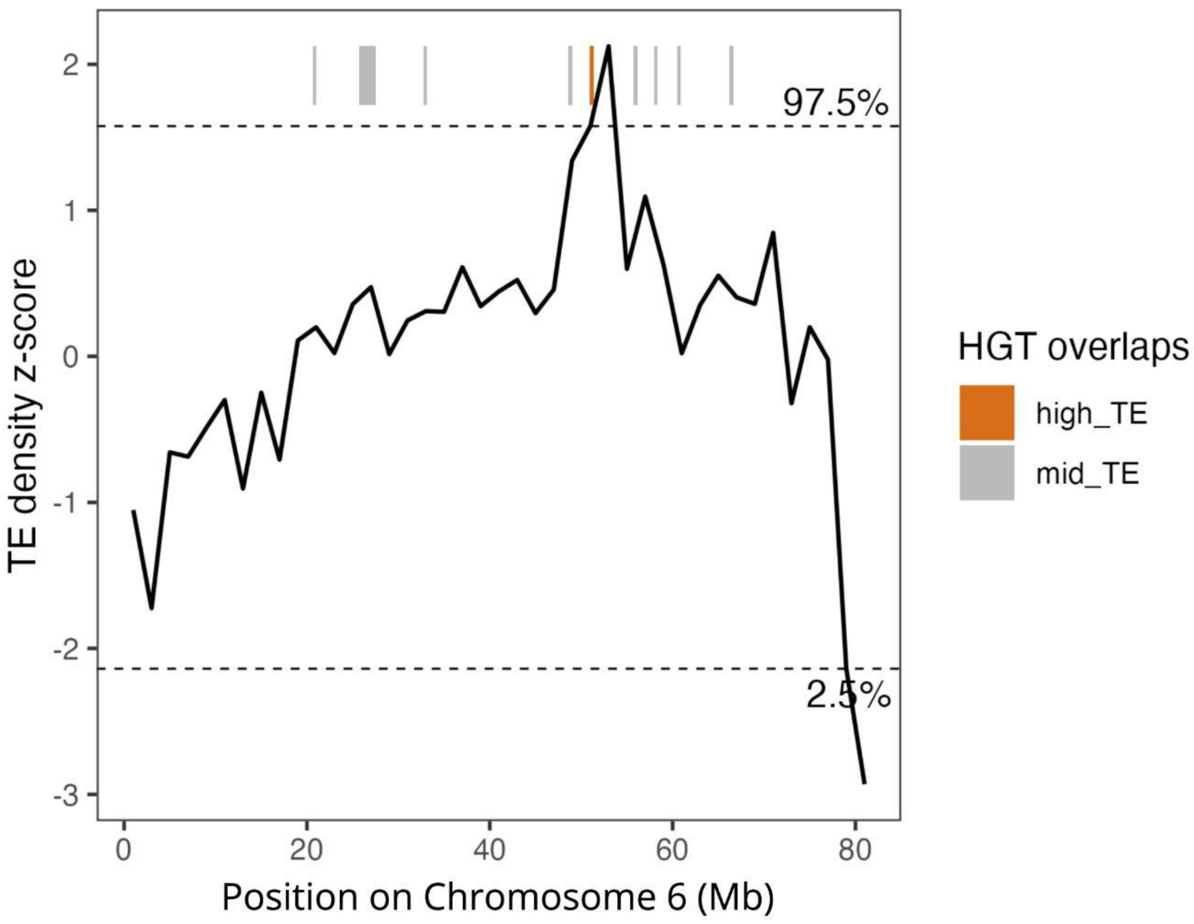


Supplementary Figure 6. Z-score transformed TE density by type in sliding 2 Mb windows across Chromosome 6 of *C. caryae*. HGTs located within the upper (97.5th percentile) and lower (2.5th percentile) tails of the TE density distribution are indicated by orange and blue bars, respectively. Gray bars indicate HGTs falling outside these tails.

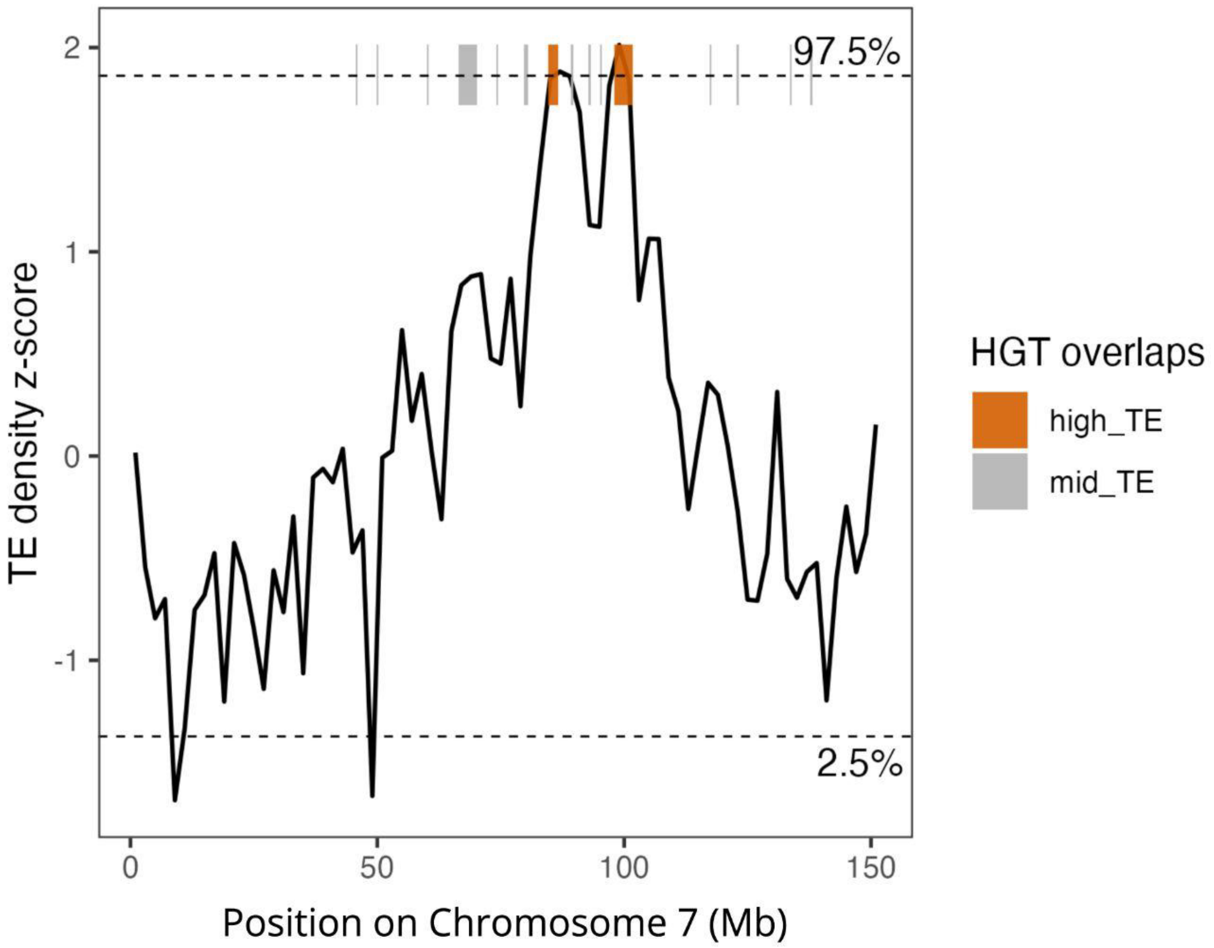


Supplementary Figure 7. Z-score transformed TE density by type in sliding 2 Mb windows across Chromosome 7 of *C. caryae*. HGTs located within the upper (97.5th percentile) tail of the TE density distribution are indicated by orange bars. Gray bars indicate HGTs falling outside this tail.

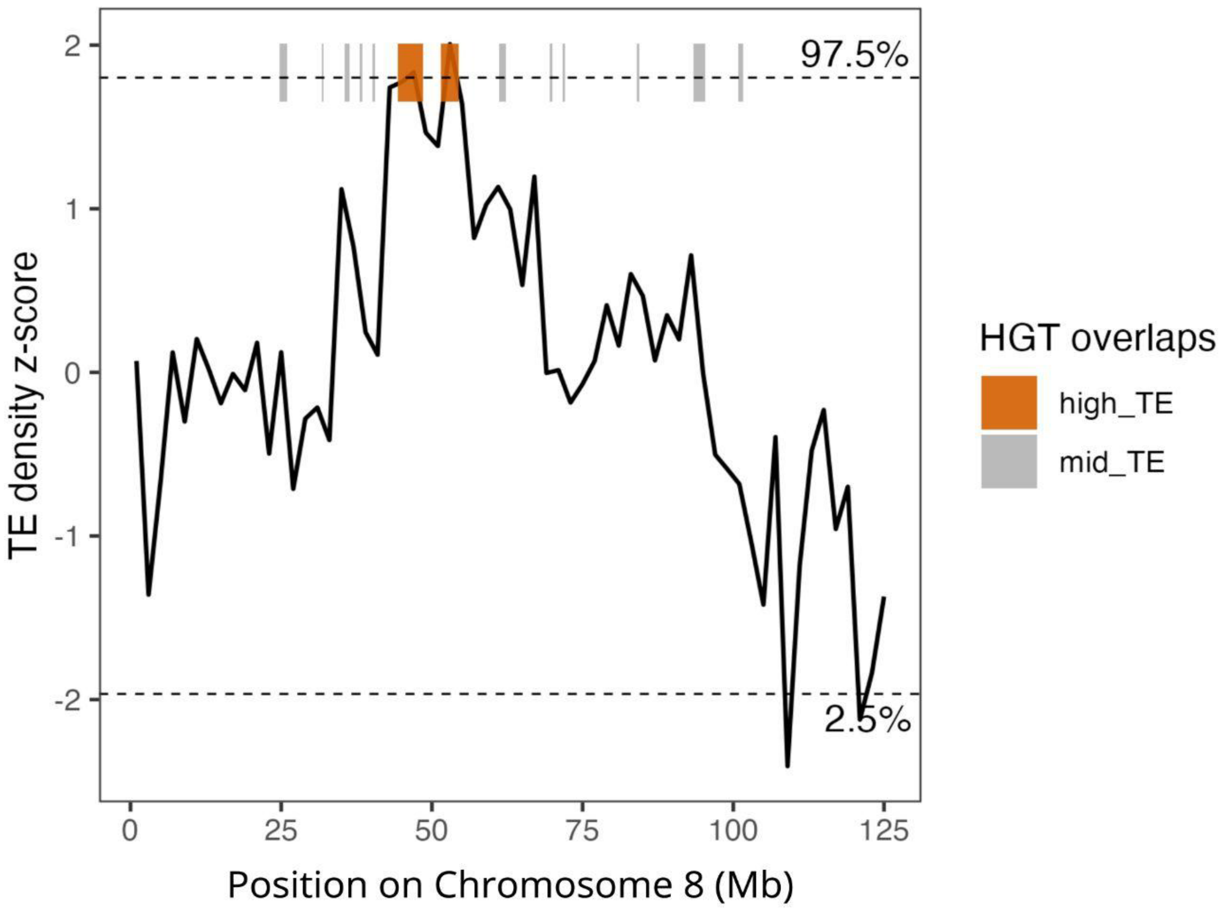


Supplementary Figure 8. Z-score transformed TE density by type in sliding 2 Mb windows across Chromosome 8 of *C. caryae*. HGTs located within the upper (97.5th percentile) tail of the TE density distribution are indicated by orange bars. Gray bars indicate HGTs falling outside this tail.

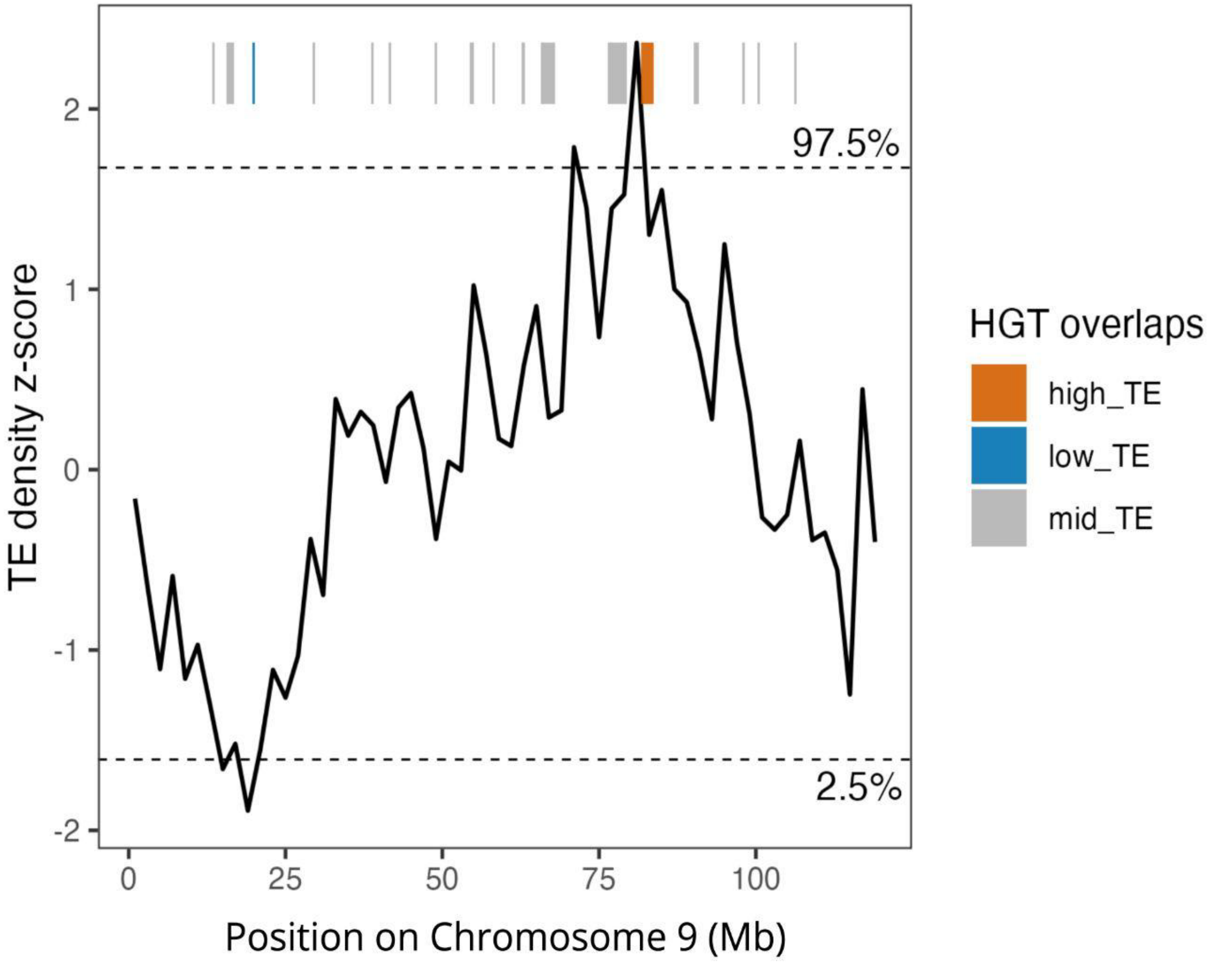


Supplementary Figure 9. Z-score transformed TE density by type in sliding 2 Mb windows across Chromosome 9 of *C. caryae*. HGTs located within the upper (97.5th percentile) and lower (2.5th percentile) tails of the TE density distribution are indicated by orange and blue bars, respectively. Gray bars indicate HGTs falling outside these tails.

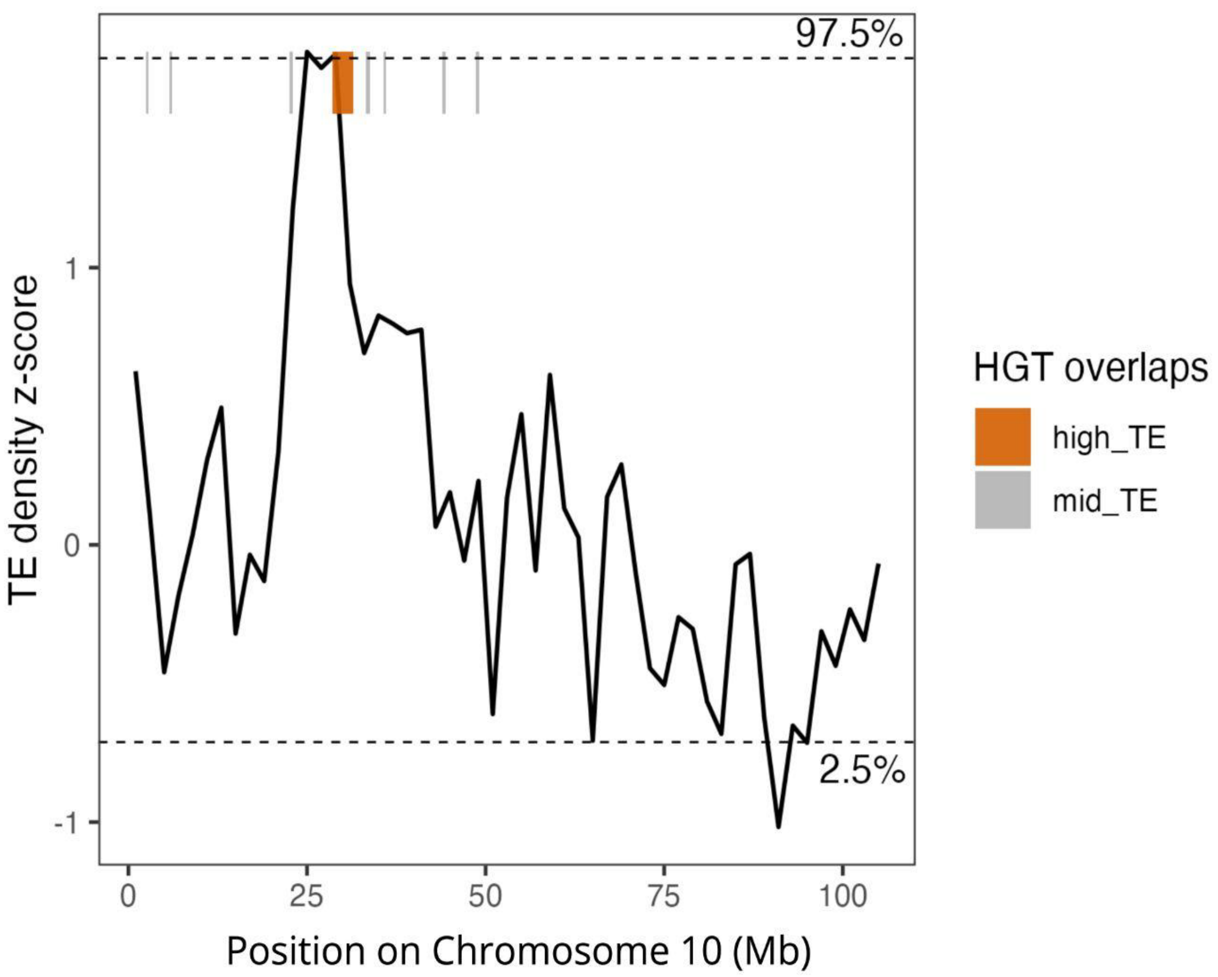


Supplementary Figure 10. Z-score transformed TE density by type in sliding 2 Mb windows across Chromosome 10 of *C. caryae*. HGTs located within the upper (97.5th percentile) tail of the TE density distribution are indicated by orange bars. Gray bars indicate HGTs falling outside this tail.

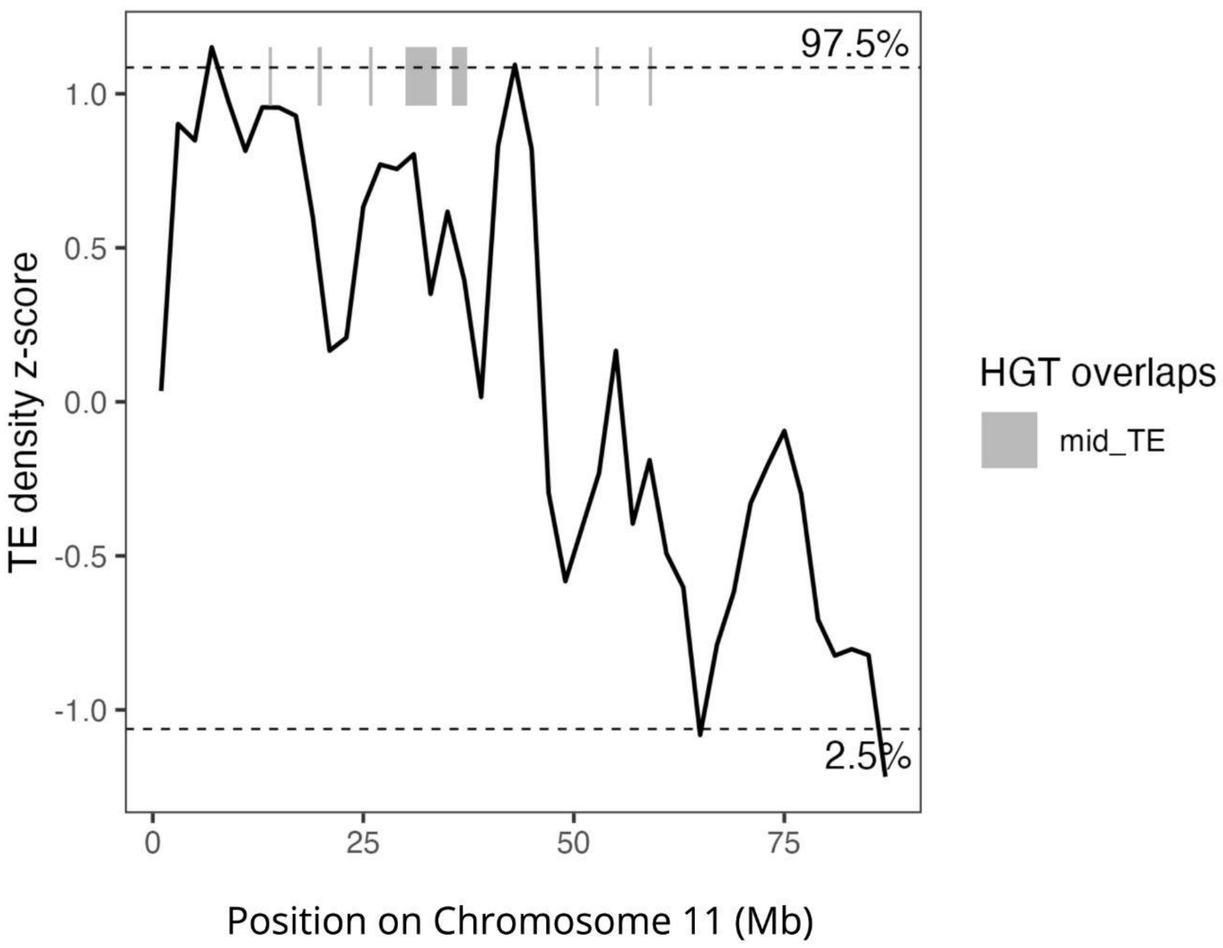


Supplementary Figure 11. Z-score transformed TE density by type in sliding 2 Mb windows across Chromosome 11 of *C. caryae*. Horizontally transferred DNA sequences (HGTs) located outside of the tails of the TE density distribution are indicated with gray bars.

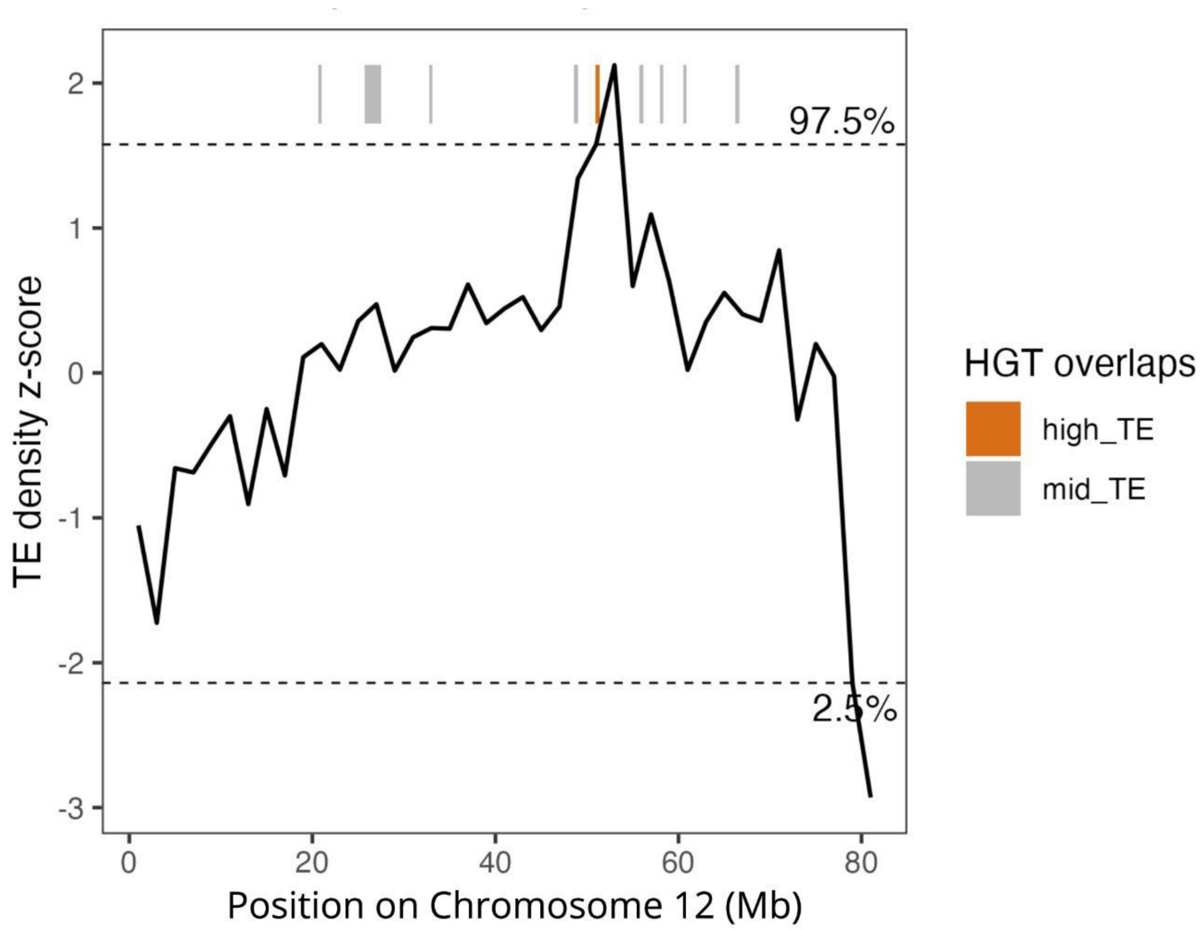


Supplementary Figure 12. Z-score transformed TE density by type in sliding 2 Mb windows across Chromosome 12 of *C. caryae*. HGTs located within the upper (97.5th percentile) tail of the TE density distribution are indicated by orange bars. Gray bars indicate HGTs falling outside this tail.

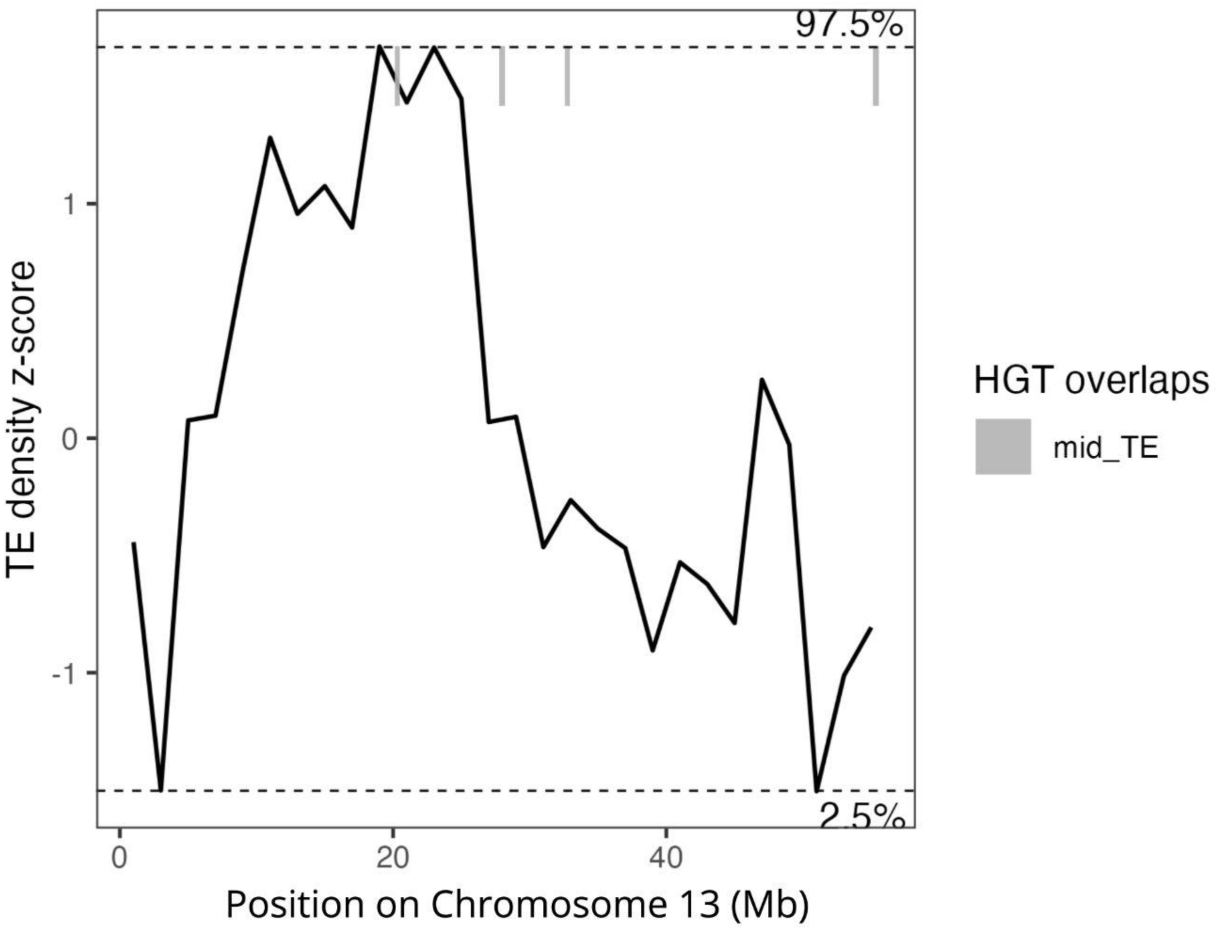


Supplementary Figure 13. Z-score transformed TE density by type in sliding 2 Mb windows across Chromosome 13 of *C. caryae*. Horizontally transferred DNA sequences (HGTs) located outside of the tails of the TE density distribution are indicated with gray bars.

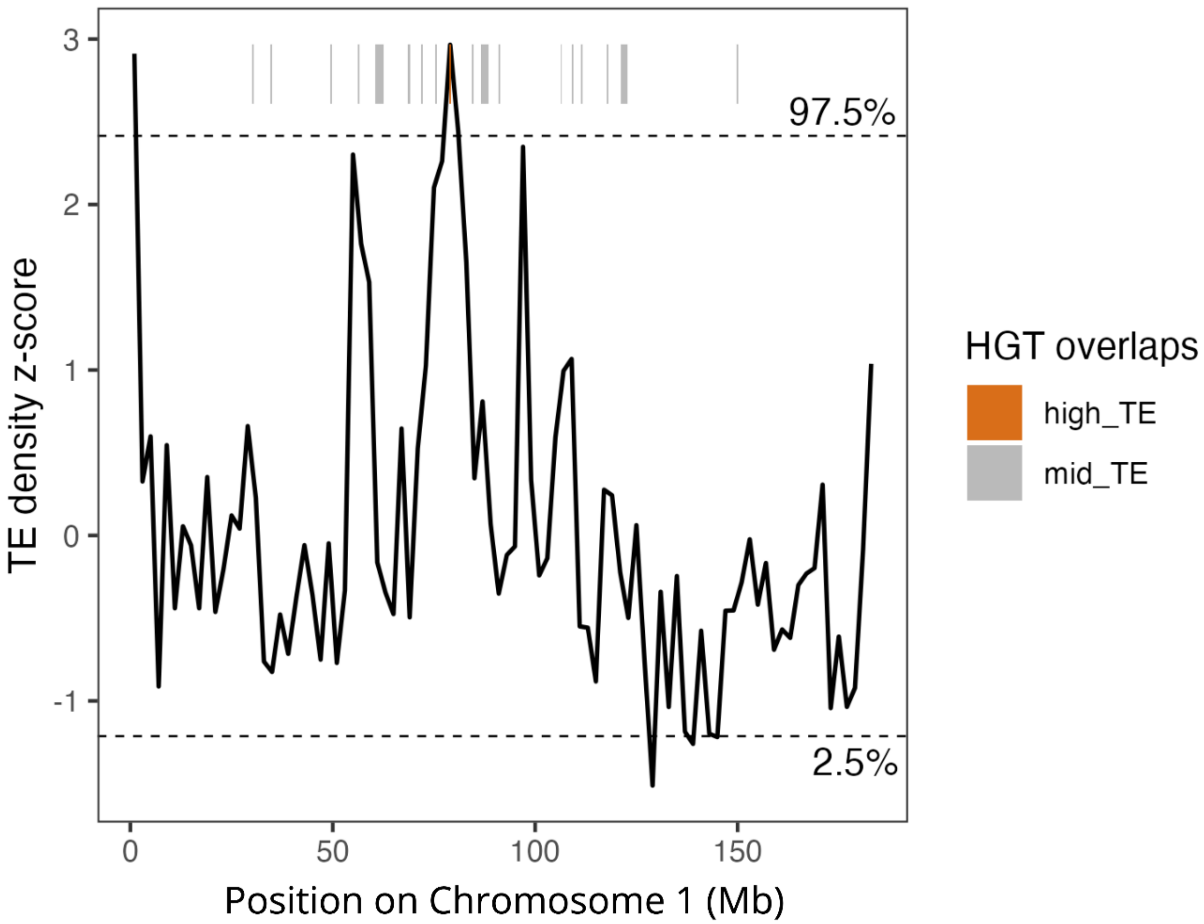


Supplementary Figure 14. Z-score transformed TE density by type in sliding 2 Mb windows across Chromosome 1 of *C. nanulus*. HGTs located within the upper (97.5th percentile) tail of the TE density distribution are indicated by orange bars. Gray bars indicate HGTs falling outside this tail.

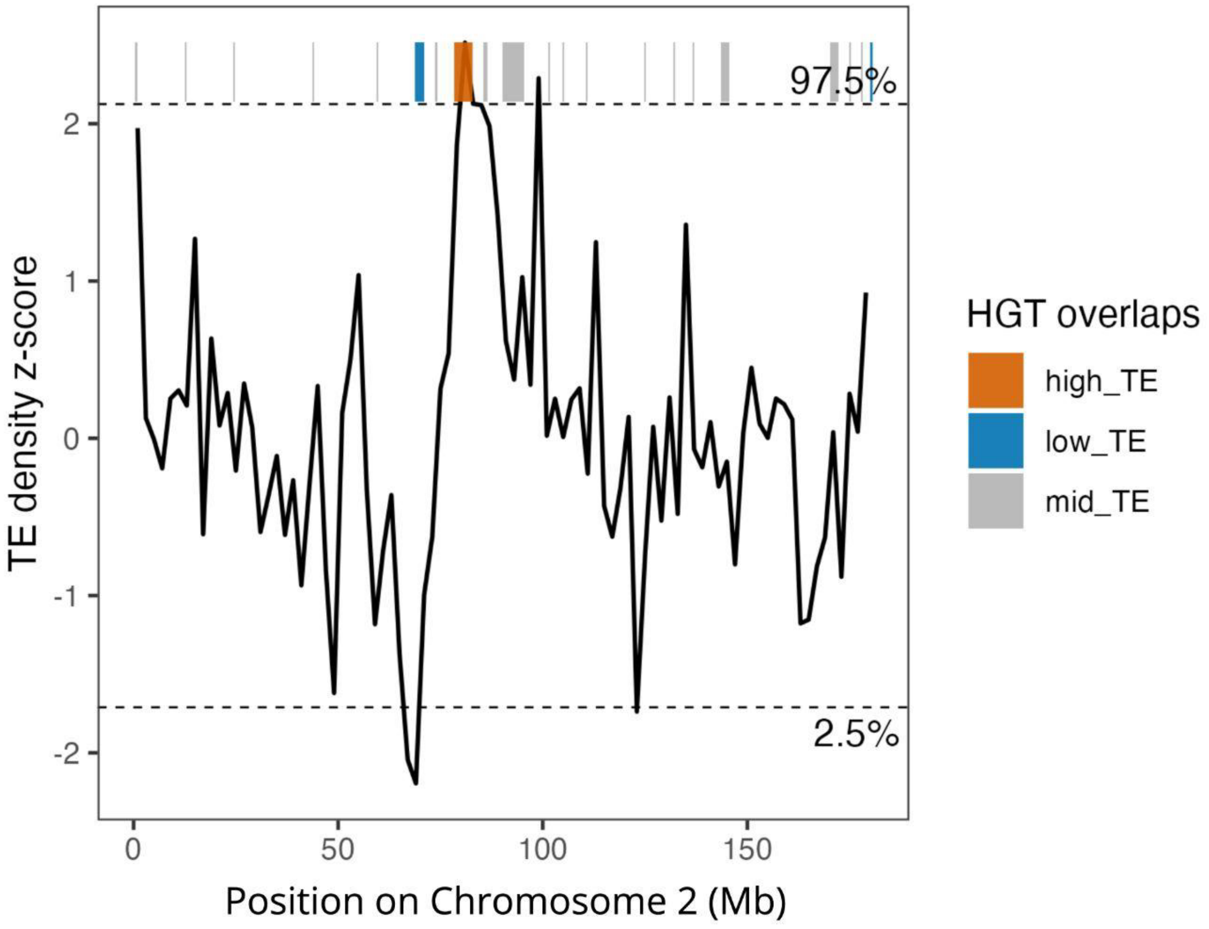


Supplementary Figure 15. Z-score transformed TE density by type in sliding 2 Mb windows across Chromosome 2 of *C. nanulus*. HGTs located within the upper (97.5th percentile) and lower (2.5th percentile) tails of the TE density distribution are indicated by orange and blue bars, respectively. Gray bars indicate HGTs falling outside these tails.

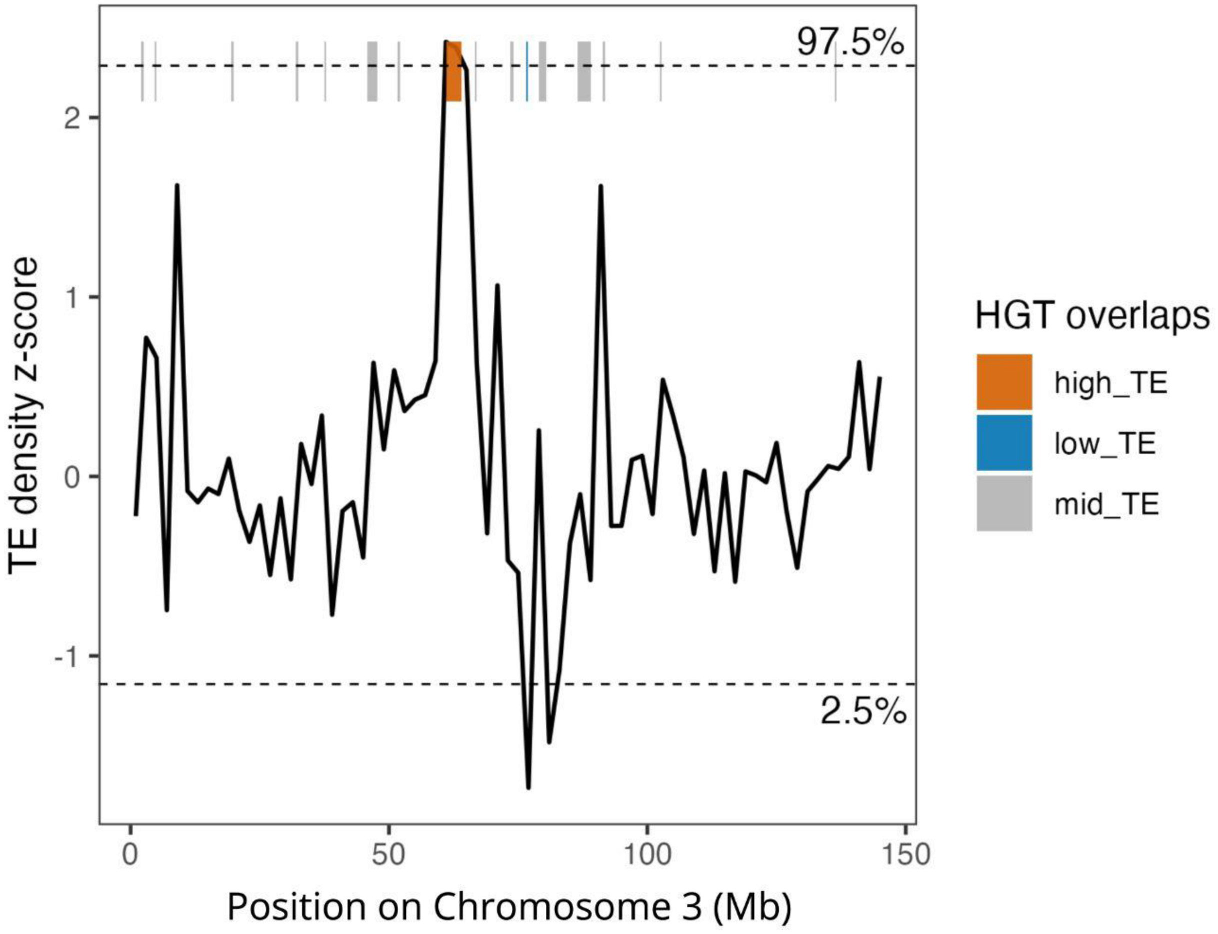


Supplementary Figure 16. Z-score transformed TE density by type in sliding 2 Mb windows across Chromosome 3 of *C. nanulus*. HGTs located within the upper (97.5th percentile) and lower (2.5th percentile) tails of the TE density distribution are indicated by orange and blue bars, respectively. Gray bars indicate HGTs falling outside these tails.

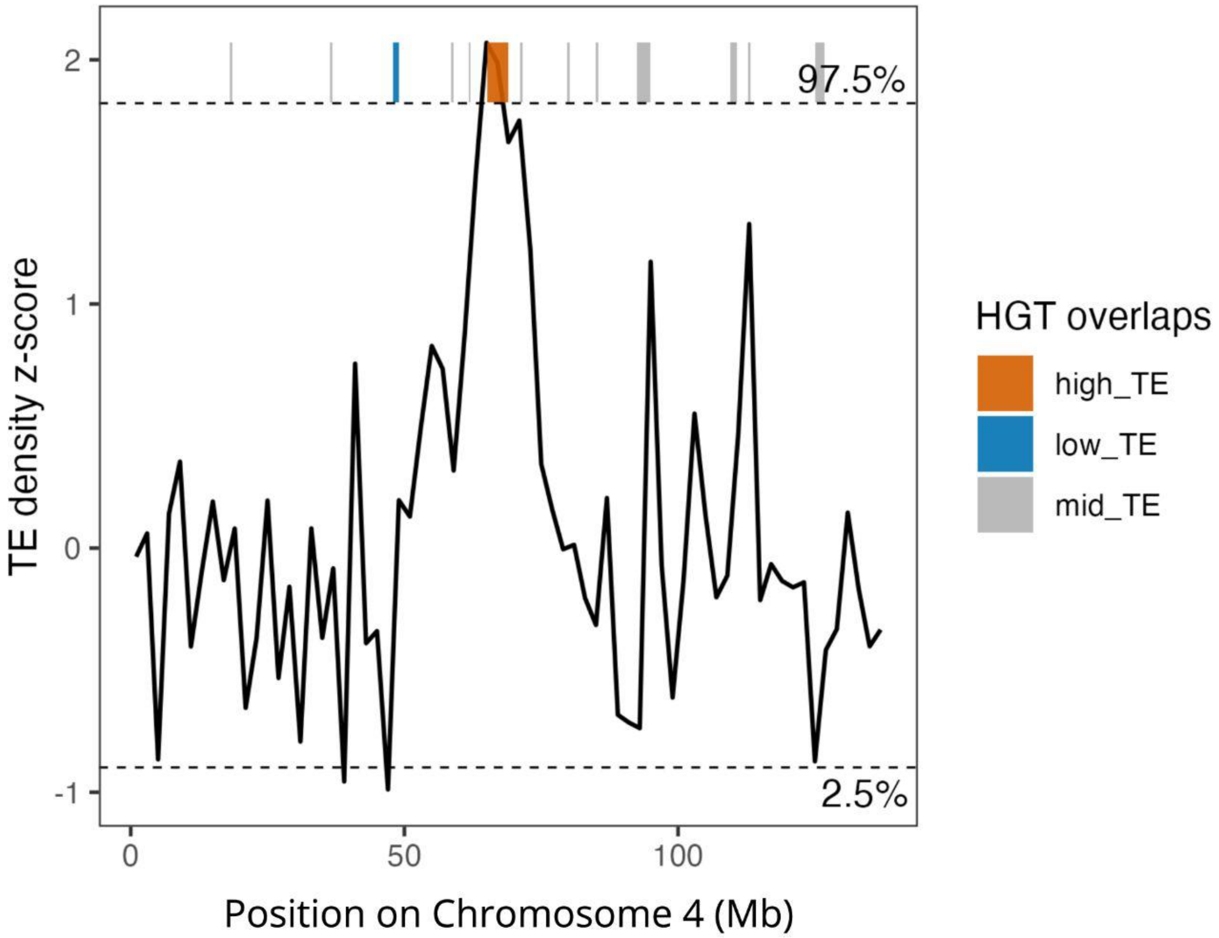


Supplementary Figure 17. Z-score transformed TE density by type in sliding 2 Mb windows across Chromosome 4 of *C. nanulus*. HGTs located within the upper (97.5th percentile) and lower (2.5th percentile) tails of the TE density distribution are indicated by orange and blue bars, respectively. Gray bars indicate HGTs falling outside these tails.

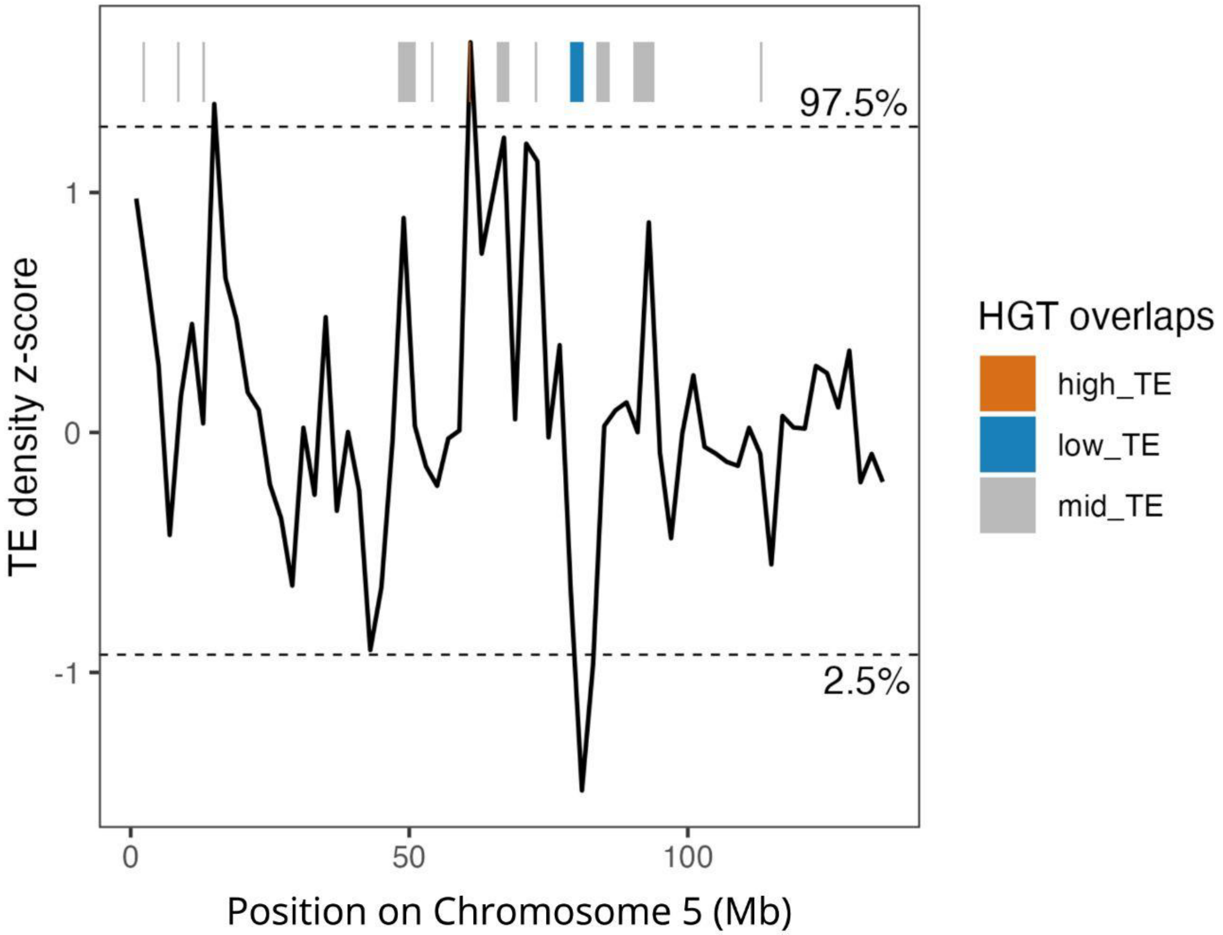


Supplementary Figure 18. Z-score transformed TE density by type in sliding 2 Mb windows across Chromosome 5 of *C. nanulus*. HGTs located within the upper (97.5th percentile) and lower (2.5th percentile) tails of the TE density distribution are indicated by orange and blue bars, respectively. Gray bars indicate HGTs falling outside these tails.

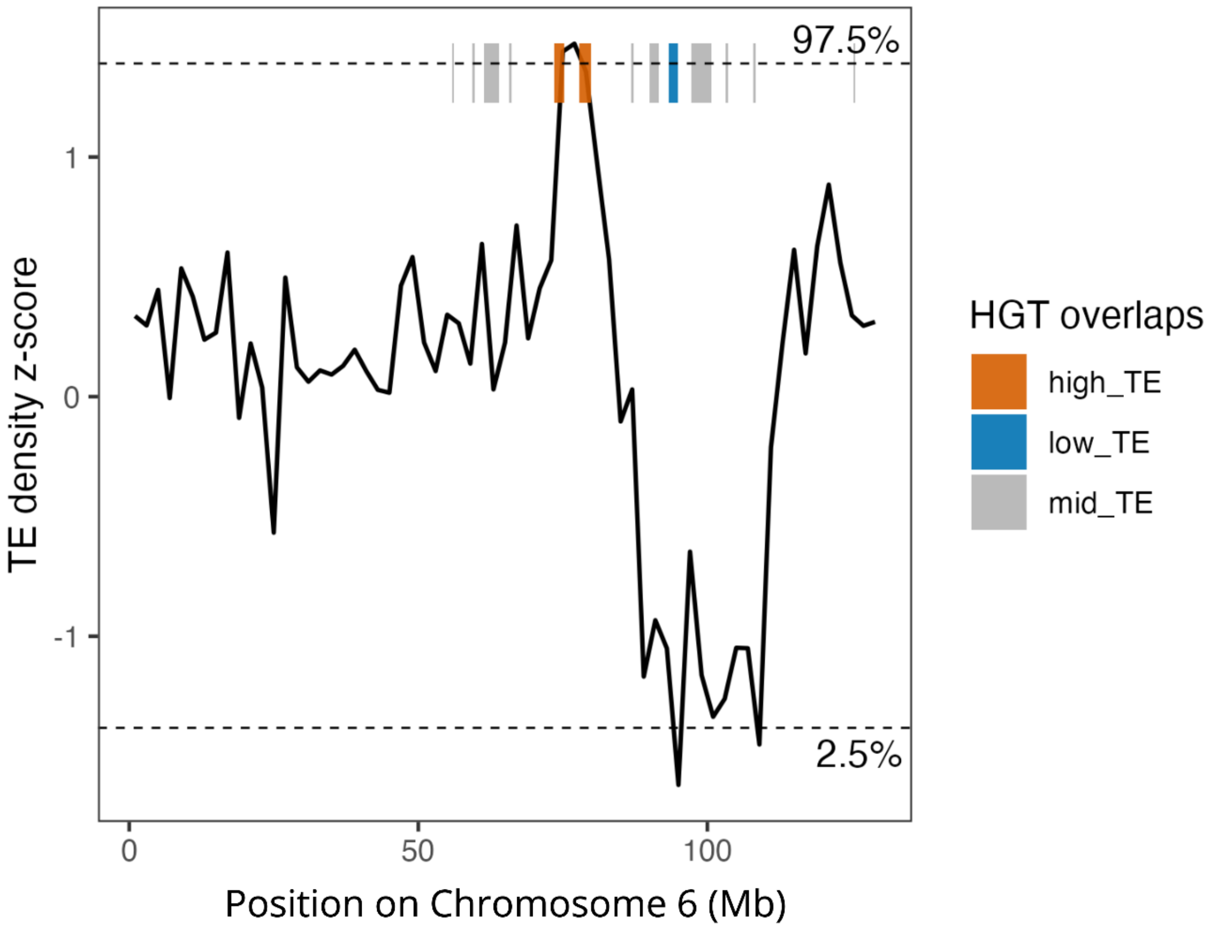


Supplementary Figure 19. Z-score transformed TE density by type in sliding 2 Mb windows across Chromosome 6 of *C. nanulus*. HGTs located within the upper (97.5th percentile) and lower (2.5th percentile) tails of the TE density distribution are indicated by orange and blue bars, respectively. Gray bars indicate HGTs falling outside these tails.

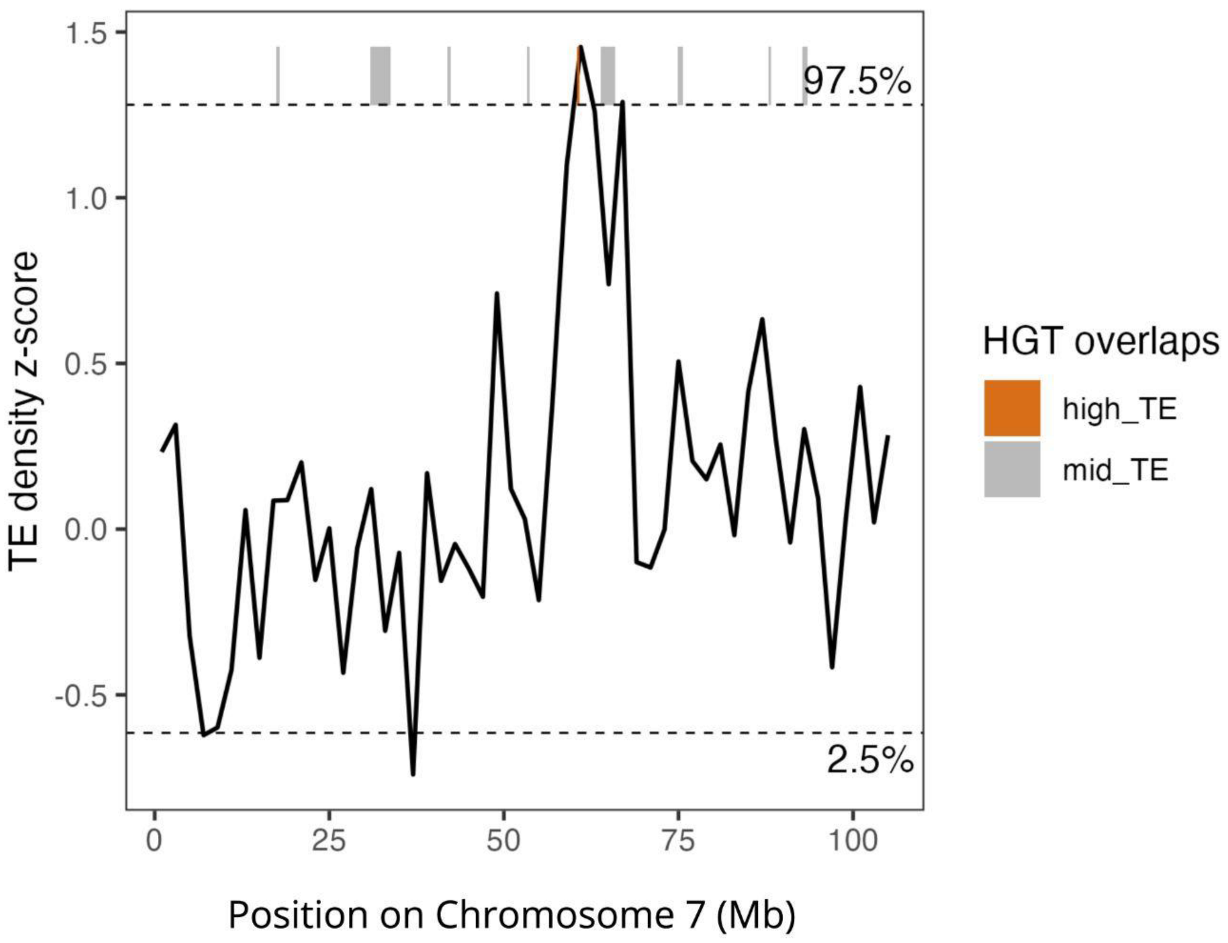


Supplementary Figure 20. Z-score transformed TE density by type in sliding 2 Mb windows across Chromosome 7 of *C. nanulus*. HGTs located within the upper (97.5th percentile) tail of the TE density distribution are indicated by orange bars. Gray bars indicate HGTs falling outside this tail.

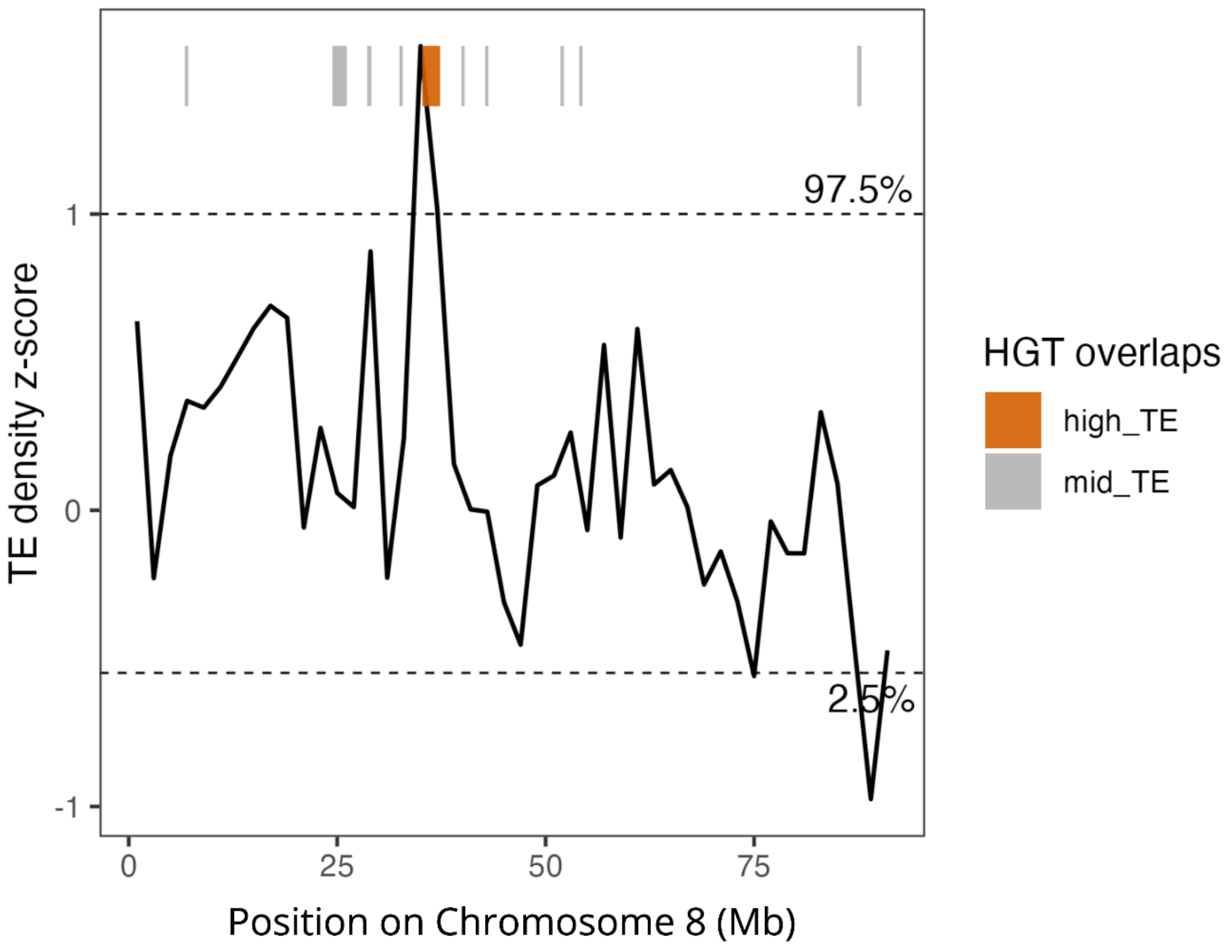


Supplementary Figure 21. Z-score transformed TE density by type in sliding 2 Mb windows across Chromosome 8 of *C. nanulus*. HGTs located within the upper (97.5th percentile) tail of the TE density distribution are indicated by orange bars. Gray bars indicate HGTs falling outside this tail.

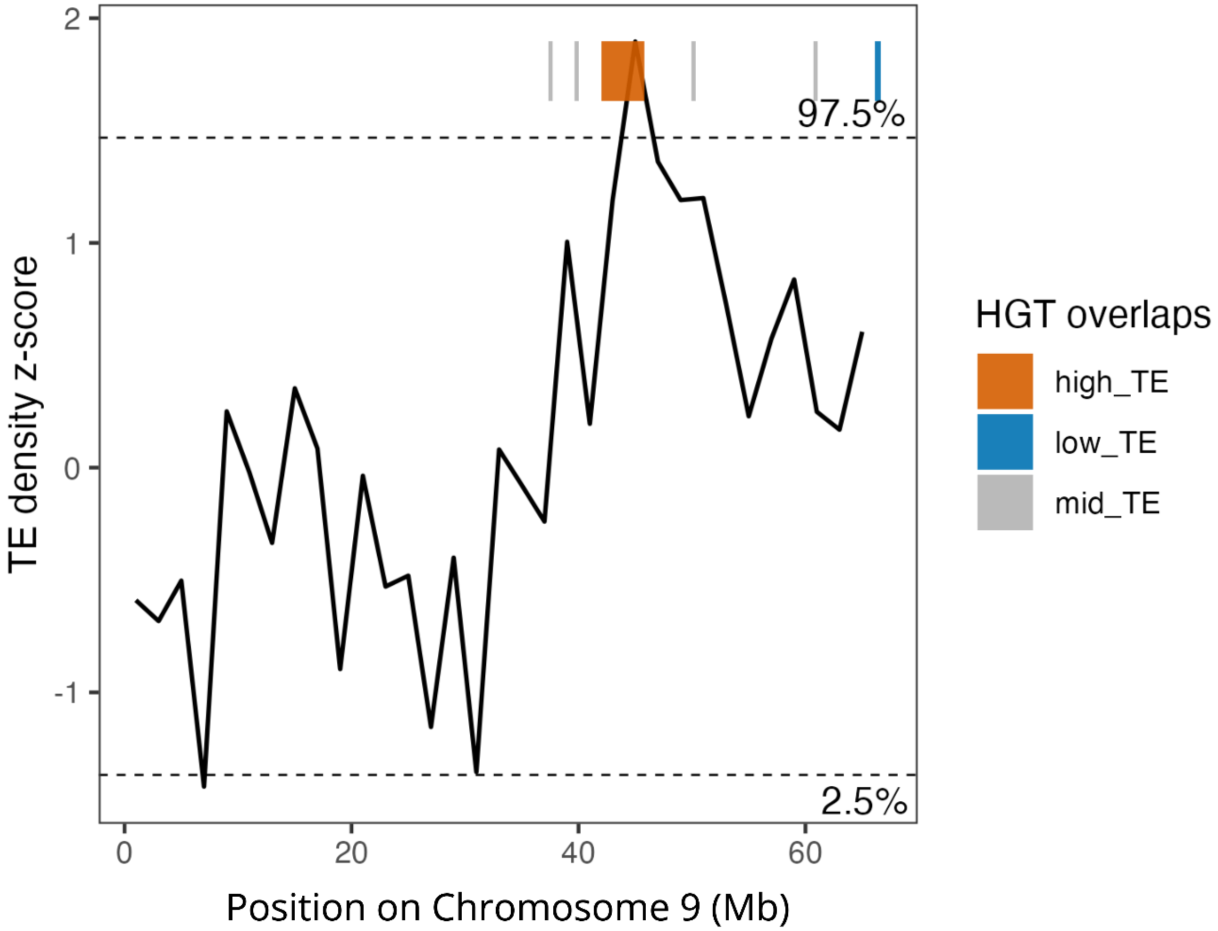


Supplementary Figure 22. Z-score transformed TE density by type in sliding 2 Mb windows across Chromosome 9 of *C. nanulus*. HGTs located within the upper (97.5th percentile) and lower (2.5th percentile) tails of the TE density distribution are indicated by orange and blue bars, respectively. Gray bars indicate HGTs falling outside these tails.

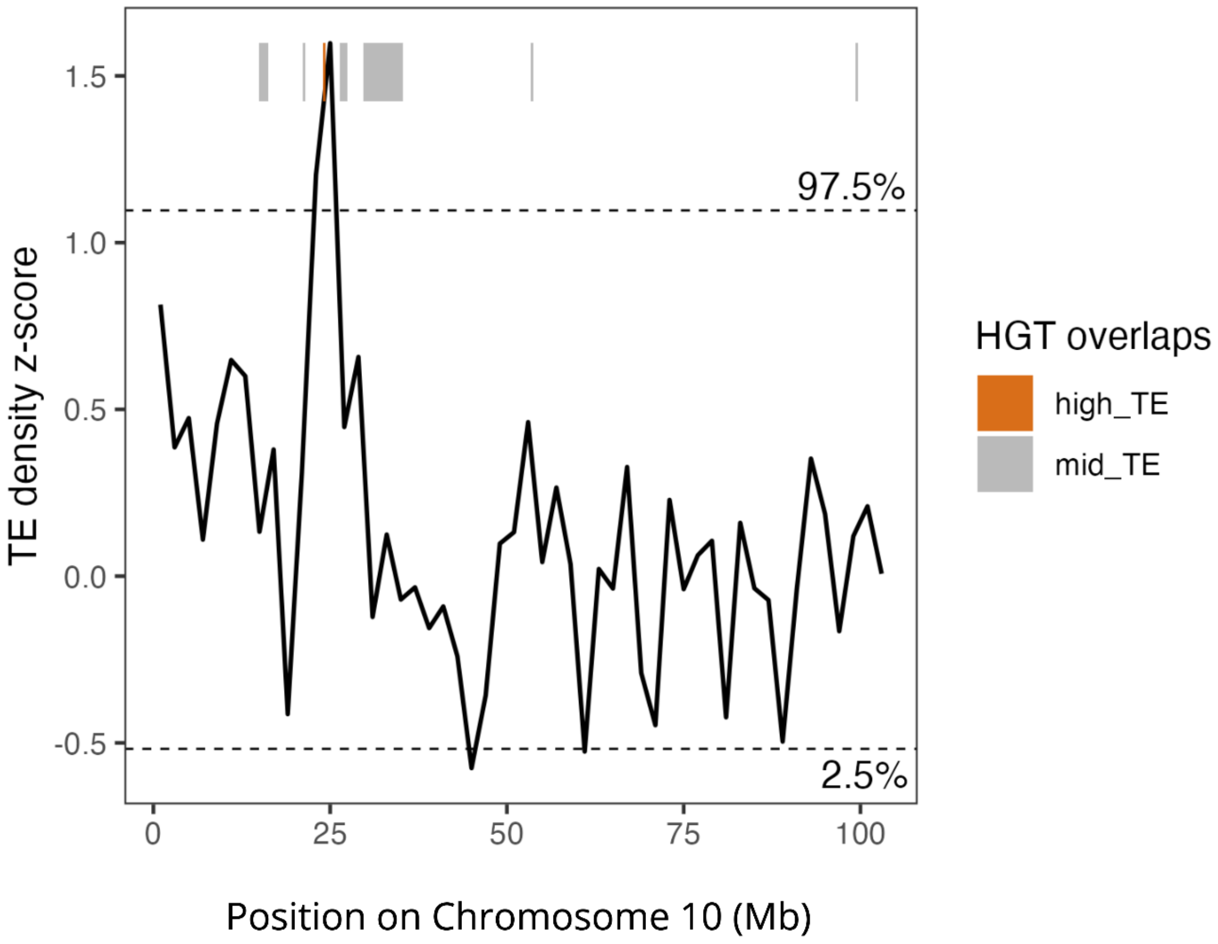


Supplementary Figure 23. Z-score transformed TE density by type in sliding 2 Mb windows across Chromosome 10 of *C. nanulus*. HGTs located within the upper (97.5th percentile) tail of the TE density distribution are indicated by orange bars. Gray bars indicate HGTs falling outside this tail.

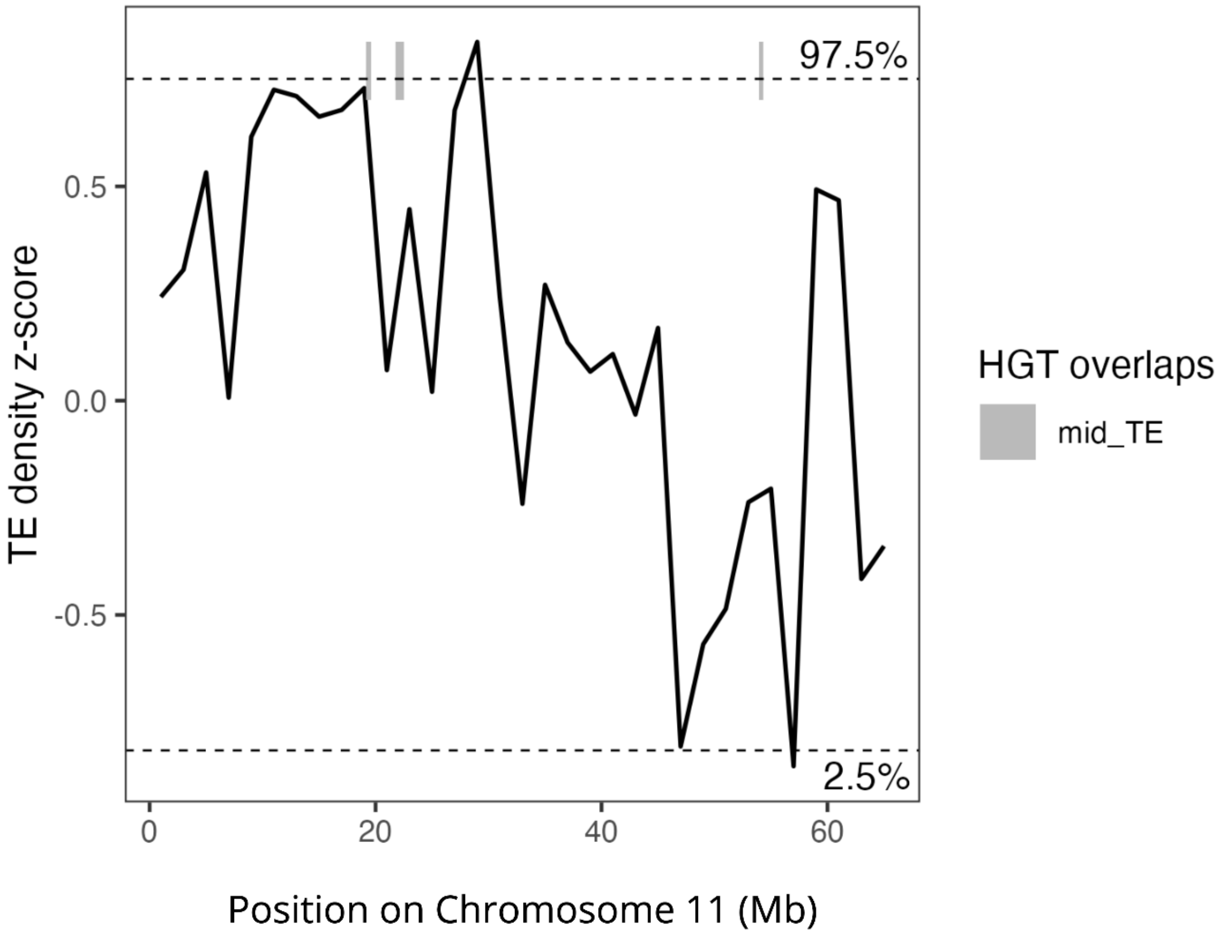


Supplementary Figure 24. Z-score transformed TE density by type in sliding 2 Mb windows across Chromosome 11 of *C. nanulus*. Horizontally transferred DNA sequences (HGTs) located outside of the tails of the TE density distribution are indicated with gray bars.

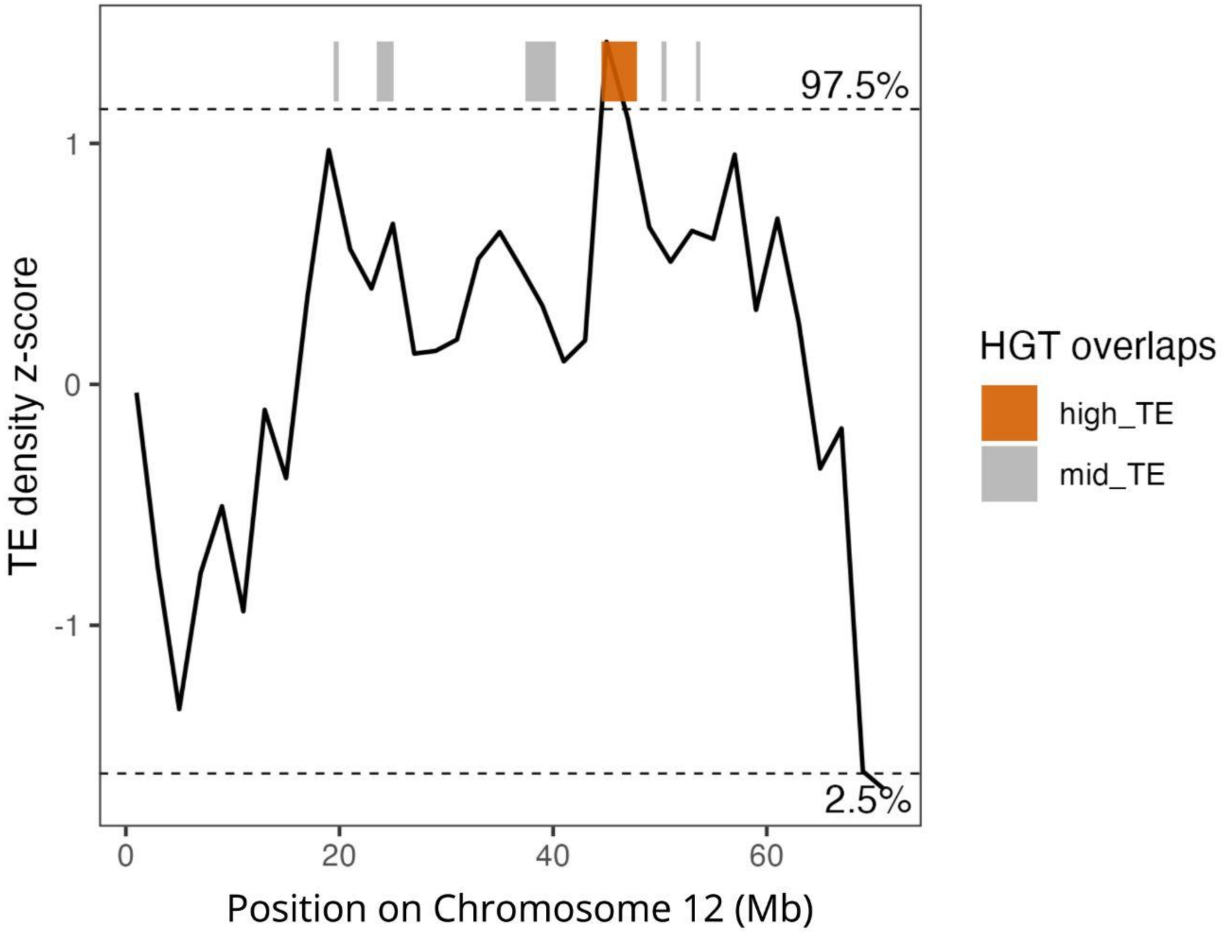


Supplementary Figure 25. Z-score transformed TE density by type in sliding 2 Mb windows across Chromosome 12 of *C. nanulus*. HGTs located within the upper (97.5th percentile) tail of the TE density distribution are indicated by orange bars. Gray bars indicate HGTs falling outside this tail.

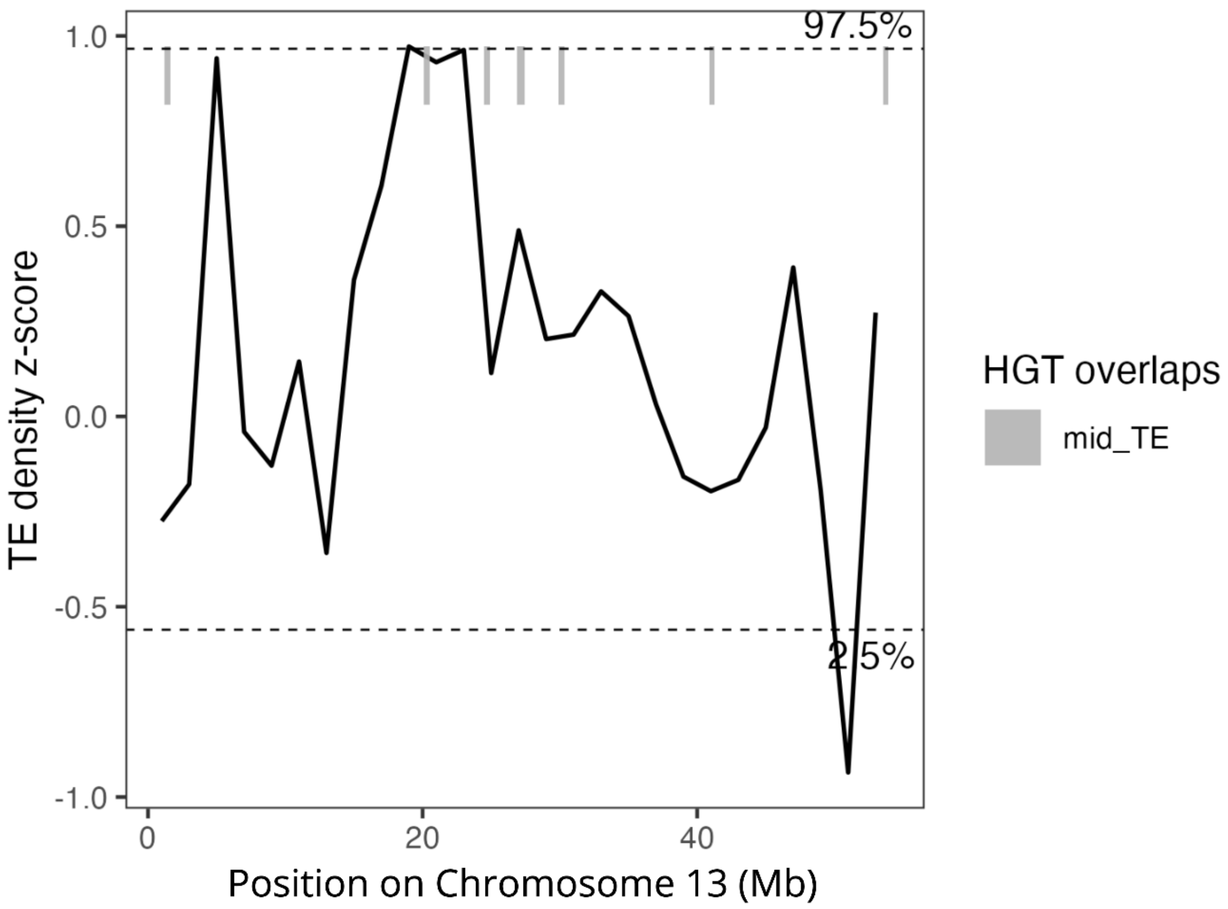


Supplementary Figure 26. Z-score transformed TE density by type in sliding 2 Mb windows across Chromosome 13 of *C. nanulus*. Horizontally transferred DNA sequences (HGTs) located outside of the tails of the TE density distribution are indicated with gray bars.

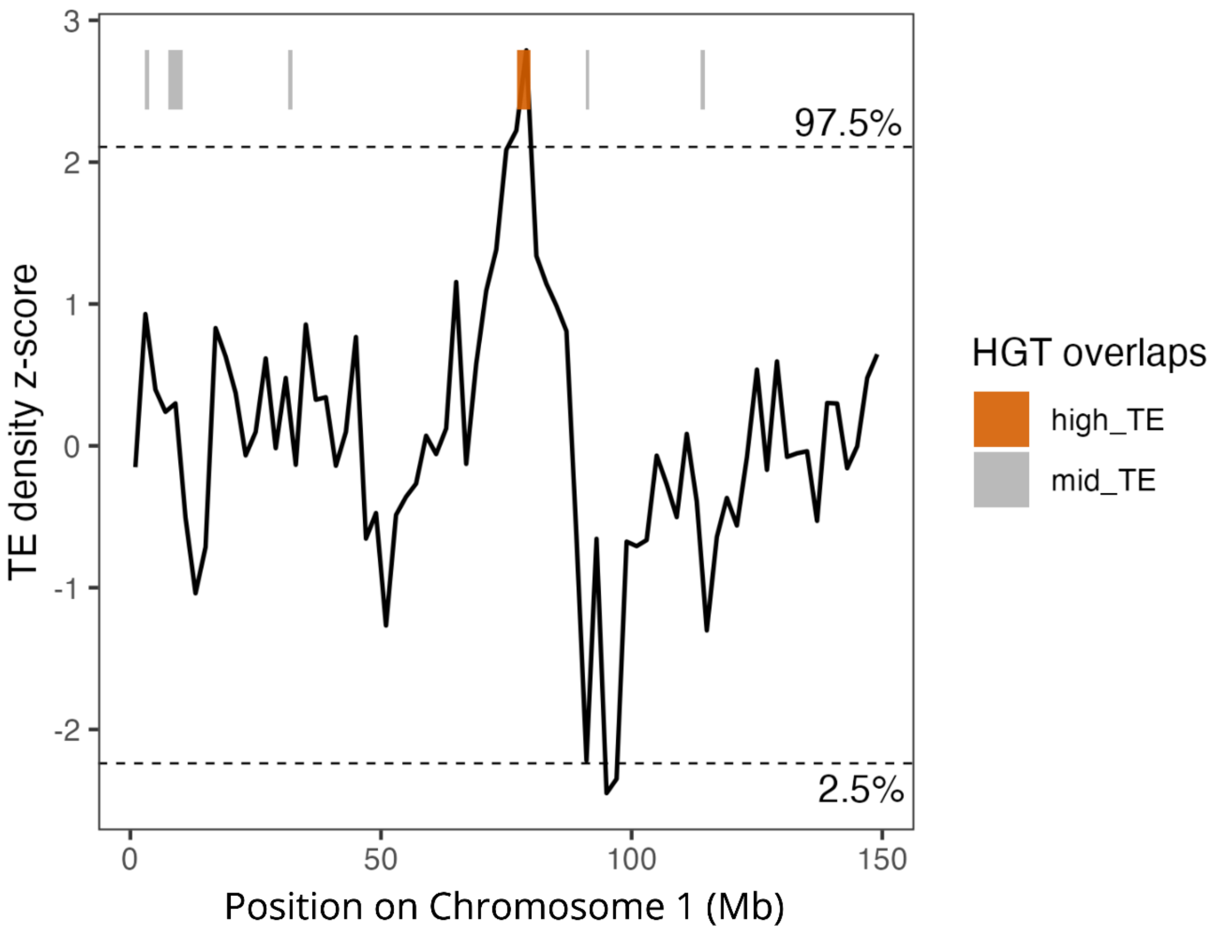


Supplementary Figure 27. Z-score transformed TE density by type in sliding 2 Mb windows across Chromosome 1 of *C. glandium*. HGTs located within the upper (97.5th percentile) tail of the TE density distribution are indicated by orange bars. Gray bars indicate HGTs falling outside this tail.

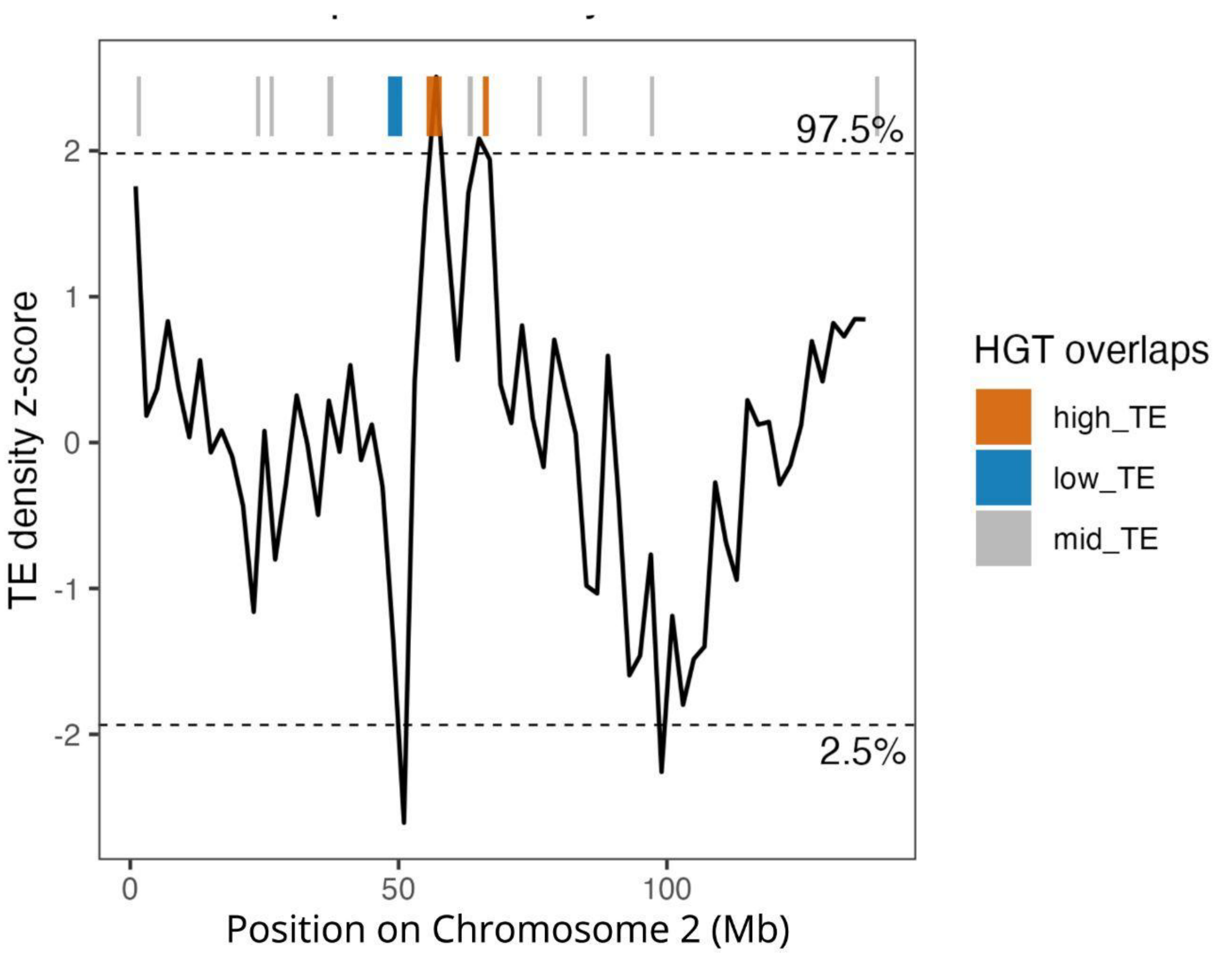


Supplementary Figure 28. Z-score transformed TE density by type in sliding 2 Mb windows across Chromosome 2 of *C. glandium*. HGTs located within the upper (97.5th percentile) and lower (2.5th percentile) tails of the TE density distribution are indicated by orange and blue bars, respectively. Gray bars indicate HGTs falling outside these tails.

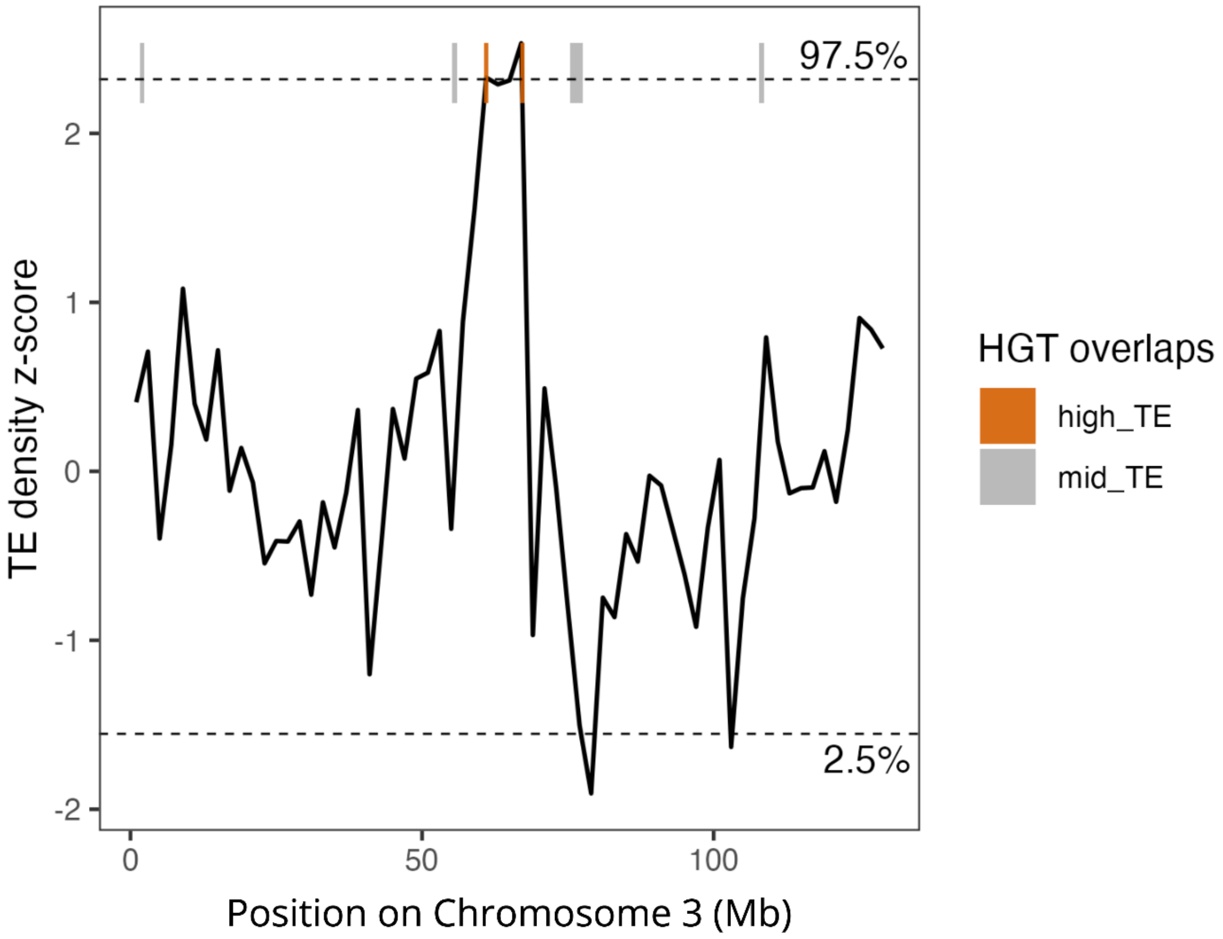


Supplementary Figure 29. Z-score transformed TE density by type in sliding 2 Mb windows across Chromosome 3 of *C. glandium*. HGTs located within the upper (97.5th percentile) tail of the TE density distribution are indicated by orange bars. Gray bars indicate HGTs falling outside this tail.

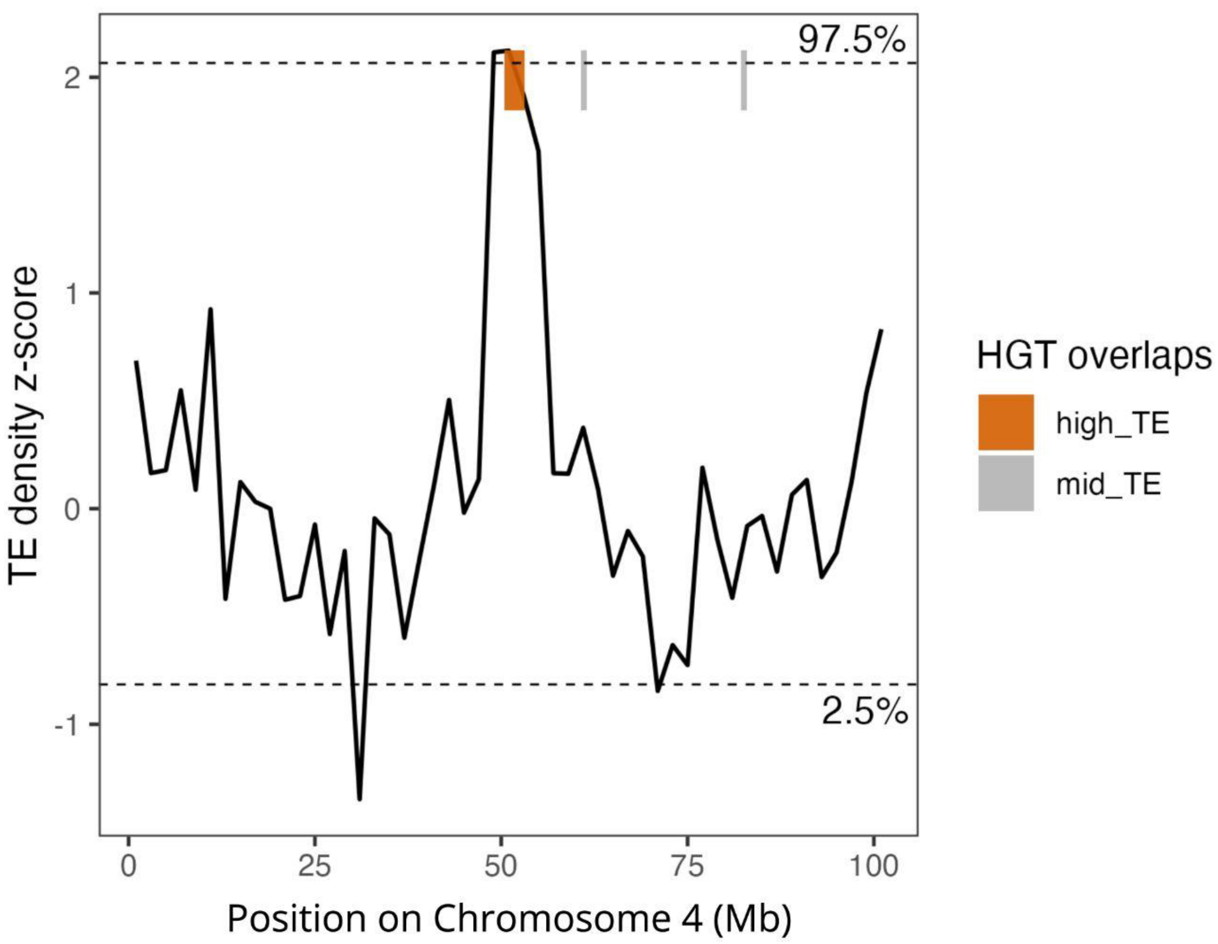


Supplementary Figure 30. Z-score transformed TE density by type in sliding 2 Mb windows across Chromosome 4 of *C. glandium*. HGTs located within the upper (97.5th percentile) tail of the TE density distribution are indicated by orange bars. Gray bars indicate HGTs falling outside this tail.

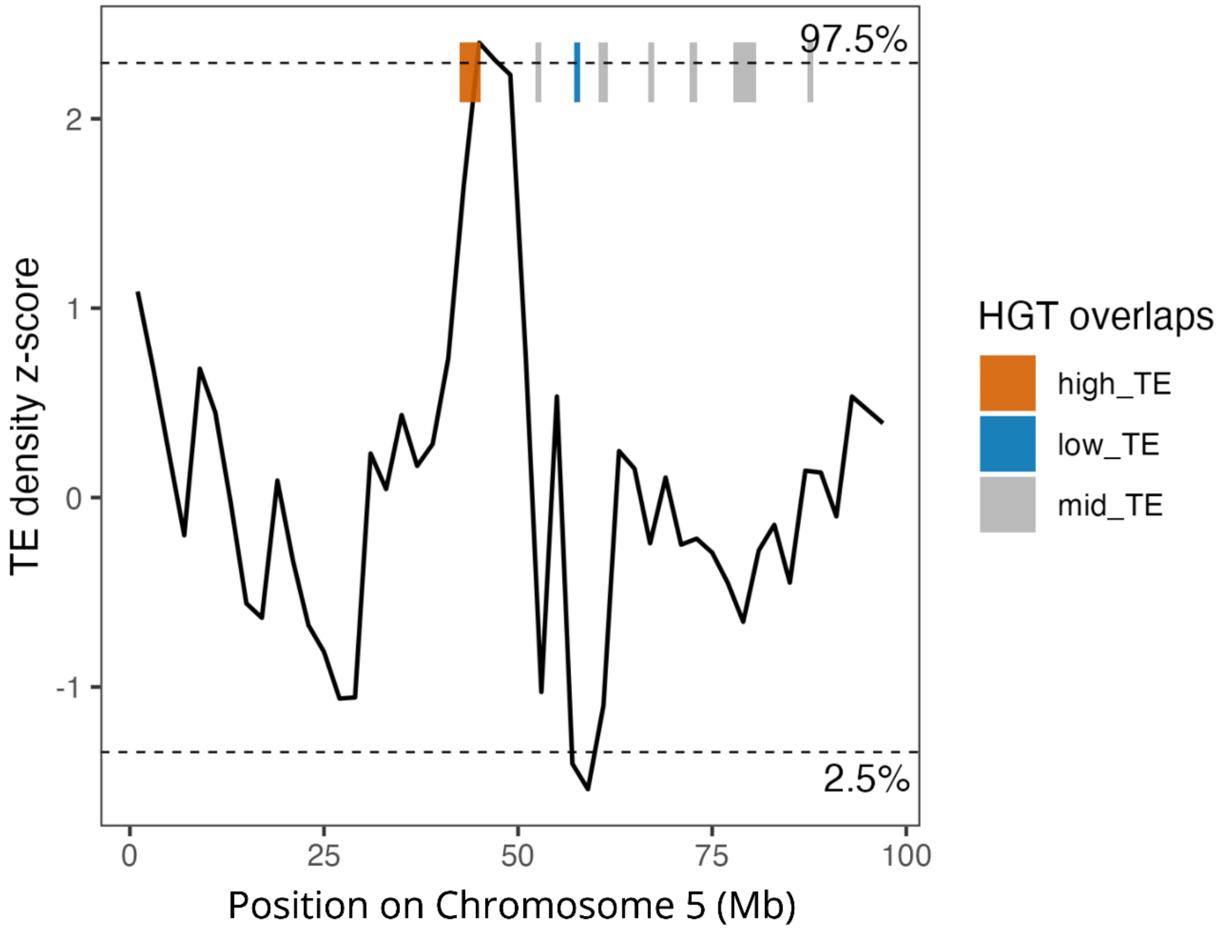


Supplementary Figure 31. Z-score transformed TE density by type in sliding 2 Mb windows across Chromosome 5 of *C. glandium*. HGTs located within the upper (97.5th percentile) and lower (2.5th percentile) tails of the TE density distribution are indicated by orange and blue bars, respectively. Gray bars indicate HGTs falling outside these tails.

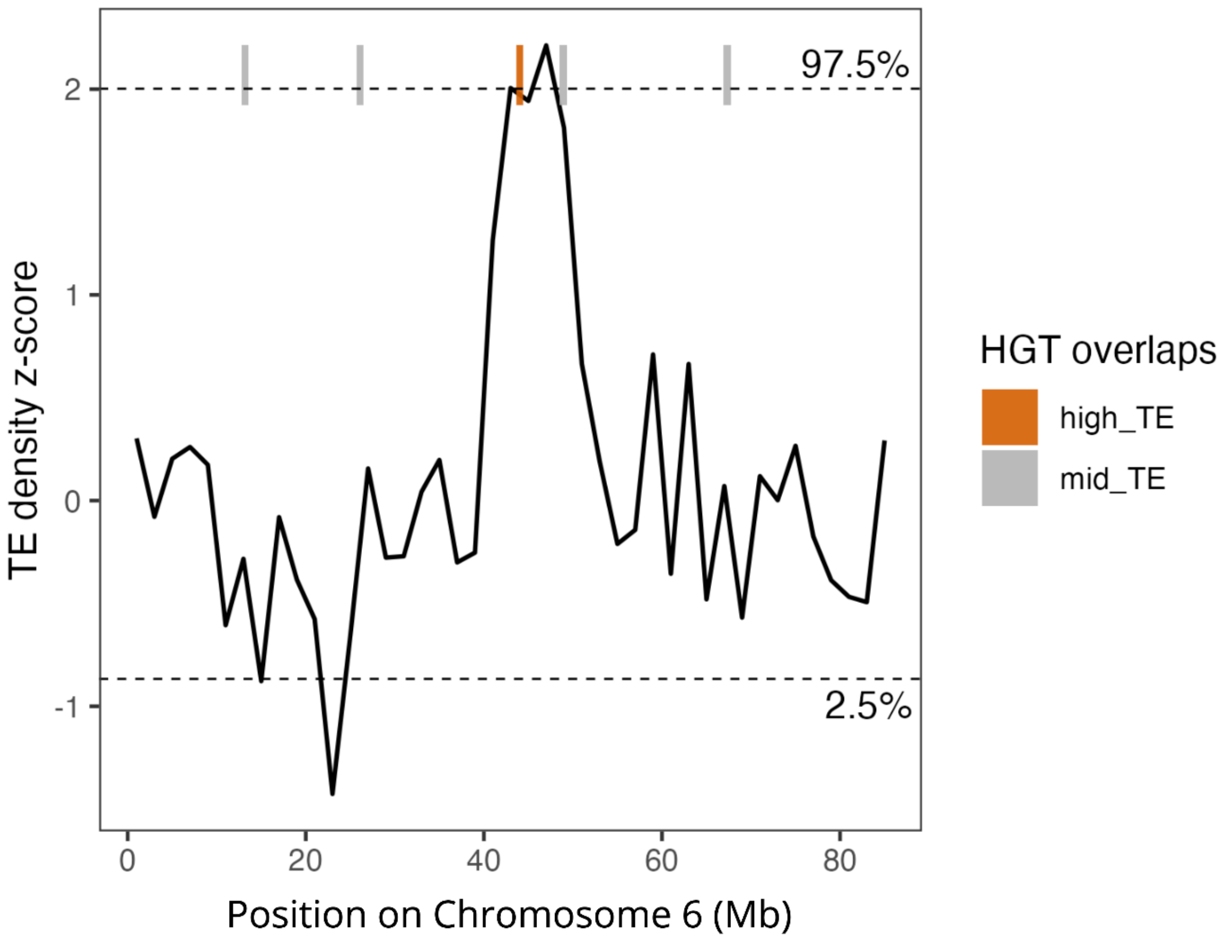


Supplementary Figure 32. Z-score transformed TE density by type in sliding 2 Mb windows across Chromosome 6 of *C. glandium*. HGTs located within the upper (97.5th percentile) tail of the TE density distribution are indicated by orange bars. Gray bars indicate HGTs falling outside this tail.

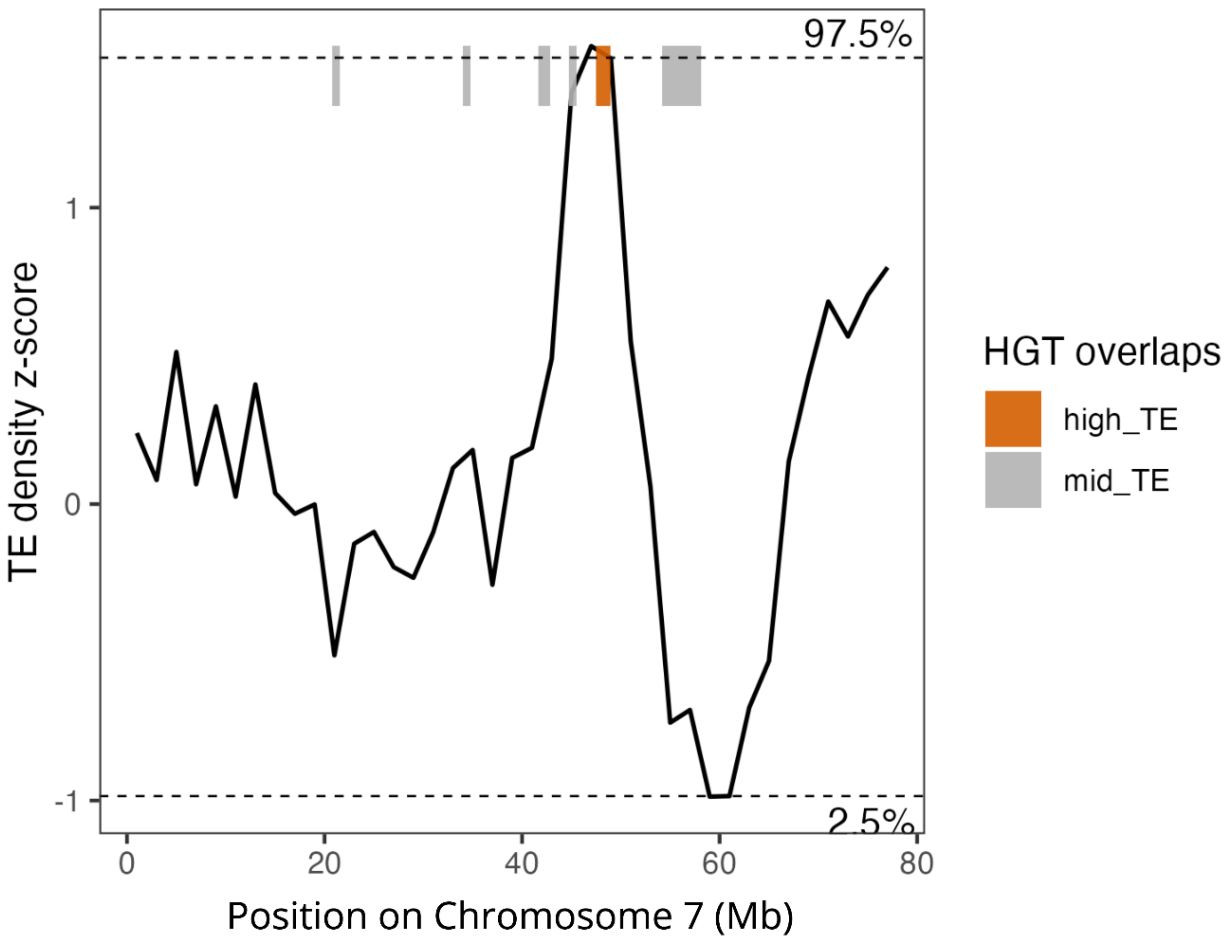


Supplementary Figure 33. Z-score transformed TE density by type in sliding 2 Mb windows across Chromosome 7 of *C. glandium*. HGTs located within the upper (97.5th percentile) tail of the TE density distribution are indicated by orange bars. Gray bars indicate HGTs falling outside this tail.

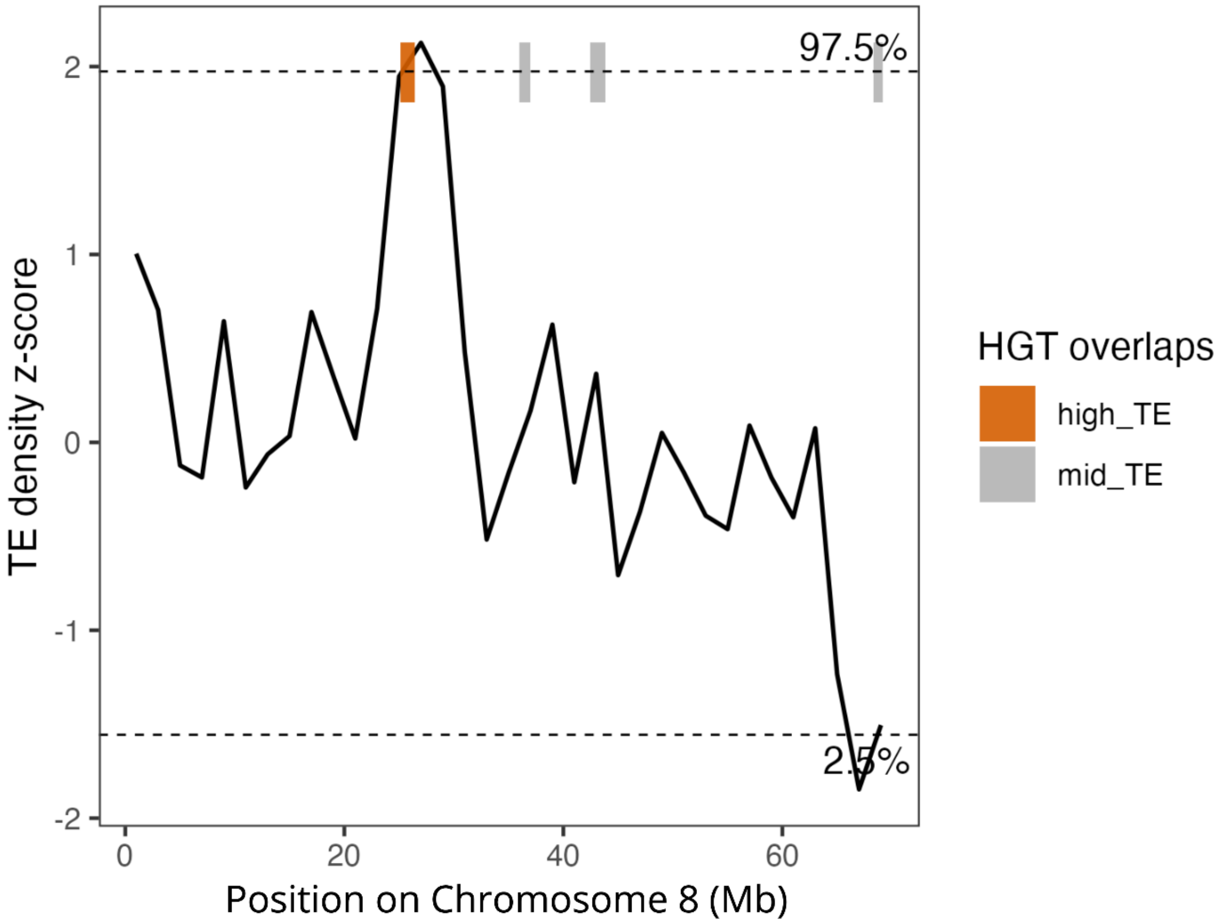


Supplementary Figure 34. Z-score transformed TE density by type in sliding 2 Mb windows across Chromosome 8 of *C. glandium*. HGTs located within the upper (97.5th percentile) tail of the TE density distribution are indicated by orange bars. Gray bars indicate HGTs falling outside this tail.

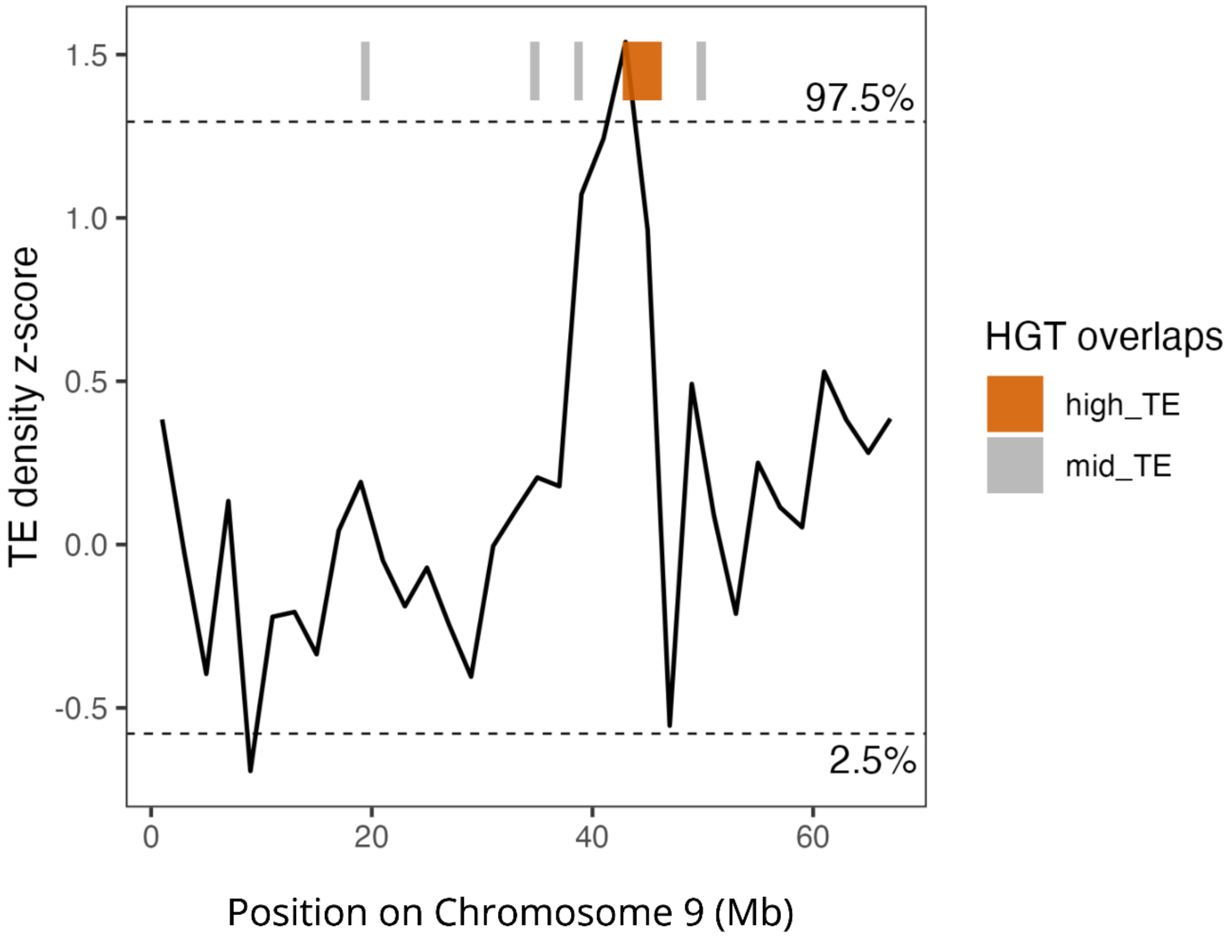


Supplementary Figure 35. Z-score transformed TE density by type in sliding 2 Mb windows across Chromosome 9 of *C. glandium*. HGTs located within the upper (97.5th percentile) tail of the TE density distribution are indicated by orange bars. Gray bars indicate HGTs falling outside this tail.

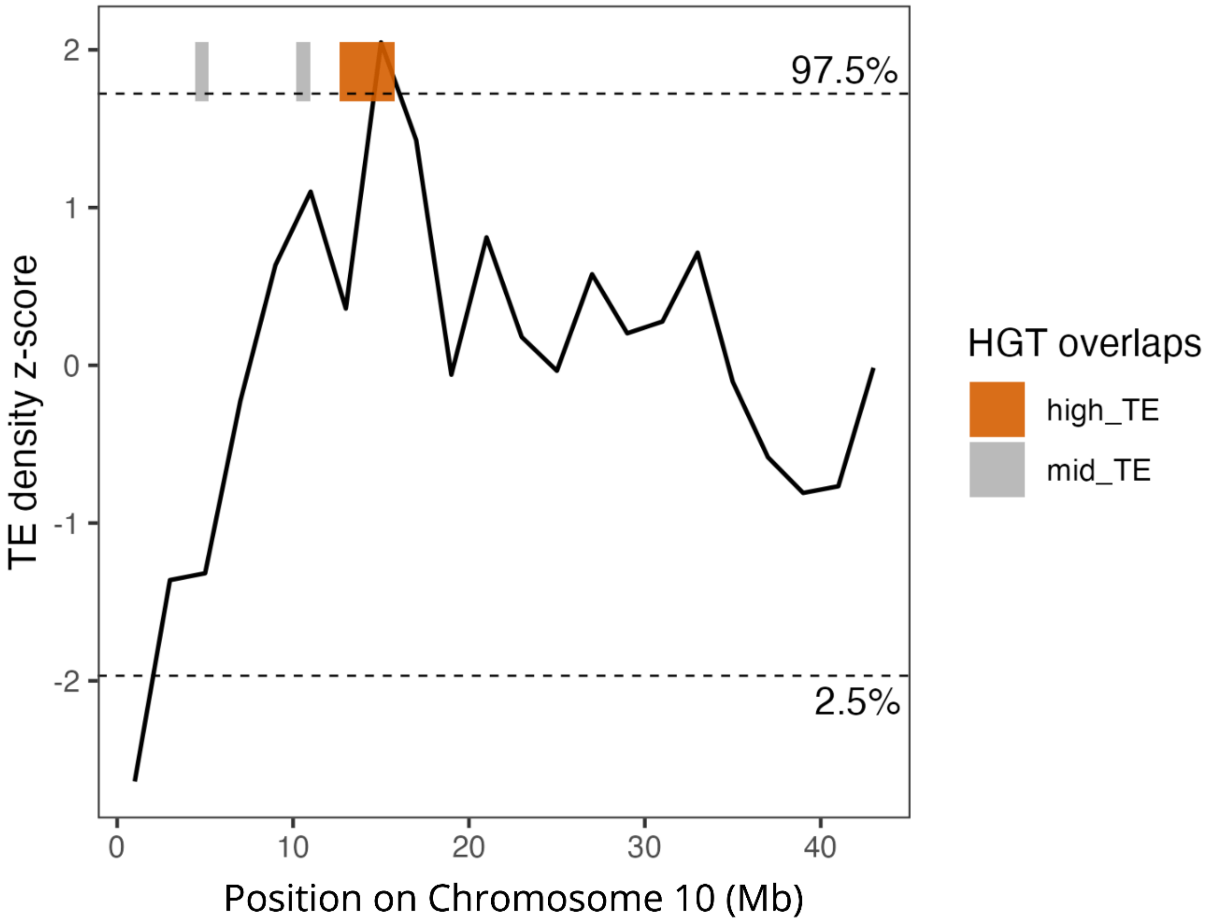


Supplementary Figure 36. Z-score transformed TE density by type in sliding 2 Mb windows across Chromosome 10 of *C. glandium*. HGTs located within the upper (97.5th percentile) tail of the TE density distribution are indicated by orange bars. Gray bars indicate HGTs falling outside this tail.

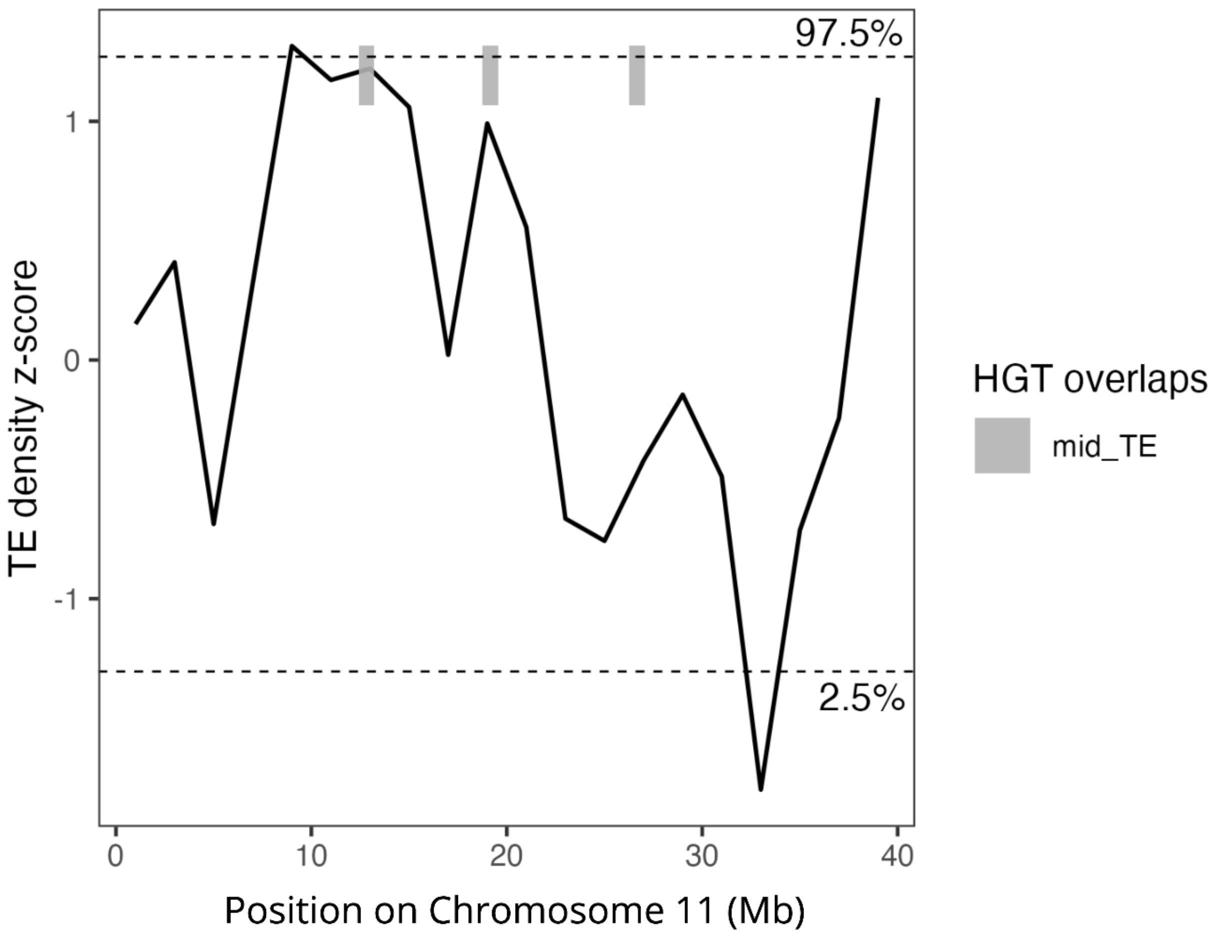


Supplementary Figure 37. Z-score transformed TE density by type in sliding 2 Mb windows across Chromosome 11 of *C. glandium*. Horizontally transferred DNA sequences (HGTs) located outside of the tails of the TE density distribution are indicated with gray bars.

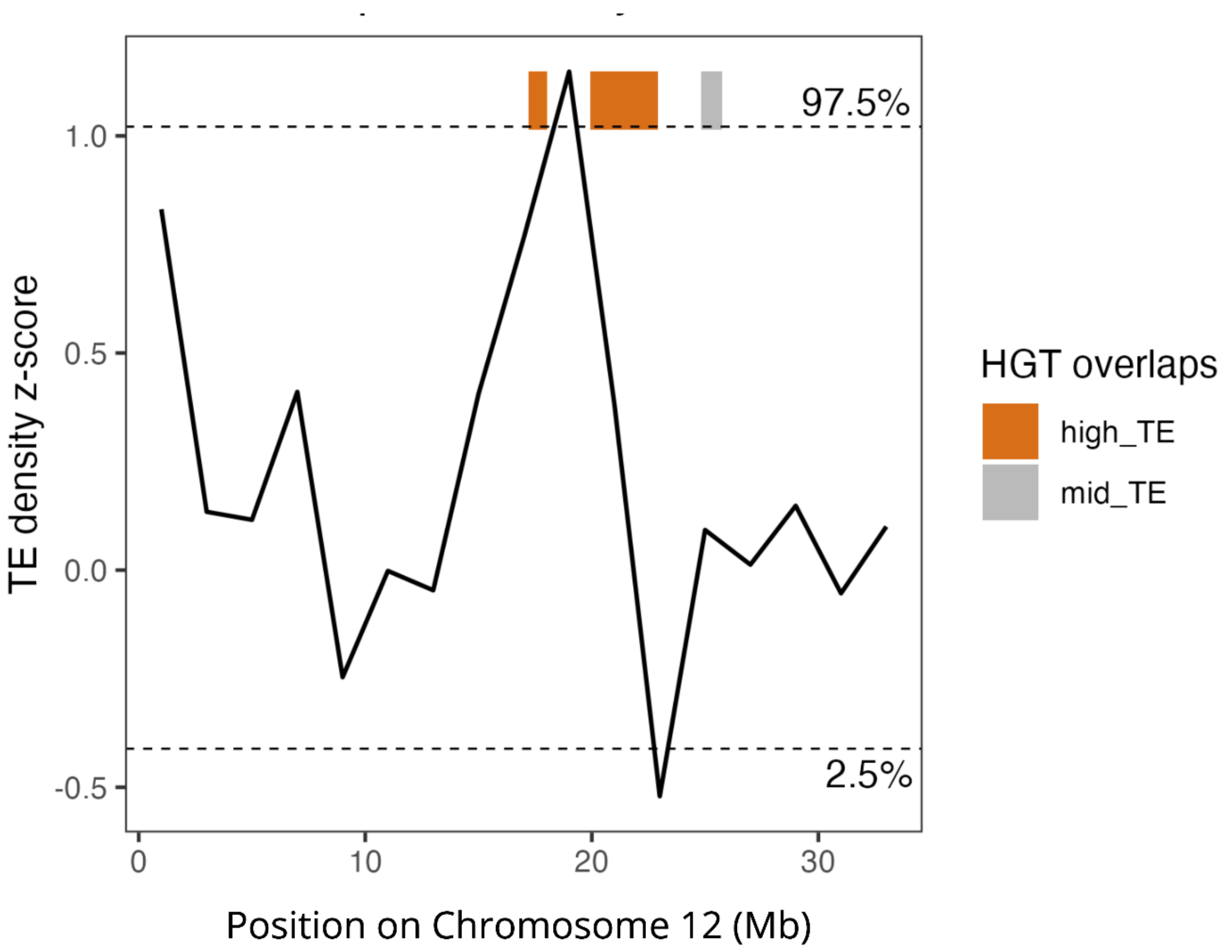


Supplementary Figure 38. Z-score transformed TE density by type in sliding 2 Mb windows across Chromosome 12 of *C. glandium*. HGTs located within the upper (97.5th percentile) tail of the TE density distribution are indicated by orange bars. Gray bars indicate HGTs falling outside this tail.

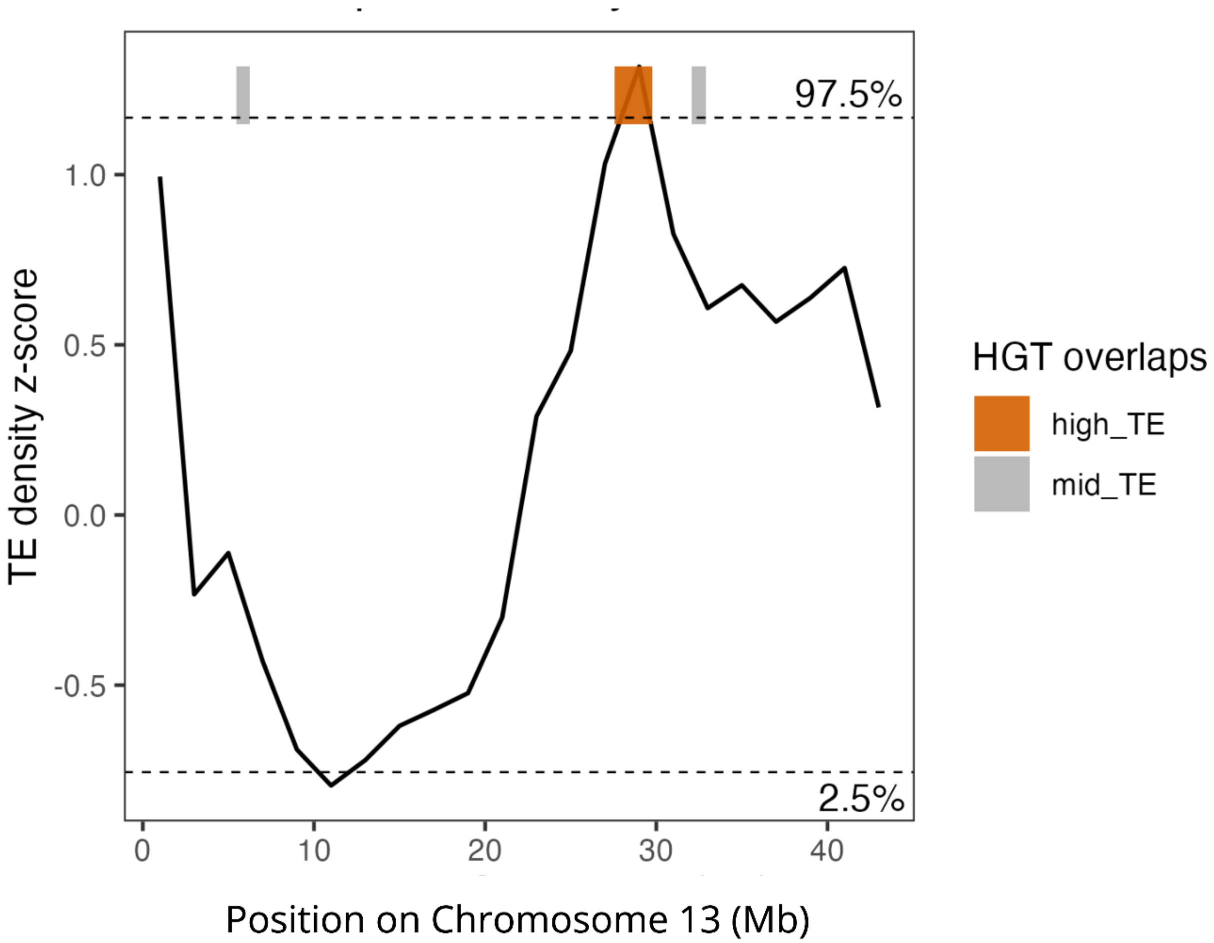


Supplementary Figure 39. Z-score transformed TE density by type in sliding 2 Mb windows across Chromosome 13 of *C. glandium*. HGTs located within the upper (97.5th percentile) tail of the TE density distribution are indicated by orange bars. Gray bars indicate HGTs falling outside this tail.

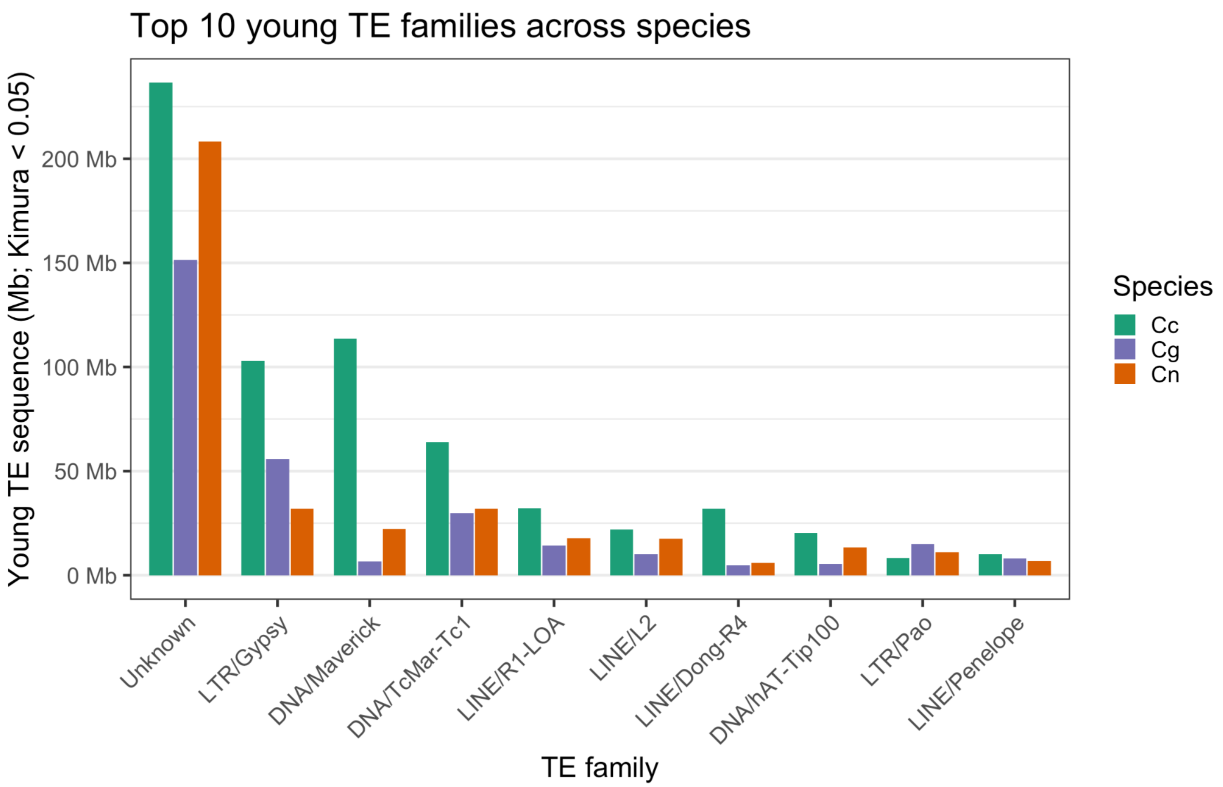


Supplementary Figure 40. Top 10 young (Kimura $D < 0.05$) genome-wide TE families across *Curculio* species: *C. caryae* (Cc, teal), *C. glandium* (Cg, purple), and *C. nanulus* (Cn, orange).

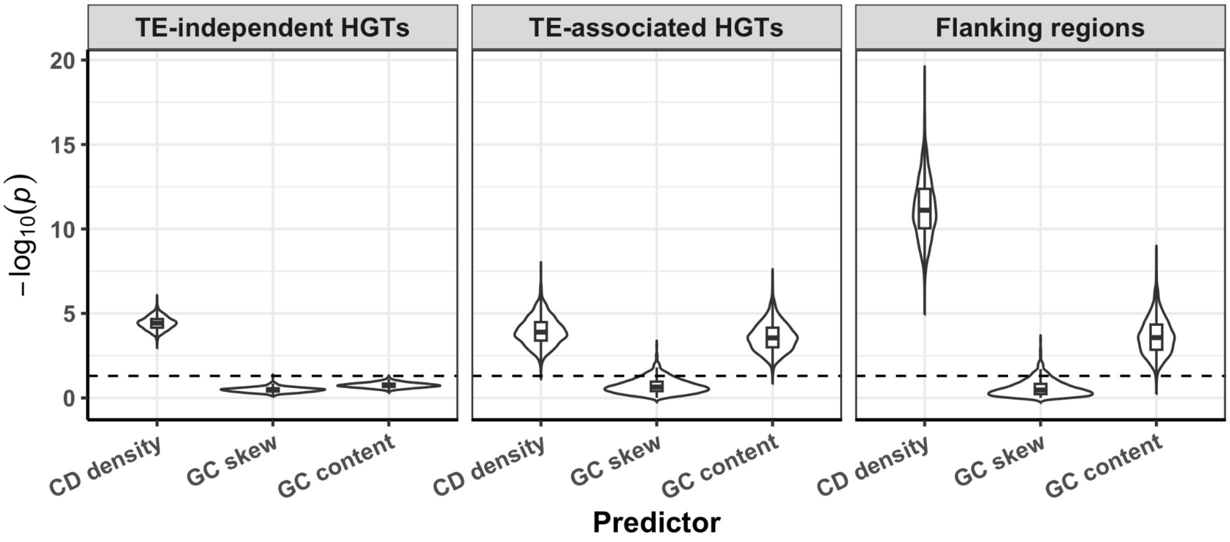


Supplementary Figure 41. Violin plots with embedded box plots summarizing the distributions of *z*-test $-\log_{10}$(*p*-values) for testing the null hypothesis that the regression coefficient $\beta_j$ for a given predictor *j* is zero ($H_0$: $\beta_j = 0$), with the alternative hypothesis that $\beta_j$ is not equal to zero ($H_0$: $\beta_j \neq 0$). The distribution summaries are based on 500 chromosome-matched multinomial regression replicates. Each model had three predictors: coding sequence (CD) density, GC skew, and GC content. The models predicted four response classes: TE-independent HGT cores, TE-associated HGT cores, HGT flanking windows, and chromosome-matched background windows as the reference class. The horizontal dashed lines denote the significance threshold of $-\log_{10}(0.05)$.

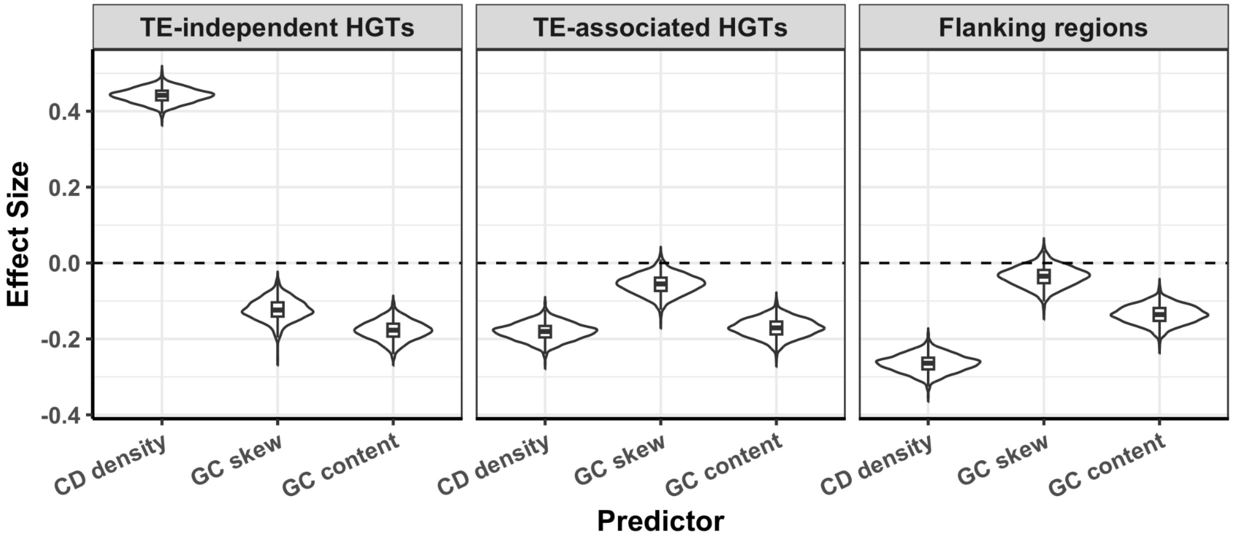


Supplementary Figure 42. Violin plots with embedded box plots summarizing the distributions of predictor effect sizes (regression coefficient values) across 500 chromosome-matched multinomial regression replicates. Each model had three predictors: coding sequence (CD) density, GC skew, and GC content. The models predicted four response classes: TE-associated HGT cores, TE-independent HGT cores, HGT flanking windows, and chromosome-matched background windows as the reference class. Positive coefficients indicate increased log odds of assignment to that response class for windows with higher values of the corresponding predictor, whereas negative coefficients indicate decreased log odds, relative to background. The horizontal dashed lines denote a regression coefficient of zero, indicating no association between the predictor and the response class.

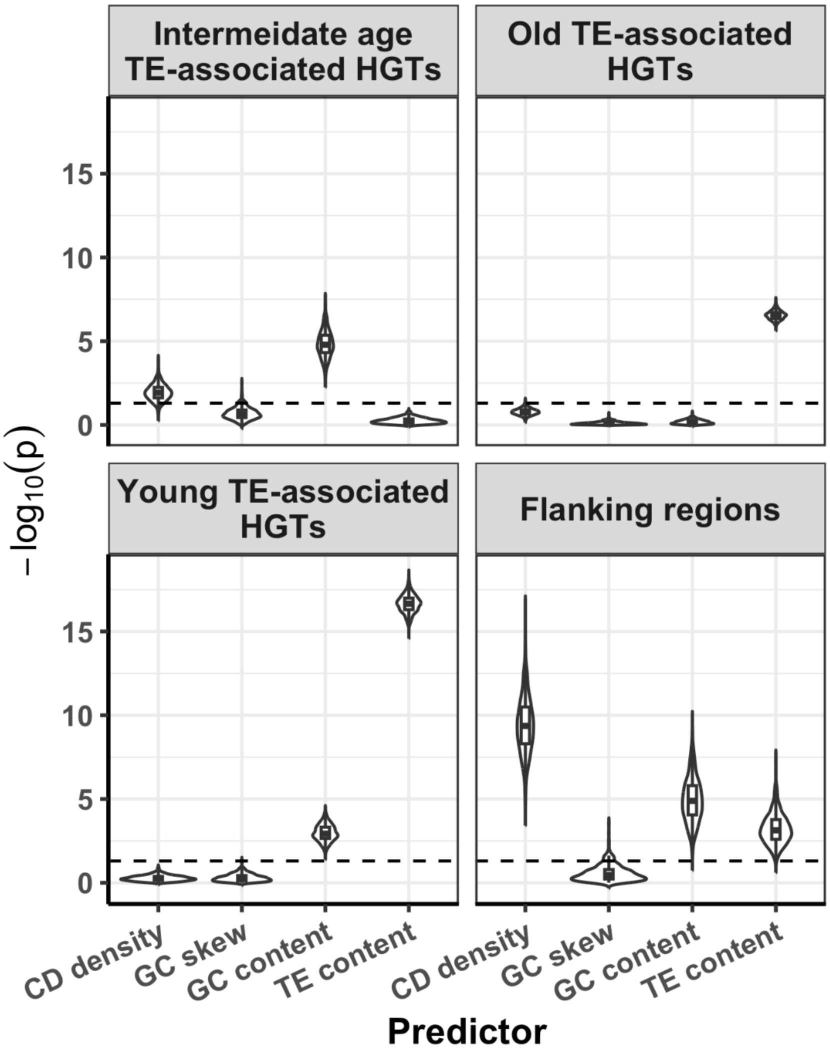


Supplementary Figure 43. Violin plots with embedded box plots summarizing the distributions of *z*-test -$\log_{10}$ (*p*-values) for testing the null hypothesis that the regression coefficient $\beta_j$ for a given predictor *j* is zero ($H_0$: $\beta_j = 0$), with the alternative hypothesis that $\beta_j$ is not equal to zero ($H_0$: $\beta_j \neq 0$). The distribution summaries are based on 500 chromosome-matched multinomial regression replicates. Each model had four predictors: coding sequence (CD) density, GC skew, GC content, and TE content. The models predicted five response classes: TE-associated HGT cores based on TE age as young, intermediate, and old, HGT flanking windows, and chromosome-matched background windows as the reference class. The horizontal dashed lines denote the significance threshold of -$\log_{10}$(0.05).

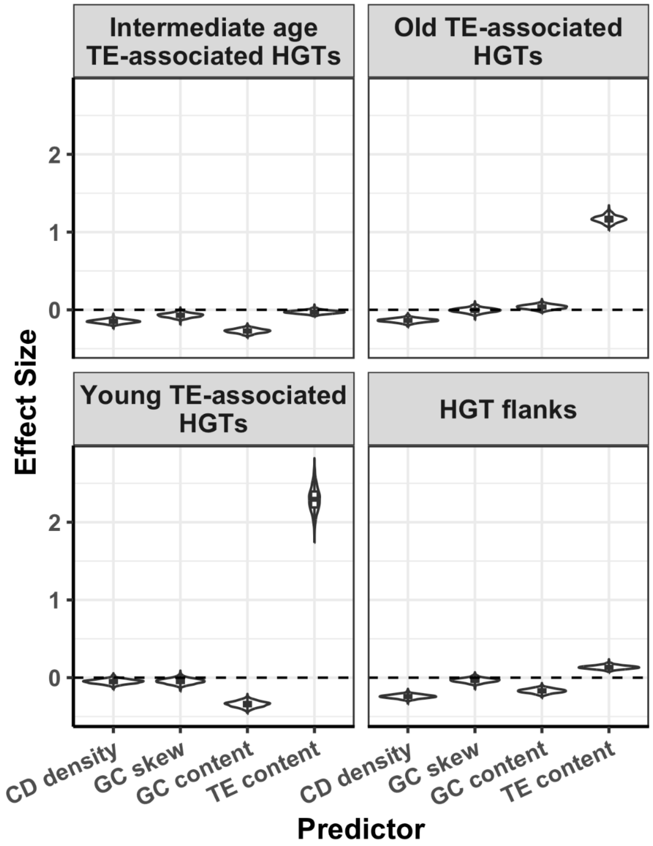


Supplementary Figure 44. Violin plots with embedded box plots summarizing the distributions of predictor effect sizes (regression coefficient values) across 500 chromosome-matched multinomial regression analyses. Each model had five predictors: coding sequence (CD) density, GC skew, GC content, and TE content. The models predicted five response classes: TE-associated HGT cores based on TE age as young, intermediate, and old, HGT flanking windows, and chromosome-matched background windows as the reference class. Positive coefficients indicate increased log odds of assignment to that response class for windows with higher values of the corresponding predictor, whereas negative coefficients indicate decreased log odds, relative to background. The horizontal dashed lines denote a regression coefficient of zero, indicating no association between the predictor and the response class.

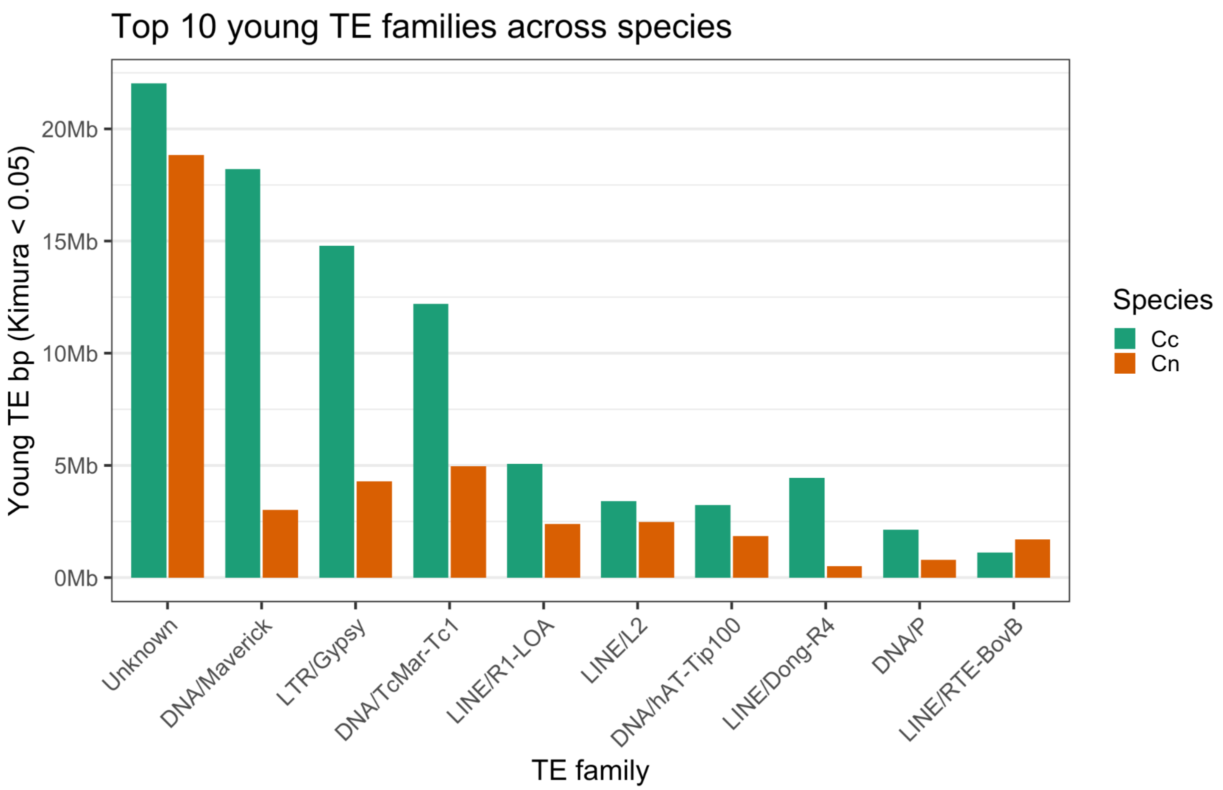


Supplementary Figure 45. Top 10 young (Kimura *D* < 0.05) node-specific TE families in *C. caryae* (Cc, teal) and *C. nanulus* (Cn, orange).

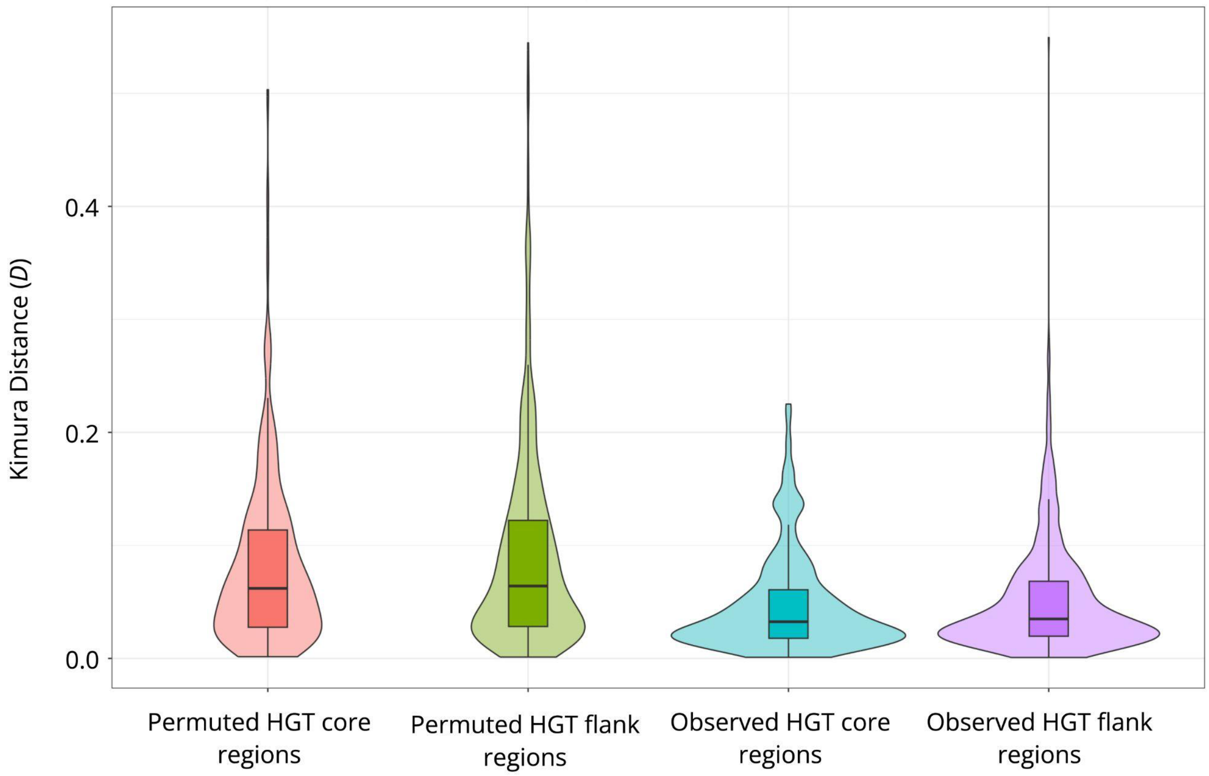


Supplementary Figure 46. Violin plots with embedded box plots summarizing the distribution of Maverick ages (Kimura *D*) across permuted HGT core, permuted HGT flank, observed HGT core, and observed HGT flank regions.

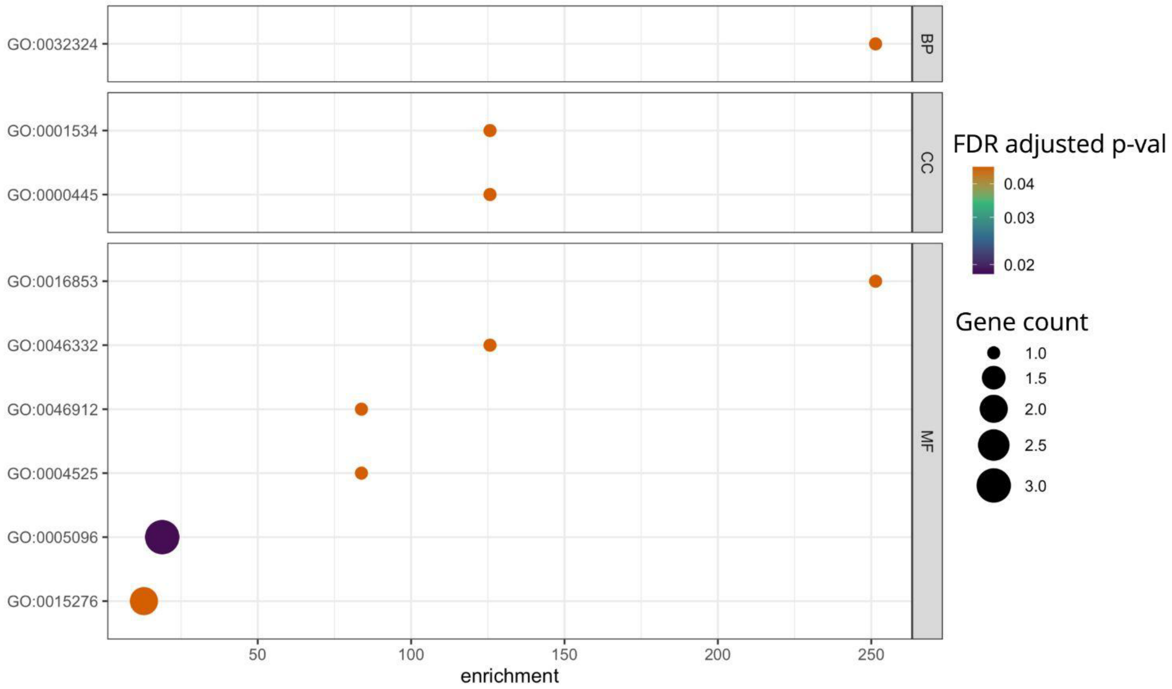


Supplemental Figure 47. Enriched Gene Ontology (GO) terms associated with HGTs in *C. caryae* by category: BP = biological process, CC = cellular component, and MF = molecular function. Gradient coloration represents FDR-adjusted *p*-values, and gene count indicates the number of genes associated with each enriched term.

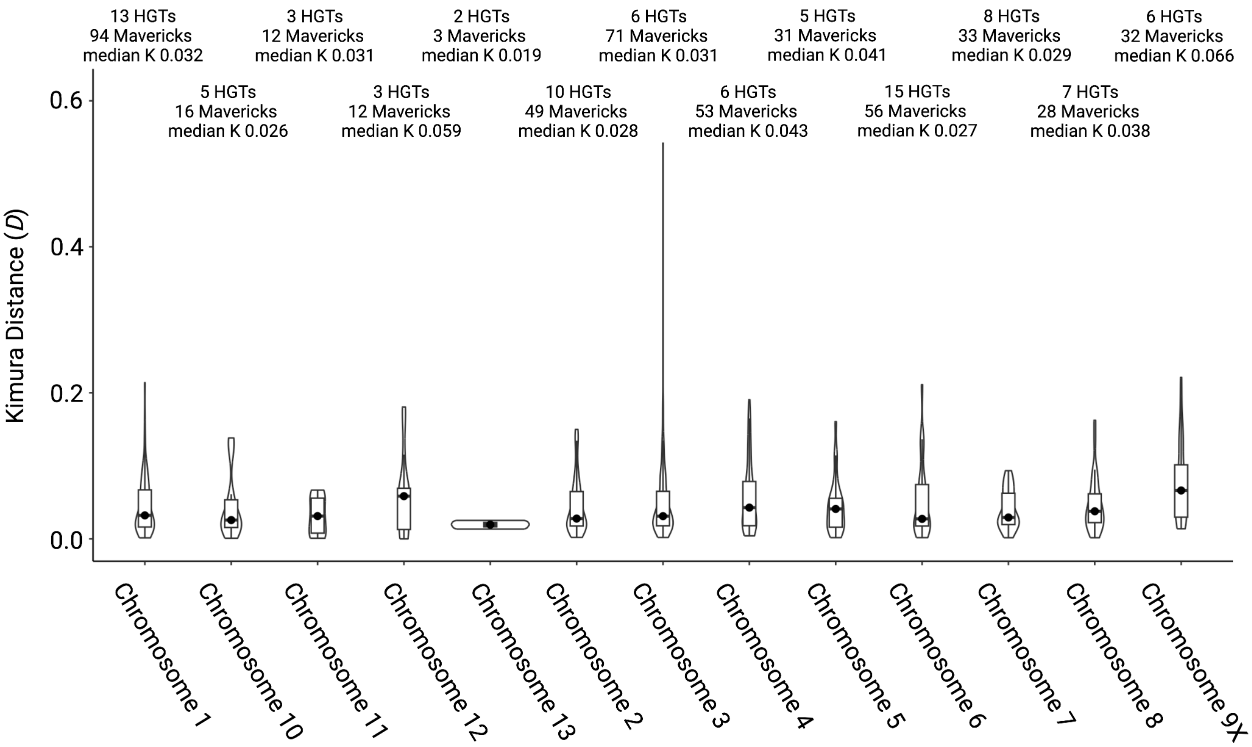


Supplemental Figure 48. Violin plots with embedded box plots summarizing the distribution of Kimura *D* estimates for Mavericks encapsulating HGTs in *C. caryae*. Chromosome 9 corresponds to the X chromosome.

Supplementary Table 1. Counts and cumulative sizes of structural variants (SVs), TEs and repeats, and HGT regions across homologous chromosomes in *Curculio* species: *C. caryae* (Cc), *C. nanulus* (Cn), and *C. glandium* (Cg).

| Homologous chromosome | Species-specific SVs >100 bp (size in Mb) | TEs and repeats (size in Mb) | HGT regions (size in kb) |
|---|---|---|---|
| Cc: 1 | 21,047 (263.09) | 355,659 (237.79) | 235 (928.06) |
| Cn: 1 | 23,759 (143.80) | 247,606 (133.89) | 26 (87.353) |
| Cg: 2 | 2,233 (137.69) | 225,385 (96.01) | 18 (58.05) |
| Cc: 2 | 37,472 (223.79) | 355,564 (225.38) | 75 (201.28) |
| Cn: 2 | 22,632 (142.24) | 251,738 (132.40) | 44 (129.97) |
| Cg: 1 | 2,208 (149.66) | 250,654 (107.15) | 12 (32.10) |
| Cc: 3 | 34,856 (205.48) | 320,544 (203.06) | 213 (932.97) |
| Cn: 3 | 19,672 (112.92) | 201,874 (104.25) | 32 (93.78) |
| Cg: 3 | 2,131 (129.61) | 214,146 (89.96) | 10 (27.05) |
| Cc: 4 | 21,531 (180.09) | 246,615 (156.21) | 51 (140.09) |
| Cn: 4 | 18,736 (108.14) | 194,417 (100.35) | 29 (86.44) |
| Cg: 4 | 1,669 (101.45) | 166,216 (70.39) | 8 (22.01) |
| Cc: 5 | 20,885 (155.91) | 238,793 (144.07) | 30 (89.65) |
| Cn: 5 | 18,354 (104.93) | 189,479 (97.58) | 29 (87.44) |

| | | | |
|---|---|---|---|
| Cg: 5 | 1,799 (97.84) | 158,256 (66.13) | 15 (45.93) |
| Cc: 7 | 18,023 (127.05) | 193,038 (114.34) | 31 (87.39) |
| Cn: 7 | 15,910 (80.01) | 147,303 (73.86) | 20 (61.99) |
| Cg: 6 | 1,645 (85.93) | 137,784 (58.90) | 5 (16.19) |
| Cc: 6 | 16,665 (135.95) | 192,167 (119.76) | 56 (158.88) |
| Cn: 6 | 15,714 (102.89) | 166,981 (87.99) | 28 (84.54) |
| Cg: 7 | 1,716 (76.8) | 113,913 (49.42) | 14 (36.36) |
| Cc: 10 | 14,726 (85.21) | 140,387 (73.36) | 14 (40.68) |
| Cn: 8 | 15,224 (80.56) | 125,292 (62.22) | 15 (41.63) |
| Cg: 8 | 1,291 (70.47) | 111,304 (46.46) | 13 (31.77) |
| Cc: 8 | 14,450 (105.92) | 162,234 (93.20) | 27 (79.73) |
| Cn: 9 | 13,904 (69.67) | 85,170 (42.94) | 15 (46.37) |
| Cg: 9 | 1,424 (67.65) | 104,771 (43.83) | 9 (22.56) |
| Cc: 12 | 11,293 (67.98) | 109,530 (56.71) | 12 (32.79) |
| Cn: 10 | 11,824 (55.65) | 140,140 (70.45) | 14 (43.92) |
| Cg: 10 | 905 (45.32) | 67,686 (28.04) | 11 (29.99) |
| Cc: 13 | 6,741 (48.56) | 72,574 (37.91) | 4 (8.48) |
| Cn: 13 | 7,463 (44.075) | 70,424 (32.34) | 10 (29.76) |
| Cg: 11 | 727 (40.68) | 54,183 (26.72) | 3 (9.35) |

| | | | |
|---|---|---|---|
| Cc: 11 | 8,998 (76.39) | 111,643 (65.12) | 12 (33.22) |
| Cn: 12 | 8,949 (52.86) | 100,271 (47.09) | 17 (57.18) |
| Cg: 12 | 628 (34.46) | 41,986 (20.14) | 10 (27.67) |
| Cc: 9 (X chromosome) | 14,276 (97.82) | 131,038 (93.20) | 30 (75.68) |
| Cn: 11 (X chromosome) | 11,198 (44.26) | 85,119 (44.32) | 4 (11.54) |
| Cg: 13 (X chromosome) | 2,217 (42.29) | 58,444 (24.89) | 8 (18.46) |

Supplementary Table 2. *Ccb* HGT annotations for Chromosomes 1 (C1) and 3 (C3). Bolded rows denote genes with significant branch-site likelihood ratio tests for positive selection.

| Gene ID | Protein annotation | C1 fragments | C3 fragments | C1 coordinates | C3 coordinates |
|---|---|---|---|---|---|
| **WP_395479791.1** | **ATP-dependent chaperone ClpB** | **1** | **1** | **134011286-134013155** | **136812448-136814956** |
| WP_395479792.1 | transcription antiterminator/RNA stability regulator CspE | 1 | 1 | 134205796-134206005 | 136615288-136615488 |
| **WP_395480257.1** | **ATP-dependent zinc metalloprotease FtsH** | **1** | **1** | **134041437-134043351** | **136841934-136843849** |
| WP_395480164.1 | type I glyceraldehyde-3-phosphate dehydrogenase | 1 | 1 | 134532379-134533350 | 137265139-137266109 |
| WP_395479678.1 | chaperonin GroEL | 1 | 2 | 134156614-134158268 | 137177374-137179012; 137221463-137222914 |
| **WP_395479659.1** | **isoleucine--tRNA ligase** | **1** | **1** | **134001521-134004334** | **136804177-136807004** |
| WP_395479776.1 | translation initiation factor IF-2 | 1 | 1 | 134033583-134036180 | 136836113-136837077 |
| **WP_395479794.1** | **leucine--tRNA ligase** | **1** | **1** | **134201489-134204062** | **136617191-136619771** |
| WP_395479804.1 | lipoyl synthase | 1 | 2 | 134191493-134192395 | 136514630-136515532; 136628929-136629831 |
| WP_395479660.1 | signal peptidase II | 1 | 1 | 134004334-134004799 | 136807004-136807469 |
| WP_395479801.1 | peptidoglycan DD-transpeptidase MrdA | 1 | 1 | 134196682-[\n]134198534 | 136622725-136624567 |
| WP_395480236.1 | NADH-quinone oxidoreductase subunit NuoG | 1 | 1 | 134515014-134517720 | 137247472-137250161 |
| WP_395479860.1 | two-component system response regulator PhoP | 1 | 2 | 134246882-134247452 | 136523760-136524425; 136572776-136573441 |
| WP_395479690.1 | pyruvate kinase | 1 | 2 | 134136677-134138116 | 137198008-137199435; 137201040-137202467 |

| | | | | | |
|---|---|---|---|---|---|
| **WP_395479665.1** | **transcription termination factor Rho** | **1** | **1** | **134220083-134221345** | **136597410-136598672** |
| WP_395479658.1 | bifunctional riboflavin kinase/FAD synthetase | 1 | 1 | 134000742-134001468 | 136803206-136804124 |
| WP_395480165.1 | signal peptide peptidase SppA | 1 | 1 | 134529506-134531177 | 137262271-137263938 |
| WP_395480266.1 | translational GTPase TypA | 1 | 1 | 134283263-134285056 | 137224800-137226590 |
| WP_395479685.1 | 2-octaprenyl-6-methoxyphenyl hydroxylase | 1 | 2 | 134144156-134145323 | 137190796-137191961; 137208508-137209676 |
| WP_395479688.1 | glucose-6-phosphate dehydrogenase | 1 | 2 | 134139312-134140447 | 137195331-137196774; 137203701-137205144 |
| **WP_395479664.1** | **proline--tRNA ligase** | **1** | **1** | **134214784-134216504** | **136602250-136603991** |
| WP_395479677.1 | co-chaperone GroES | 1 | 1 | 134158315-134158608 | 137177034-137177327 |
| **WP_395479781.1** | **DEAD/DEAH family ATP-dependent RNA helicase** | **1** | **1** | **134027062-134028334** | **136827634-136829330** |
| WP_395479784.1 | 3-deoxy-7-phosphoheptulonate synthase | 1 | 1 | 134017859-134018869 | 136819167-136819807 |
| WP_395479763.1 | acetyl-CoA carboxylase biotin carboxylase subunit | | 1 | | 136849019-136849673 |
| WP_395480224.1 | pyruvate dehydrogenase (acetyl-transferring), homodimeric type | | 1 | | 136703291-136705035 |
| WP_395479942.1 | arginine--tRNA ligase | 1 | | 134321652-134323387 | |
| WP_395480018.1 | shikimate kinase AroK | | 1 | | 136692605-136693119 |
| WP_395479814.1 | archaetidylserine decarboxylase | 1 | | 134180320-134181189 | |
| WP_395480232.1 | cytidine deaminase | | 1 | | 137318499-137319314 |
| WP_395479875.1 | ATP-dependent Clp protease ATP-binding subunit ClpX | 1 | | 134308438-134309712 | |
| WP_395480167.1 | cardiolipin synthase | | 1 | | 137256671-137257852 |

| | | | | | |
|---|---|---|---|---|---|
| ACICP0_RS01840 | (d)CMP kinase | 1 | | 134404298-134404935 | |
| WP_395480219.1 | magnesium/cobalt transporter CorA | 1 | | 134434714-134435659 | |
| WP_395479880.1 | ubiquinol oxidase subunit II | 1 | | 134297338-134298157 | |
| WP_395479881.1 | cytochrome o ubiquinol oxidase subunit I | 1 | | 134295226-134297220 | |
| WP_395480211.1 | ribosome biogenesis GTPase Der | 1 | | 134448976-134449738 | |
| WP_395479918.1 | molecular chaperone DnaK | 2 | | 134253849-134255290; 271083270-271083644 | |
| WP_395480154.1 | dipeptide/tripeptide permease DtpA | | 1 | | 137329703-137331149 |
| WP_395479887.1 | 1-deoxy-D-xylulose-5-phosphate synthase | | 2 | | 137234124-137235917; 137227594-137229398 |
| WP_395479830.1 | phosphopyruvate hydratase | | 1 | | 137155687-137156966 |
| WP_395479636.1 | 3-hydroxyacyl-ACP dehydratase FabZ | 1 | | 134371636-134372091 | |
| ACICP0_RS03070 | cell division protein FtsW | | 1 | | 136560104-136561200 |
| WP_395479870.1 | cell division protein FtsZ | | 2 | | 136554166-136554764; 136638432-136639042 |
| WP_395480233.1 | class II fumarate hydratase | | 1 | | 137338087-137339433 |
| WP_395479771.1 | phosphoglucosamine mutase | 1 | | 134039140-134040456 | |
| WP_395479890.1 | glutamine--fructose-6-phosphate transaminase (isomerizing) | 1 | | 134272349-134274188 | |
| WP_395479984.1 | glycine--tRNA ligase subunit alpha | 1 | | 134344071-134344975 | |
| WP_395479985.1 | glycine--tRNA ligase subunit beta | 1 | | 134341993-134344053 | |
| WP_395479873.1 | xanthine phosphoribosyltransferase | 1 | | 134314626-134315087 | |
| WP_395479600.1 | glutamate--cysteine ligase | 1 | | 134416623-134417111 | |

| | | | | | |
|---|---|---|---|---|---|
| WP_395479732.1 | protease modulator HflC | 1 | | 134067855-134068853 | |
| WP_395480163.1 | histone-like nucleoid-structuring protein H-NS | | 1 | | 137267742-137268142 |
| WP_395479808.1 | molecular chaperone HtpG | 2 | | 134185083-134186956; 134189638-134189741 | |
| WP_395479616.1 | integration host factor subunit beta | 1 | | 134402094-134402377 | |
| WP_395480226.1 | translation initiation factor IF-3 | | 1 | | 136679537-136680084 |
| WP_395479630.1 | 1-deoxy-D-xylulose-5-phosphate reductoisomerase | 1 | | 134379728-134380923 | |
| WP_395480141.1 | 4-(cytidine 5'-diphospho)-2-C-methyl-D-erythritol kinase | | 1 | | 137303168-137304015 |
| WP_395480214.1 | flavodoxin-dependent (E)-4-hydroxy-3-methylbut-2-enyl-diphosphate synthase | 1 | | 134443396-134444501 | |
| WP_395480245.1 | (2E,6E)-farnesyl-diphosphate-specific ditrans,polycis-undecaprenyl-diphosphate synthase | 1 | | 134378806-134379533 | |
| WP_395479909.1 | 3-deoxy-8-phosphooctulonate synthase | | 1 | | 136669688-136670530 |
| WP_395479874.1 | endopeptidase La | 1 | | 134309917-134312242 | |
| WP_395480225.1 | dihydrolipoyl dehydrogenase | | 1 | | 136700382-136701816 |
| WP_395479638.1 | lipid-A-disaccharide synthase | 1 | | 134369696-134370818 | |
| WP_395479871.1 | UDP-3-O-acyl-N-acetylglucosamine deacetylase | | 2 | | 136553153-136554051; 136639157-136640055 |
| WP_395479722.1 | cystathionine beta-lyase | 1 | | 134082788-134083960 | |
| WP_395480008.1 | methionine adenosyltransferase | | 1 | | 136709752-136710876 |

| | | | | | |
|---|---|---|---|---|---|
| WP_395480134.1 | septum site-determining protein MinD | | 1 | | 137281853-137282665 |
| WP_395479714.1 | membrane-bound lytic murein transglycosylase MltC | 1 | | 134093035-134094096 | |
| WP_395479858.1 | tRNA 2-thiouridine(34) synthase MnmA | 1 | | 134243406-134244529 | |
| WP_395479759.1 | rod shape-determining protein MreC | 1 | | 134063160-134063981 | |
| WP_395479695.1 | UDP-N-acetylglucosamine 1-carboxyvinyltransferase | 1 | | 134128210-134129446 | |
| WP_395479868.1 | UDP-N-acetylmuramate--L-alanine ligase | | 2 | | 136557379-136558377; 136634831-136635829 |
| WP_395480261.1 | UDP-N-acetylmuramoyl-L-alanyl-D-glutamate--2,6-diaminopimelate ligase | | 1 | | 136565065-136566435 |
| WP_395479717.1 | A/G-specific adenine glycosylase | 1 | | 134091633-134092462 | |
| WP_395480176.1 | NADH-quinone oxidoreductase subunit A | 1 | | 134510056-134510324 | |
| WP_395479774.1 | transcription termination factor NusA | | 1 | | 136837121-136838586 |
| WP_395479623.1 | 2-oxoglutarate dehydrogenase complex dihydrolipoyllysine-residue succinyltransferase | 1 | | 134386914-134388104 | |
| WP_395479812.1 | oligoribonuclease | 1 | | 134182206-134182742 | |
| WP_395479906.1 | leucyl aminopeptidase | | 1 | | 136677432-136678358 |
| WP_395480016.1 | glucose-6-phosphate isomerase | | 1 | | 136695734-136697346 |
| WP_395479617.1 | 6-phosphogluconolactonase | 1 | | 134397434-134398412 | |

| | | | | | |
|---|---|---|---|---|---|
| WP_395479861.1 | two-component system sensor histidine kinase PhoQ | | 2 | | 136571316-136572764; 136524437-136525882 |
| WP_395480240.1 | metalloprotease PmbA | 1 | | 134450640-134451977 | |
| WP_395479656.1 | nicotinate phosphoribosyltransferase | | 1 | | 136798399-136799600 |
| WP_395480143.1 | aminoacyl-tRNA hydrolase | | 1 | | 137305868-137306432 |
| WP_395479832.1 | glutamine hydrolyzing CTP synthase | | 1 | | 137156994-137158623 |
| WP_395479599.1 | recombinase RecA | 1 | | 134421914-134422906 | |
| WP_395480179.1 | exodeoxyribonuclease V subunit beta | 1 | | 134497423-134499278 | |
| WP_395480123.1 | lipopolysaccharide heptosyltransferase RfaC | | 1 | | 137375170-137376093 |
| WP_395480002.1 | 50S ribosomal protein L1 | | 1 | | 136722923-136723607 |
| WP_395480001.1 | 50S ribosomal protein L10 | | 1 | | 136723950-136724202 |
| WP_395480000.1 | 50S ribosomal protein L7/L12 | | 1 | | 136724452-136724824 |
| WP_395480190.1 | 50S ribosomal protein L16 | 1 | | 134484908-134485310 | |
| WP_395479999.1 | DNA-directed RNA polymerase subunit beta | | 2 | | 136725154-136727774; 136727782-136729112 |
| WP_395479998.1 | DNA-directed RNA polymerase subunit beta' | | 1 | | 136729205-136731079 |
| WP_395479940.1 | RNA pyrophosphohydrolase | | 1 | | 136543213-136543701 |
| WP_395479615.1 | 30S ribosomal protein S1 | 1 | | 134402458-134404144 | |
| WP_395480200.1 | 30S ribosomal protein S5 | 1 | | 134462941-134463428 | |
| WP_395480197.1 | 30S ribosomal protein S8 | 1 | | 134482306-134482698 | |
| WP_395480206.1 | 30S ribosomal protein S11 | 1 | | 134459877-134460261 | |

| | | | | | |
|---|---|---|---|---|---|
| WP_395479963.1 | 16S rRNA (guanine(966)-N(2))-methyltransferase RsmD | | 1 | | 136748028-136748551 |
| WP_395479620.1 | succinate dehydrogenase flavoprotein subunit | 1 | | 134391905-134393672 | |
| ACICP0_RS03110 | preprotein translocase subunit SecA | | 2 | | 136549490-136551997; 136641211-136643722 |
| WP_395480203.1 | preprotein translocase subunit SecY | 1 | | 134460961-134462270 | |
| WP_395479634.1 | molecular chaperone Skp | 1 | | 134373247-134373744 | |
| WP_395479959.1 | transcriptional regulator SlyA | 1 | | 134330553-134330980 | |
| WP_395479724.1 | spermidine N1-acetyltransferase | 1 | | 134077399-134077937 | |
| WP_395480221.1 | Fe-S cluster assembly protein SufB | | 1 | | 136751499-136752985 |
| WP_395479640.1 | tRNA adenosine(34) deaminase TadA | 1 | | 134364325-134364642 | |
| WP_395479856.1 | thiamine/thiamine pyrophosphate ABC transporter permease | 1 | | 134238793-134240350 | |
| WP_395480026.1 | threonine--tRNA ligase | | 2 | | 136680660-136681890; 136680090-136680633 |
| WP_395480015.1 | metalloprotease TldD | | 1 | | 136698690-136700122 |
| WP_395480138.1 | type I DNA topoisomerase | | 1 | | 137289396-137291967 |
| WP_395479650.1 | elongation factor Tu | | 1 | | 136771958-136773080 |
| WP_395479935.1 | 4-hydroxy-3-polyprenylbenzoate decarboxylase | | 1 | | 136533483-136534953 |
| WP_395479718.1 | uracil phosphoribosyltransferase | 1 | | 134089785-134090410 | |
| WP_395480122.1 | lipid IV(A) 3-deoxy-D-manno-octulosonic acid transferase | | 1 | | 137384137-137385348 |
| WP_395480259.1 | tol-pal system protein YbgF | | 1 | | 137150885-137151306 |

| | | | | | |
|---|---|---|---|---|---|
| WP_395479922.1 | membrane protein insertase YidC | 1 | | 134248610-134250247 | |
| WP_395479923.1 | membrane protein insertion efficiency factor YidD | 1 | | 134248360-134248597 | |
| WP_395480242.1 | fructose-1-phosphate/6-phosphogluconate phosphatase | 1 | | 134417204-134417763 | |
| WP_395479621.1 | succinate dehydrogenase iron-sulfur subunit | 1 | | 134391172-134391884 | |
| WP_395479709.1 | MBL fold metallo-hydrolase | 1 | | 134115558-134116146 | |
| WP_395479723.1 | single-stranded DNA-binding protein | 1 | | 134079177-134079696 | |
| WP_395479760.1 | rod shape-determining protein | 1 | | 134062011-134063050 | |
| WP_395479846.1 | metal ABC transporter permease | 1 | | 134178515-134179307 | |
| WP_395480022.1 | YchF/TatD family DNA exonuclease | | 1 | | 136687410-136688080 |
| WP_395480142.1 | ribose-phosphate pyrophosphokinase | | 1 | | 137304282-137305231 |
| WP_395480158.1 | 2-hydroxyacid dehydrogenase | | 1 | | 137342741-137343736 |
| WP_395480207.1 | DNA-directed RNA polymerase subunit alpha | 1 | | 134458242-134459201 | |
| ACICP0_RS03380 | DNA polymerase III subunit chi | | 1 | | 136676868-136677295 |

Supplementary Table 3. Pairwise Jukes-Cantor genetic distances estimated across four *Ccb*-derived homologous regions: the Chromosome 1 region, the upstream Chromosome 3 region (A), the downstream Chromosome 3 region (B), the extrachromosomal contig (JAKZMK010000236.1), and the *Ccb* reference genome.

| | **Chromosome 1** | **Chromosome 3A** | **Chromosome 3B** | ***Ccb* genome** | **JAKZMK010000236.1** |
|---|---|---|---|---|---|
| **Chromosome 1** | - | 0.0277 | 0.0560 | 0.222 | 0.00162 |
| **Chromosome 3A** | | - | 0.110 | 0.217 | 0.0143 |
| **Chromosome 3B** | | | - | 0.229 | 0.0253 |
| ***Ccb* genome** | | | | - | 0.214 |

Supplementary Table 4. Genome-wide repeat family composition in *C. caryae.* Counts and cumulative sizes are shown for each TE family in regions classified as old or young based on the Kimura *D* thresholds described in *Methods*.

| TE family | Counts of old TEs | Counts of young TEs | Cumulative size of old TEs (bp) | Cumulative size of young TEs (bp) | Proportion of young TE counts (col2/[col1+2) | Proportion of young TE size col4/(col3+4) |
|---|---|---|---|---|---|---|
| Unknown | 973,281 | 297,975 | 368,832,962 | 236,460,405 | 0.234 | 0.391 |
| DNA/Maverick | 141,401 | 45,094 | 103,679,464 | 113,497,558 | 0.242 | 0.523 |
| LTR/Gypsy | 102,346 | 43,780 | 73,303,501 | 102,943,396 | 0.300 | 0.584 |
| DNA/TcMar-Tc1 | 110,048 | 43,108 | 34,797,779 | 63,913,645 | 0.281 | 0.647 |
| LINE/R1-LOA | 99,143 | 26,974 | 53,294,578 | 32,231,503 | 0.214 | 0.377 |
| LINE/Dong-R4 | 28,058 | 14,077 | 20,127,356 | 31,967,322 | 0.334 | 0.614 |
| LINE/L2 | 49,374 | 14,496 | 18,841,058 | 21,892,504 | 0.227 | 0.537 |
| DNA/hAT-Tip100 | 38,478 | 16,050 | 14,290,869 | 20,424,795 | 0.294 | 0.588 |
| DNA/P | 26,179 | 10,282 | 7,917,624 | 11,940,737 | 0.282 | 0.601 |

| | | | | | | |
|---|---|---|---|---|---|---|
| LINE/I | 8,261 | 3,554 | 5,440,727 | 10,762,498 | 0.301 | 0.664 |
| LINE/Penelope | 25,131 | 11,384 | 7,916,460 | 10,210,735 | 0.312 | 0.563 |
| LINE/I-Jockey | 60,745 | 6,476 | 31,856,392 | 8,741,410 | 0.096 | 0.215 |
| LTR/Pao | 14,986 | 3,833 | 7,358,761 | 8,151,393 | 0.204 | 0.526 |
| LINE/CR1 | 18,860 | 6,495 | 8,024,478 | 7,571,152 | 0.256 | 0.485 |
| LINE/RTE-BovB | 49,019 | 5,938 | 23,477,268 | 6,994,769 | 0.108 | 0.230 |
| DNA/TcMar-m44 | 10,357 | 6,712 | 3,106,917 | 6,675,792 | 0.393 | 0.682 |
| DNA/Academ-1 | 31,907 | 6,620 | 11,319,228 | 6,241,069 | 0.172 | 0.355 |
| DNA/Crypton-I | 12,571 | 6,397 | 3,205,642 | 5,933,605 | 0.337 | 0.649 |
| LTR/Copia | 7,146 | 2,796 | 3,639,347 | 5,702,881 | 0.281 | 0.610 |
| DNA/hAT-Ac | 13,572 | 5,356 | 3,225,804 | 5,427,226 | 0.283 | 0.627 |
| DNA/CMC-Chapaev-3 | 7,419 | 3,830 | 2,170,790 | 3,778,964 | 0.340 | 0.635 |
| DNA/hAT-Blackjack | 4,003 | 2,560 | 908,160 | 3,623,481 | 0.390 | 0.800 |
| DNA/CMC-Transib | 13,861 | 2,256 | 3,856,709 | 3,429,823 | 0.140 | 0.471 |
| DNA/hAT-Tag1 | 3,326 | 2,381 | 902,899 | 3,118,653 | 0.417 | 0.775 |
| LTR/DIRS | 2,705 | 1,330 | 1,414,810 | 3,112,689 | 0.330 | 0.688 |
| LINE | 8,379 | 3,616 | 2,482,089 | 3,042,868 | 0.301 | 0.551 |
| DNA | 13,209 | 2,767 | 2,825,787 | 2,912,762 | 0.173 | 0.508 |
| DNA/MULE-MuDR | 5,001 | 2,060 | 1,316,320 | 2,662,063 | 0.292 | 0.669 |
| RC/Helitron | 3,222 | 2,012 | 682,616 | 1,805,910 | 0.384 | 0.726 |
| LINE/R1 | 9,841 | 992 | 4,927,087 | 1,596,994 | 0.092 | 0.245 |
| DNA/PiggyBac | 10,682 | 1,934 | 3,003,633 | 1,580,541 | 0.153 | 0.345 |
| DNA/Crypton-V | 9,767 | 812 | 2,116,807 | 1,523,719 | 0.077 | 0.419 |
| DNA/PIF-ISL2EU | 4,853 | 924 | 1,203,164 | 1,203,482 | 0.160 | 0.500 |
| DNA/TcMar-Mariner | 29,155 | 1,395 | 6,204,198 | 1,155,689 | 0.046 | 0.157 |
| DNA/Crypton-A | 3,202 | 1,898 | 797,748 | 1,118,009 | 0.372 | 0.584 |

| | | | | | | |
|---|---|---|---|---|---|---|
| DNA/hAT-hATm | 3,311 | 770 | 1,113,751 | 1,097,921 | 0.189 | 0.496 |
| DNA/PIF-Harbinger | 11,632 | 1,137 | 3,247,565 | 1,028,550 | 0.089 | 0.241 |
| DNA/TcMar-Fot1 | 7,340 | 588 | 2,977,426 | 890,098 | 0.074 | 0.230 |
| DNA/Sola-1 | 2,029 | 422 | 598,101 | 637,317 | 0.172 | 0.516 |
| DNA/MULE-NOF | 18 | 162 | 8,959 | 577,712 | 0.900 | 0.985 |
| DNA/hAT-Charlie | 7,059 | 1,223 | 1,504,390 | 569,520 | 0.148 | 0.275 |
| DNA/Merlin | 3,943 | 761 | 2,114,116 | 543,556 | 0.162 | 0.205 |
| DNA/CMC-EnSpm | 1,662 | 216 | 208,814 | 542,776 | 0.115 | 0.722 |
| DNA/TcMar-Pogo | 3,229 | 542 | 905,917 | 463,905 | 0.144 | 0.339 |
| LINE/RTE | 2,423 | 576 | 426,344 | 427,680 | 0.192 | 0.501 |
| LINE/RTE-RTE | 5,417 | 333 | 3,131,003 | 416,433 | 0.058 | 0.117 |
| DNA/Sola-2 | 71 | 189 | 42,950 | 412,220 | 0.727 | 0.906 |
| DNA/Ginger-2 | 1,731 | 494 | 445,313 | 342,540 | 0.222 | 0.435 |
| DNA/TcMar-Tigger | 4,146 | 263 | 1,169,562 | 311,588 | 0.060 | 0.210 |
| SINE/5S-Deu-L2 | 4,230 | 1,451 | 891,936 | 299,471 | 0.255 | 0.251 |
| DNA/hAT-hAT19 | 314 | 300 | 96,011 | 289,610 | 0.489 | 0.751 |
| LINE/CRE | 956 | 154 | 372,149 | 154,666 | 0.139 | 0.294 |
| DNA/hAT-hobo | 258 | 324 | 76,252 | 135,410 | 0.557 | 0.640 |
| DNA/Kolobok-E | 746 | 187 | 81,816 | 109,145 | 0.200 | 0.572 |
| DNA/TcMar-ISRm11 | 74 | 54 | 38,270 | 38,360 | 0.422 | 0.501 |
| DNA/P-Fungi | 308 | 28 | 60,178 | 28,566 | 0.083 | 0.322 |
| LINE/R2 | 135 | 3 | 81,543 | 7,827 | 0.022 | 0.088 |
| DNA/hAT-hATx | 288 | 10 | 124,293 | 1,467 | 0.034 | 0.012 |

Supplementary Table 5. Genome-wide repeat family composition in *C. nanulus.* Counts and cumulative sizes are shown for each TE family in regions classified as old or young based on the Kimura *D* thresholds described in *Methods*.

| TE family | Counts of old TEs | Counts of young TEs | Cumulative size of old TEs (bp) | Cumulative size of young TEs (bp) | Proportion of young TE counts (col2/[col1+2) | Proportion of young TE size col4/(col3+4) |
|---|---|---|---|---|---|---|
| Unknown | 860198 | 260337 | 302511088 | 208,232,464 | 0.232 | 0.408 |
| DNA/TcMar-Tc1 | 80666 | 26423 | 23127520 | 31989071 | 0.247 | 0.58 |
| LTR/Gypsy | 44402 | 12243 | 24497727 | 31863409 | 0.216 | 0.565 |
| DNA/Maverick | 33164 | 7098 | 21049344 | 22138849 | 0.176 | 0.513 |
| LINE/R1-LOA | 76653 | 17104 | 40687808 | 17760158 | 0.182 | 0.304 |
| LINE/L2 | 32337 | 11171 | 13849309 | 17604366 | 0.257 | 0.56 |
| LINE/RTE-BovB | 61032 | 9083 | 33051603 | 13526336 | 0.13 | 0.29 |
| DNA/hAT-Tip100 | 36628 | 14475 | 12368538 | 13298514 | 0.283 | 0.518 |
| LTR/Pao | 20461 | 5576 | 9273342 | 10939533 | 0.214 | 0.541 |
| DNA/Academ-1 | 33921 | 7455 | 13040093 | 8988532 | 0.18 | 0.408 |
| LINE/Penelope | 20346 | 8566 | 6447707 | 6848810 | 0.296 | 0.515 |
| LINE/Dong-R4 | 3271 | 2460 | 2265263 | 5847783 | 0.429 | 0.721 |
| DNA/P | 19102 | 5079 | 4814945 | 5377613 | 0.21 | 0.528 |
| LINE/I-Jockey | 42426 | 3664 | 25562598 | 4947234 | 0.079 | 0.162 |
| LINE/CR1 | 10254 | 4057 | 3829837 | 4725926 | 0.283 | 0.552 |
| DNA/hAT-Ac | 12018 | 5323 | 2227163 | 4013086 | 0.307 | 0.643 |
| LINE/RTE-RTE | 10414 | 2644 | 5790173 | 3884001 | 0.202 | 0.401 |
| DNA/CMC-Transib | 9056 | 3801 | 3024239 | 3237049 | 0.296 | 0.517 |
| DNA/TcMar-m44 | 6211 | 3207 | 1788629 | 2905524 | 0.341 | 0.619 |
| LTR/Unknown | 10003 | 2602 | 5278446 | 2865878 | 0.206 | 0.352 |
| LINE/I | 9375 | 1935 | 5217129 | 2855489 | 0.171 | 0.354 |
| DNA/hAT-Tag1 | 4475 | 2519 | 1033830 | 2783305 | 0.36 | 0.729 |
| RC/Helitron | 4790 | 2064 | 1120022 | 2591361 | 0.301 | 0.698 |
| LTR/Copia | 2749 | 1718 | 1081923 | 2515427 | 0.385 | 0.699 |
| DNA/Crypton-I | 11594 | 2313 | 2666221 | 2007677 | 0.166 | 0.43 |
| DNA/hAT-Blackjack | 6069 | 2737 | 1111437 | 1665107 | 0.311 | 0.6 |
| DNA | 9460 | 1606 | 1740290 | 1325904 | 0.145 | 0.432 |

| | | | | | | |
|---|---|---|---|---|---|---|
| LINE/R1 | 10262 | 1130 | 5448623 | 1137964 | 0.099 | 0.173 |
| DNA/PIF-Harbinger | 9246 | 1267 | 2319490 | 1017153 | 0.121 | 0.305 |
| DNA/PiggyBac | 5812 | 819 | 1471122 | 983247 | 0.124 | 0.401 |
| DNA/MULE-MuDR | 6013 | 956 | 1522523 | 831278 | 0.137 | 0.353 |
| DNA/CMC-Chapaev-3 | 5359 | 975 | 1555815 | 796680 | 0.154 | 0.339 |
| DNA/TcMar-Mariner | 36584 | 1550 | 6938399 | 610651 | 0.041 | 0.081 |
| DNA/Sola-1 | 3330 | 513 | 769268 | 604814 | 0.133 | 0.44 |
| LTR/DIRS | 2998 | 388 | 1042305 | 551993 | 0.115 | 0.346 |
| DNA/PIF-ISL2EU | 3805 | 576 | 818300 | 544116 | 0.131 | 0.399 |
| DNA/hAT-hATm | 3146 | 670 | 940214 | 540416 | 0.176 | 0.365 |
| LINE | 1589 | 446 | 273277 | 491207 | 0.219 | 0.643 |
| DNA/TcMar-Fot1 | 6482 | 360 | 2362612 | 476519 | 0.053 | 0.168 |
| DNA/Crypton-V | 3456 | 591 | 690060 | 423050 | 0.146 | 0.38 |
| DNA/hAT-Charlie | 2149 | 368 | 563546 | 411466 | 0.146 | 0.422 |
| DNA/TcMar-Pogo | 3297 | 404 | 866928 | 342591 | 0.109 | 0.283 |
| LTR | 2358 | 228 | 699280 | 321452 | 0.088 | 0.315 |
| DNA/Sola-2 | 245 | 119 | 86740 | 250419 | 0.327 | 0.743 |
| DNA/Crypton-A | 1056 | 288 | 334374 | 249879 | 0.214 | 0.428 |
| LINE/RTE | 375 | 138 | 374971 | 226368 | 0.269 | 0.376 |
| DNA/hAT-hATx | 1170 | 291 | 359224 | 217521 | 0.199 | 0.377 |
| DNA/P-Fungi | 213 | 75 | 99828 | 207693 | 0.26 | 0.675 |
| DNA/TcMar-ISRm11 | 121 | 188 | 72152 | 193501 | 0.608 | 0.728 |
| DNA/Ginger-2 | 1881 | 290 | 587495 | 183350 | 0.134 | 0.238 |
| DNA/Kolobok-E | 363 | 94 | 86261 | 179904 | 0.206 | 0.676 |
| DNA/TcMar-Tigger | 4560 | 279 | 1021166 | 151625 | 0.058 | 0.129 |
| LINE/CRE | 830 | 110 | 356181 | 139651 | 0.117 | 0.282 |
| DNA/MULE-NOF | 33 | 141 | 5363 | 134550 | 0.81 | 0.962 |
| DNA/CMC-Chapaev | 234 | 55 | 85393 | 125985 | 0.19 | 0.596 |

| | | | | | | |
|---|---|---|---|---|---|---|
| DNA/hAT-hAT5 | 129 | 73 | 60617 | 118998 | 0.361 | 0.663 |
| DNA/PIF-Spy | 198 | 40 | 111411 | 107056 | 0.168 | 0.49 |
| SINE/5S-Deu-L2 | 2359 | 568 | 680975 | 85843 | 0.194 | 0.112 |
| DNA/hAT-hAT19 | 826 | 201 | 138464 | 85611 | 0.196 | 0.382 |
| DNA/CMC-EnSpm | 233 | 26 | 65423 | 54667 | 0.1 | 0.455 |
| DNA/hAT-hobo | 177 | 140 | 46377 | 52914 | 0.442 | 0.533 |
| DNA/Zator | 72 | 83 | 6888 | 43826 | 0.535 | 0.864 |
| DNA/Merlin | 77 | 44 | 21021 | 16412 | 0.364 | 0.438 |
| LINE/R2 | 309 | 7 | 165209 | 125 | 0.022 | 0.001 |
| LINE/CR1-Zenon | 233 | 0 | 25961 | 0 | 0.0 | 0.0 |

Supplementary Table 6. Genome-wide repeat family composition in *C. glandium.* Counts and cumulative sizes are shown for each TE family in regions classified as old or young based on the Kimura *D* thresholds described in *Methods*.

| TE family | Counts of old TEs | Counts of young TEs | Cumulative size of old TEs (bp) | Cumulative size of young TEs (bp) | Proportion of young TE counts (col2/[col1+2) | Proportion of young TE size col4/(col3+4) |
|---|---|---|---|---|---|---|
| Unknown | 743872 | 250051 | 191785851 | 151502164 | 0.252 | 0.441 |
| LTR/Gypsy | 45924 | 31407 | 22461590 | 55787330 | 0.406 | 0.713 |
| DNA/TcMar-Tc1 | 116649 | 33615 | 32025015 | 29768072 | 0.224 | 0.482 |
| LTR/Pao | 22403 | 11226 | 10771189 | 15084764 | 0.334 | 0.583 |
| LINE/R1-LOA | 44497 | 10659 | 13030921 | 14172632 | 0.193 | 0.521 |
| LINE/L2 | 36957 | 8537 | 15664468 | 10065055 | 0.188 | 0.391 |
| LINE/Penelope | 17216 | 10663 | 4852781 | 8059290 | 0.382 | 0.624 |
| DNA/Academ-1 | 25865 | 7488 | 6151110 | 7379495 | 0.225 | 0.545 |
| DNA/TcMar-m44 | 9500 | 7420 | 2301472 | 6814775 | 0.439 | 0.748 |
| DNA/Maverick | 8958 | 2322 | 3764669 | 6665791 | 0.206 | 0.639 |
| LINE/CR1 | 12905 | 4637 | 5246366 | 5881359 | 0.264 | 0.529 |
| DNA/hAT-Tip100 | 17266 | 8160 | 3691537 | 5501090 | 0.321 | 0.598 |
| LINE/Dong-R4 | 1249 | 2491 | 668075 | 4762971 | 0.666 | 0.877 |

| | | | | | | |
|---|---|---|---|---|---|---|
| DNA/Crypton-I | 10389 | 4863 | 2821320 | 4312149 | 0.319 | 0.604 |
| LINE/I | 8536 | 2592 | 4651643 | 4254887 | 0.233 | 0.478 |
| LINE/RTE-BovB | 20074 | 2433 | 7066956 | 3667257 | 0.108 | 0.342 |
| DNA/P | 10564 | 4183 | 2418784 | 3373657 | 0.284 | 0.582 |
| LTR/Unknown | 5695 | 4349 | 2208040 | 3140765 | 0.433 | 0.587 |
| LTR/Copia | 4420 | 1994 | 1671567 | 3035052 | 0.311 | 0.645 |
| LINE/RTE-RTE | 2496 | 1124 | 1636015 | 2061595 | 0.310 | 0.558 |
| DNA/hAT-hATm | 1592 | 1491 | 587392 | 1922984 | 0.484 | 0.766 |
| DNA | 5455 | 2678 | 1030447 | 1907902 | 0.329 | 0.649 |
| LINE/I-Jockey | 12222 | 1741 | 4421364 | 1742375 | 0.125 | 0.283 |
| DNA/Sola-1 | 5058 | 2447 | 1103116 | 1732967 | 0.326 | 0.611 |
| RC/Helitron | 1903 | 1892 | 339178 | 1572617 | 0.499 | 0.823 |
| DNA/CMC-Transib | 6354 | 1215 | 1482578 | 1239609 | 0.161 | 0.455 |
| DNA/CMC-EnSpm | 2192 | 1470 | 617442 | 1041833 | 0.401 | 0.628 |
| DNA/CMC-Chapaev-3 | 4327 | 909 | 1306944 | 1011536 | 0.174 | 0.436 |
| DNA/PIF-Harbinger | 4790 | 1150 | 1122901 | 965153 | 0.194 | 0.462 |
| DNA/PIF-ISL2EU | 3036 | 1470 | 386093 | 951258 | 0.326 | 0.711 |
| DNA/hAT-Ac | 2838 | 1336 | 502184 | 797476 | 0.320 | 0.614 |
| DNA/PiggyBac | 7941 | 1169 | 2657553 | 755198 | 0.128 | 0.221 |
| DNA/TcMar-Fot1 | 3916 | 725 | 1435248 | 717265 | 0.156 | 0.333 |
| LINE | 2200 | 1061 | 725017 | 629303 | 0.325 | 0.465 |
| DNA/Crypton-V | 4457 | 693 | 634542 | 584270 | 0.135 | 0.479 |
| DNA/hAT-Charlie | 625 | 394 | 121299 | 505776 | 0.387 | 0.807 |
| DNA/TcMar-Tigger | 758 | 400 | 125732 | 327220 | 0.345 | 0.722 |
| DNA/MULE-MuDR | 1578 | 486 | 320271 | 316670 | 0.235 | 0.497 |
| DNA/Crypton-A | 843 | 525 | 287581 | 290684 | 0.384 | 0.503 |
| DNA/TcMar-Mariner | 8400 | 410 | 1328198 | 290543 | 0.047 | 0.179 |
| LTR/DIRS | 1439 | 153 | 590006 | 234679 | 0.096 | 0.285 |

| | | | | | | |
|---|---|---|---|---|---|---|
| DNA/Sola-2 | 253 | 267 | 25990 | 231561 | 0.513 | 0.899 |
| DNA/PIF-Spy | 173 | 160 | 41185 | 209842 | 0.480 | 0.836 |
| DNA/hAT-Tag1 | 1467 | 378 | 341837 | 196726 | 0.205 | 0.365 |
| DNA/Ginger-1 | 50 | 332 | 3984 | 192706 | 0.869 | 0.980 |
| DNA/hAT-Blackjack | 6555 | 436 | 1577829 | 168779 | 0.062 | 0.097 |
| DNA/hAT-hAT19 | 1676 | 191 | 371910 | 166050 | 0.102 | 0.309 |
| LINE/R1 | 4644 | 119 | 1614508 | 139193 | 0.025 | 0.079 |
| DNA/Kolobok-E | 1870 | 346 | 435300 | 128692 | 0.156 | 0.228 |
| DNA/hAT-hATx | 702 | 173 | 108731 | 104353 | 0.198 | 0.490 |
| DNA/TcMar-Pogo | 475 | 137 | 133787 | 101343 | 0.224 | 0.431 |
| DNA/TcMar-ISRm11 | 468 | 94 | 158617 | 98868 | 0.167 | 0.384 |
| DNA/CMC-Chapaev | 158 | 76 | 27970 | 53379 | 0.325 | 0.656 |
| DNA/Ginger-2 | 406 | 15 | 114491 | 23142 | 0.036 | 0.168 |
| SINE/tRNA-V | 188 | 75 | 37941 | 15488 | 0.285 | 0.290 |
| LTR | 7 | 4 | 2604 | 3007 | 0.364 | 0.536 |
| LINE/R2 | 145 | 1 | 76491 | 2404 | 0.007 | 0.030 |
| LINE/RTE | 947 | 3 | 201869 | 93 | 0.003 | 0.000 |
| LINE/CRE | 245 | 1 | 46362 | 28 | 0.004 | 0.001 |
| LINE/RTE-X | 103 | 0 | 47729 | 0 | 0.000 | 0.000 |
